# FEMA-Long: Modeling unstructured covariances for discovery of time-dependent effects in large-scale longitudinal datasets

**DOI:** 10.1101/2025.05.09.653146

**Authors:** Pravesh Parekh, Nadine Parker, Diliana Pecheva, Evgeniia Frei, Marc Vaudel, Diana M. Smith, Alison Rigby, Piotr Jahołkowski, Ida Elken Sønderby, Viktoria Birkenæs, Nora Refsum Bakken, Chun Chieh Fan, Carolina Makowski, Jakub Kopal, Robert Loughnan, Donald J. Hagler, Dennis van der Meer, Stefan Johansson, Pål Rasmus Njølstad, Terry L. Jernigan, Wesley K. Thompson, Oleksandr Frei, Alexey A. Shadrin, Thomas E. Nichols, Ole A. Andreassen, Anders M. Dale

**Author notes:** Correspondence should be addressed to: Dr. Pravesh Parekh (PP), Prof. Anders M. Dale (AMD).

## Abstract

While linear mixed-effects (LME) models are common for analyzing longitudinal data, most users rely on random intercepts or simple stationary covariance, due to unavailability of computationally tractable solutions. Here, we extend the Fast and Efficient Mixed-Effects Algorithm (FEMA) and present FEMA-Long, a computationally tractable approach to flexibly modeling longitudinal covariance suitable for high-dimensional data. FEMA-Long can: i) model unstructured covariance, ii) model covariates as smooth functions using splines, iii) discover time-dependent effects of covariates with spline interactions, and iv) use these flexible longitudinal modeling strategies to perform longitudinal genome-wide association studies and discover time-dependent genetic effects, in a computationally scalable manner, suitable for high-dimensional data. Through extensive simulations, we show that estimates from FEMA-Long are accurate, while being up to several thousand times faster and with minimal carbon footprint. To show the utility of FEMA-Long for discovering novel biological signal, using data from the Norwegian Mother, Father and Child Cohort Study (MoBa), we performed a longitudinal genome-wide association study with non-linear SNP-by-time interaction on length, weight, and BMI of 68,273 infants with up to six measurements in the first year of life. We found dynamic patterns of random effects including time-varying heritability and genetic correlations, as well as several genetic variants showing time-dependent effects, highlighting the applicability of FEMA-Long to enable novel discoveries. FEMA-Long is available at: https://github.com/cmig-research-group/cmig_tools.

**Author summary:** Most large-scale datasets have complexities such as repeated measures, related individuals, or other dependencies across samples, preventing the use of standard regression approaches for analysis. In such circumstances, linear mixed-effects modeling is often employed. However, for high-dimensional datasets, fitting these models is quite challenging. Further, most standard uses of linear mixed-effects modeling focus on simpler covariance models, which may not hold. Here, we introduce FEMA-Long, a novel computationally efficient analytical framework for fitting linear mixed-effects models with time-varying random effects, as well as allowing the effect of the covariates to change smoothly over time by using splines. This is particularly relevant when, for example, studying the effect of genetic variants on phenotypes, where the effects could be non-linear over time. The FEMA-Long framework allows time-varying heritability as well as discovery of genetic variants that show time-dependent effects. By performing a genome-wide association study on data from the Norwegian Mother, Father and Child Cohort Study (MoBa) using FEMA-Long, we show the discovery of genetic variants with time-dependent effects on infant length, weight, and BMI during the first year of life. Our results highlight the potential of using FEMA-Long to make novel discoveries that can lead to biological insights on the genetics of complex traits as well as improve the potential of using genetics for personalized prediction.

## Introduction

Large-scale longitudinal datasets are critical for novel discoveries in health and medicine and can transform healthcare (1). Longitudinal data allows studying the dynamic processes underlying health and disease development/progression by modeling trajectories, examining time-dependent effects of variables, investigating association with health indicators and outcomes, and making predictions of outcomes. Examples of large-scale densely-sampled datasets include the UK Biobank (https://www.ukbiobank.ac.uk), All of Us (https://allofus.nih.gov), the Adolescent Brain Cognitive Development^℠^ Study (https://abcdstudy.org), and many others (2).

To leverage the full potential of these studies, we need computationally efficient modeling strategies, optimized for large-scale high-dimensional datasets. These strategies need to account for non-independence of observations (e.g., repeated measurements and relationships between individuals), other sources of variance (e.g., shared environment), covariance in the data (e.g., correlations across time), flexible modeling of longitudinal trajectories, and enabling discovery of time-dependent effects. Linear mixed-effects (LME) models are a powerful class of analytical strategies that can account for these issues in longitudinal data such as non-independence of the samples and other known sources of variance modeled as random effects, in addition to including variables with non-linear effects on the outcome and non-linear interactions. Examples of these random effects include the similarity of measurements for the same subject, variance explained by genetics (heritability), variance attributable to shared environment, etc.

When using LME models on longitudinal datasets, random effects should be specified as time-varying. For example, heritability can vary significantly over time (3–8). Similarly, correlations of measurements within subjects can change over time (e.g., autocorrelation). However, time-varying random effects are seldom modeled, given the significant computational burden of fitting such models. Thus, researchers use simplified LME models like compound symmetry (constant covariance over time) or making assumptions about the form of covariance (e.g., autoregressive models) which may not fit the data well. In addition, there is a lack of LME modeling software that scales to the needs of high-dimensional data (9) such as when estimating the marginal effects of several million single nucleotide polymorphisms (SNPs) in genome-wide association studies (GWAS), or when fitting mass-univariate models to millions of outcome variables, as is common in neuroimaging. This has severely limited our ability to maximize the use of high-dimensional longitudinal datasets.

Previously, we developed the Fast and Efficient Mixed-Effects Algorithm (FEMA) (10) for fitting a large number of LME models in a matter of seconds to minutes. FEMA massively accelerates LME fitting by using a covariance-regression approach to estimate random effects parameters and then quantizes these for vectorized fitting of many phenotypes simultaneously. For example, FEMA can fit ∼169,000 LME models with ∼10,000 observations in ∼54 seconds, a speedup of more than 3000 times, compared to standard LME solvers. In the present work, we introduce FEMA-Long, a framework adapted and optimized for high-dimensional longitudinal datasets. The features of FEMA-Long include i) modeling time-varying random effects using unstructured covariance, ii) modeling non-linear effects using splines, iii) examining time-dependent effects using spline interactions, and iv) performing GWAS, supporting the discovery of time-dependent effects. Through extensive simulations we show that FEMA-Long accurately models known sources of variance in a computationally fast manner while maintaining false positive rates. We demonstrate the use of FEMA-Long by performing a longitudinal GWAS on length, weight, and body mass index (BMI) of 68,273 infants with up to six repeated measurements (299,447 observations) during the first year of life, a highly dynamic period of development, and discovering SNPs showing time-dependent effects.

## Description of the method

### Nomenclature

We use the term *phenotype* to refer to a continuous outcome variable (or dependent variable) with the error in the phenotype approximately following a Gaussian distribution; we use the term covariates to refer to independent variables, inclusive of covariates of interest and covariates of no interest; and the term *random effects* refers to various known sources of nesting and similarity between observations in the data. For longitudinal data, the term *subject* or *participant* refers to a unique individual followed over time while the term *observation* refers to one measurement for a given subject. *Visit* or *timepoint* corresponds to an ordered set of repeated measurements for the same subject; for example, *visit 1* means the first measurement (first observation) on the subject, with increasing visit numbers indicating subsequent measurements. For mathematical notation, we use boldface for matrices or vectors, and regular font weight for scalar quantities.

### FEMA-Long algorithm overview

Given a set of continuous phenotype(s), covariates, and random effects, the goal is to estimate the fixed effects (beta coefficients for the covariates) while accounting for random effects. In the original FEMA framework (10), we first calculate the ordinary least squares (OLS) residuals of the phenotype; then, we use them to estimate the variance components of the random effects using method of moments (MoM) or a regression estimator with a non-negativity constraint (11). Then, the fixed effects are estimated using a generalized least squares (GLS) solution which incorporates a covariance matrix using the estimated variance components of the random effects. For scalability to large number of variables (such as for voxel-wise or vertex-wise data derived from brain imaging data), we use a binning approach – for all outcome variables that have similar variance components for the random effects, the GLS estimation is performed using a common covariance matrix that relies on the average of the variance components of all the outcome variables within that bin.

Here, we present FEMA-Long, an extension of FEMA that is adapted and optimized for large longitudinal datasets. The FEMA-Long framework includes i) time-varying covariances in the random effects using unstructured covariances; ii) modeling non-linear effects using splines; iii) examining time-dependent effects using spline interactions; and iv) performing GWAS and other similar omics analyses, where we test the association of a large number of variables of interest (e.g., genotyping matrix with millions of SNPs) with outcome(s), independently (i.e., marginal rather than joint effect), after controlling for covariates and random effects.

In FEMA-Long, we use unstructured covariance to model time-varying covariance of random effects. The elements of the unstructured covariance matrix are estimated by using the MoM estimation for every pair of visits that are nested within subjects. This results in a matrix of covariance terms for each random effect, which is then used for GLS estimation of the fixed effects. Importantly, this method does not assume a balanced design of all subjects having data for all the visits.

FEMA-Long also includes the option to model continuous outcome variables as a smooth function (for example, smooth function of age) using splines, represented by a set of basis functions. We also allow interaction terms between covariates, including those covariates that are modeled via splines. While the estimation of the unstructured covariance matrix uses pre-defined set of timepoints (visits), the spline modeling uses a continuous variable (such as age). This, therefore, brings the framework of generalized additive mixed models to FEMA, allowing flexible modeling of outcome variables as well as performing non-linear interaction analyses.

Finally, while the original FEMA was designed for many outcome variables and a common model (mass-univariate analyses), FEMA-Long includes a variant of FEMA that is intended for GWAS, where we fit the outcome(s) repeatedly on many models, each differing only in their genetic covariate. For this, we leverage the Frisch–Waugh–Lovell theorem (12,13): first, use stage-1 FEMA to estimate the random effects variance components and fixed effects (omitting genetic variants); then, we GLS residualize genotype vector for the covariates using the stage-1 variance components. Finally, we perform a GLS estimation using the GLS residuals of the phenotype and the GLS residuals of the genotype vector. This framework also allows modeling the interaction of genetic variants with spline basis functions, thereby allowing discovery of genetic variants that may show a nonlinear, time-dependent effect on the phenotype.

Further details on each of these steps is described below.

### General FEMA model set up (stage-1)

Let a dataset have *n* observations indexed with *i* ∈ 1 … *n*. Let ***X*** be a full-rank matrix of *p* covariates, with each column of ***X*** being a different covariate. ***X***_*i*_ are the values of the covariates for observation *i* (*X*_*i*_ is a scalar if *p* = 1 and a vector if *p* > 1). The phenotype is indicated by ***y***, thus *y*_*i*_ indicates the value of the phenotype for observation *i*. For brevity, we present the framework for a single phenotype (i.e., ***y*** is a vector), although FEMA fits the same model for each phenotype.

The linear model setup is:

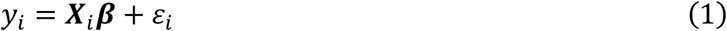

where ***β*** is a vector of *p* weights associated with each covariate, and *ε*_*i*_ is the unmodeled measurement error for each observation *i*. In a mixed modeling framework, we assume that the error term, *ε*, is normally distributed with zero mean and variance-covariance ***V***:

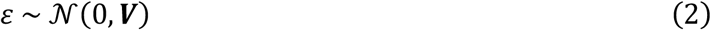

Then, the total residual variance is 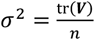. This normalized residual variance can be decomposed as a combination of random effects:

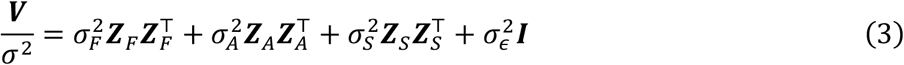

where, generically, ***Z*** is a random effects indicator matrix with entries of *Z*_*i*_ indicating membership of observation *i*; for example, in equation (3), ***Z***_*F*_ represents family with a value of 1 indicating observations that belong to the same family, entries of ***Z***_*A*_ represent the genetic relationship between a pair of observations, ***Z***_*S*_ represents subject effect with a value of 1 indicating repeated measurements for the same subject, and ***I*** represents the independent unmodeled variance across each observation; each random effect is associated with a corresponding variance component *σ*^2^. The choice of which random effects should be included depends on the data.

Within FEMA, we first use OLS to get an estimate of the fixed effects:

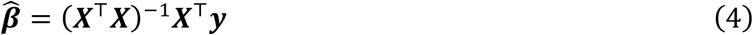

Then, the OLS residuals are:

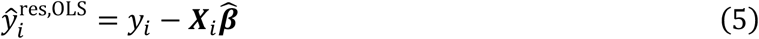

and the total residual variance 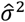 estimated as:

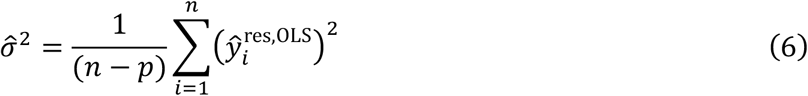

Following (3), for each pair of observations {*i, i*′}, the expected value of the product of corresponding residuals 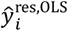 and 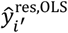 is:

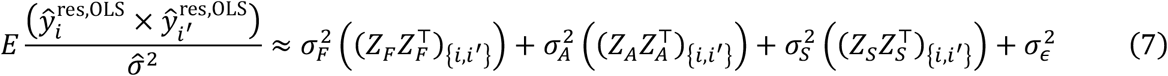

where the *E* operator is the expectation with respect to the random effects and generically (*ZZ* ^⊤^)_{*i,i*′}_ denotes the {*i, i*′} element of the random effects design matrix. Solving (7) results in the estimated values of random effects variance components, which can be used to construct a covariance matrix 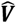:

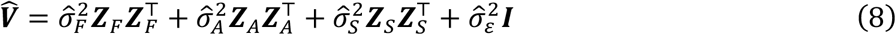

Finally, the stage-1 fixed effects can be estimated using the GLS solution:

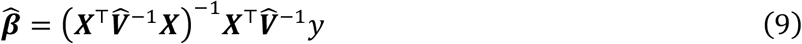

with variance

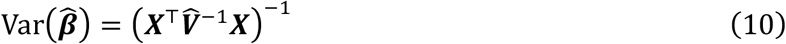

More details of these steps can be found in (10). We refer to this overall procedure as stage-1, which includes estimation of the random effects parameters as well as GLS estimation of the fixed effects coefficients.

### Estimating unstructured covariance

The original framework in FEMA assumed compound symmetry (i.e., constant covariance over time; sometimes also referred as diagonal covariance patterns since only the variance components are being modeled), an assumption which may not hold for longitudinal studies. In FEMA-Long, we allow modeling arbitrary unstructured covariance, where the parameters of the unstructured covariance matrix are estimated using the available data. Specifically, using the OLS residuals from equation (5), we solve equation (11) for every pair of visits (*v, v*′):

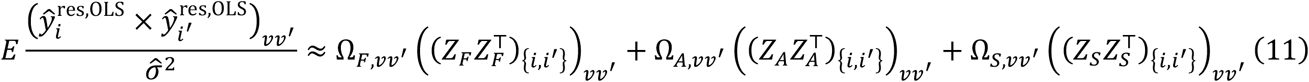

When solving (11), if *v* = *v*′, we use non-negativity constrained solution to estimate the variance component for that visit number *v*; if *v* ≠ *v*^′^, we drop the non-negativity constraint and estimate the covariance of the random effects between pairs of visits *v* and *v*^′^. Then, the estimated variance-covariance components can be put together to get *v* × *v* variance-covariance matrix for each random effect. Note that when estimating the variance components (i.e., the diagonal entries of the variance-covariance matrices), the subject and error terms are colinear, and therefore these variance parameters (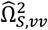 and 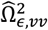) cannot be uniquely identified, and thus are estimated as one. Similarly, when estimating the covariance components (i.e., the off-diagonal entries of the variance-covariance matrices), the system of equations constructed from 11 makes the error term unidentifiable (see **Supplementary Figures S1** and **S2** for a demonstration of why these terms cannot be identified). The implication of this is that the estimated subject covariance matrix should be interpreted as describing the longitudinal stability of the phenotype within individuals, rather than as separate subject and noise variance components: the off-diagonal covariances between visits reflect the degree to which measurements remain stable within individuals over time, while diagonal elements capture the total variance at each visit, including both stable subject differences and visit-specific noise. If the estimated subject covariance matrix were purely diagonal, this would indicate little longitudinal stability and variability dominated by visit-specific noise; conversely, substantial off-diagonal covariance would indicate that measurements are correlated across visits, consistent with persistent subject-level differences over time.. This overall formulation does not require a balanced dataset – i.e., subjects with differing number of visits are allowed and the covariances are estimated according to the available data, with no observations being removed.

After estimating the elements of the unstructured covariance matrix, we construct the covariance matrix 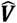 following equation (8)), with appropriate visit-wise reordering (such that the correct variance or covariance term for each random effect is used). Then, we use this covariance matrix 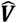 to perform GLS estimation for the fixed effects (equation (9)). When computing the standard errors using equation (10), we use a nearest symmetric positive semidefinite algorithm (based on (14) and implemented in (15)) to ensure that the covariance matrix is positive semidefinite; if this process fails, we generate a convergence warning message.

### Modeling spline basis functions and omnibus test

FEMA-Long allows modeling the outcome as a set of smooth basis functions *f*(*t*) (typically splines; indexed by *s* ∈ {1 … *S*}) of continuous variable ***t***:

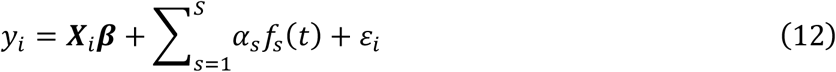

This leads to representing the variable ***t*** by ***S*** derived columns in the design matrix ***X***. Then, the set of coefficients, *α*_*s*_, define the non-linear function 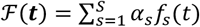, which represents the trajectory of the outcome over, for example, time. These basis functions can also be used to model interactions with another covariate.

In FEMA, the default option is to create natural cubic splines with unit height at knots, similar to the *nsk* option in the *splines2* package (16,17) in *R* (with intercept set as TRUE). Users can specify the knot placement based on domain knowledge, experiment design, or choose to place the knots at the quartiles of the variable. We also support creating other spline basis functions, namely natural cubic splines or B-splines which internally rely on a call to *ns* or *bs* functions in the *splines* package (18) in *R*.

By default, we modify the created basis functions such that the linear (or main) effect of ***t*** can be retained in the model. First, we create a linearly spaced vector of values ***t***_lin_ over the range of values in ***t***. Then, we create basis functions using these values. At this point, the user has an option of regressing out powers of ***t***_lin_ – for example, regressing out the zeroth and first powers of ***t***_lin_ would result in demeaned basis functions that are orthogonal to the linear effect of ***t***_lin_. Next, we perform a singular value decomposition on the demeaned basis functions. We then rescale the orthonormal basis functions such that each basis function has a bounded value between [−1, 1]. Finally, we linearly interpolate the modified basis functions to get corresponding basis function values for ***t***. These modified orthogonal basis functions are added as columns of ***X***. If the design matrix of the covariates already includes an intercept, then, the intercept taken together with these orthogonal basis functions covers the span of the variable.

Once the coefficients have been estimated, we can perform an omnibus Wald test to test the null hypothesis that the linear combination of the estimated coefficients is zero. Generically, given a vector or matrix of weights ***L*** with rank *r* equal to number of linearly independent rows, for a vector of estimated coefficients 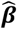 the multivariate Wald statistics *W* is (19):

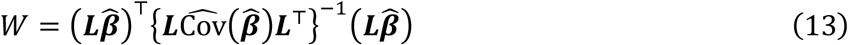

where 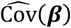 is the estimated variance-covariance matrix of the fixed effects. The Wald statistic *W* follows a *χ*^2^ distribution with *r* degrees of freedom. Note that equation (13) can be used to perform Wald tests on any combination of coefficients estimated in (12); i.e., not just limited to the estimated spline coefficients. Therefore, for an omnibus test of the null hypothesis that the linear combination of estimated coefficients of the basis functions is zero, ***L*** = ***I***(*S*), i.e., an identity matrix of size *S* (the number of basis functions). Alternatively, it is possible to use an *F* distribution to calculate the *p* value with two degrees of freedom: the numerator degree of freedom is the rank of ***L*** and the denominator degree of freedom is *n* − *p* − *S* (see (20) for a discussion on the similarity and differences in these approaches).

### Scaling FEMA-Long for GWAS-like analyses

Within GWAS, the goal is to estimate the effect of genetic variants on the phenotype(s), considering one variant at a time. Let ***G*** represent the time-invariant genotyping matrix, in which entry *g*_*i*_ ∈ {0, 1, 2} denotes the best-guess genotype (imputed genotypes which have a posterior probability above a threshold, followed by conversion to discrete count indicating the most probable genotype (21)) for the single nucleotide polymorphism (SNP) for observation *i*; 0 represents homozygous for the reference allele, 1 represents heterozygous, and 2 represents homozygous for the other allele. Let *γ*_*g*_ be the weight associated with this SNP ***g***. Since we are only interested in the marginal effect of each SNP ***g*** in ***G***, instead of fitting the full model (with covariates) for every SNP, we employ a two-stage procedure.

The Frisch-Waugh-Lovell theorem (12,13) states that the estimate of coefficients from a full regression model are equivalent to regression done in parts (or partial regressions) if both the *X* and *y* variables are residualized (22). This implies that we can estimate *γ*_*g*_ by first fitting a reduced model without genetic effect (stage-1), then residualizing each genetic variant ***g*** for the covariates, and finally fitting a second model using the residuals from stage 1 as the dependent variable and the residualized genotype as the independent variable (see (23) for a proof of the Frisch-Waugh-Lovell theorem in the GLS case).

Concretely, we first fit a reduced model excluding the genotyping vector. We estimate the variance components (compound symmetry or unstructured covariance) and the GLS estimates for the covariates using the stage-1 model as described previously. Then, the GLS residuals can be calculated as:

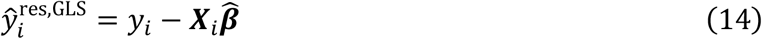

Next, we residualize ***g*** for the covariates. The coefficients of ***X*** on ***g, ζ***, can be calculated as:

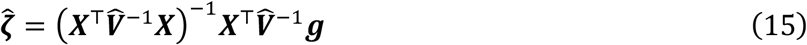

resulting in the GLS residuals:

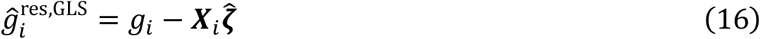

It is important to note that this residualized genotype vector 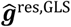 is phenotype specific. This follows from 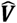 being composed from phenotype specific variance components (equation (8)). Now, using the stage-1 residuals 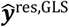 and the residualized genotype vector 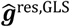, we can estimate 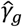 as:

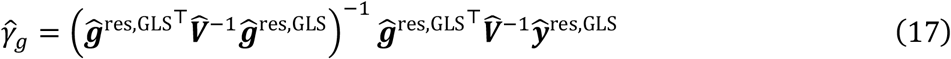

Then, for each genetic variant (and for each phenotype), the GLS residuals can be calculated as:

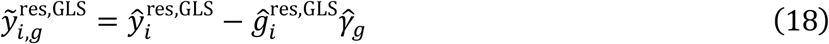

We use the notation 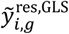 to distinguish these residuals (which are specific to each genetic variant) from 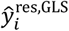 which are the stage-1 GLS residuals obtained from FEMA after accounting for the covariates (without the genetic variant). Therefore, the residuals 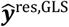 from (14) are the residualized phenotype for (17) which are estimated using the residualized genotype 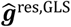 (from (16)).

The mean squared error is given by:

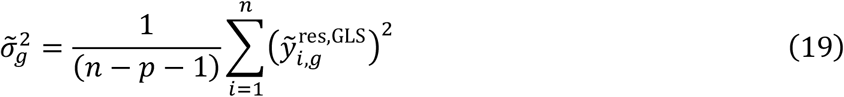

where the denominator is normalized by (*n* − *p* − 1) to account for the genotyping vector ***g*** as an additional independent variable (13,24). The variance for each 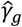 can be estimated as:

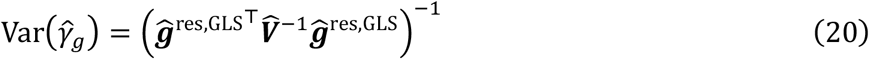

Since the 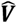 term has been normalized by 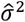, the standard error is scaled by 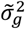.

The above framework can be applied to other GWAS-like analyses where one is interested in examining the effect of many predictors, taken one at a time (for example, given protein assays, the effect of each protein on a phenotype). Note, that in this framework, we assume that the individual contribution of each SNP is small and unlikely to impact the estimation of the covariance components in stage-1 (see Supplementary Material and Supplementary Figures S46-S51 for a comparison of the estimates between the two approaches).

### Modeling time-dependent genetic effects using spline interactions

This GWAS framework can also include interaction terms – linear interaction or smooth interaction. In case of linear interaction, if the main effect of the interacting variable has already been included as an *X* variable in stage-1 regression, then, ***g*** is a matrix with the first column containing the main effect of the genetic variant and the second column containing the elementwise (Hadamard) product of the genetic variant and the interacting variable. In this case, we residualize both columns of ***g*** for ***X*** (equation (15)), followed by estimation using equation (17), resulting in two coefficients: the coefficient for the main effect and the coefficient for the interaction term. The denominator of the mean squared error (equation (19)) is adjusted for two parameters, i.e., (*n* − *p* − 2).

For modeling spline interactions, ***g*** is a matrix having the elementwise product of the genetic variant and the smooth function *f*(*t*). The estimation follows the same procedure, with the estimation of as many coefficients as the columns of ***g***. If the basis functions have been appropriately modified and a constant term added (see the section on Modeling spline basis functions), then both the main effect of the genetic variant as well as the interaction of the genetic variant with smooth functions can be estimated. Generically, if there are *S* basis functions (with or without the constant term), the denominator in equation (19) is penalized by (*n* − *p* − *S*). Having estimated the coefficients, we can use equation (13) to either perform an omnibus test across the main and the time-dependent genetic effect, or perform an omnibus test for just the time-dependent effect and a separate test for just the main effect of the SNP (if desired). Depending on the number of tests performed, multiple comparison correction may be necessary.

Concretely, the GWAS interaction can be framed as modeling the effects of ***g*** on the outcome by altering regression coefficients *α*_*s*_ in (12):

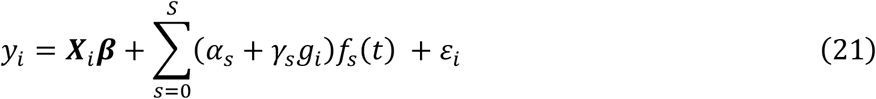

where *γ*_*s*_ coefficients are equivalent to the interaction term between variable ***g*** and the basis functions *f*(*t*). Note that *s* = 0 corresponds to the main effect of the SNP. This formulation is equivalent to:

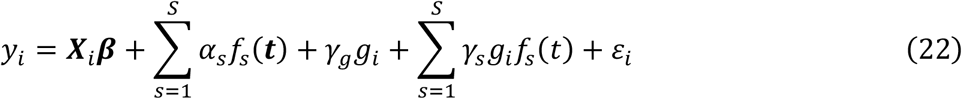

For example, if the time 𝓉 was modeled as smooth functions *f*(𝓉), then after estimating the regression coefficients, FEMA-Long can be used to calculate the following quantities:

The estimated cross-sectional effect of the genotype ***g*** on the outcome, at a specific time 𝓉_0_:

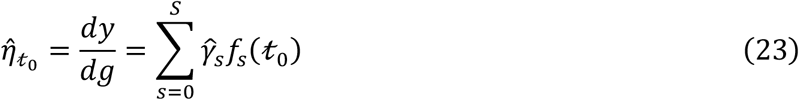

and the instantaneous effect of genotype ***g*** on the rate of change of the outcome at time 𝓉_0_:

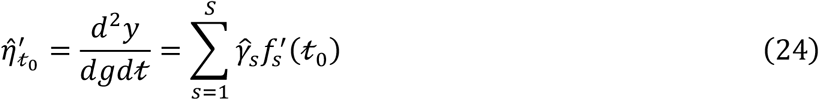

Both these quantities can be plotted as a function of time, 𝓉, to assess the effect of ***g*** on the outcome over time.

### Simulations

## Verification and Comparison

### Methods

To validate the FEMA-Long approach for fitting LMEs with unstructured covariances, we performed a series of simulations. All analyses were performed using MATLAB R2023a (25).

#### Simulation 1: parameter recovery

In the first simulation, we created phenotypes with known variance-covariance matrices and known effect sizes. Then, using FEMA, we estimated the parameters and compared them with the ground truth. We created a grid of known sources of variances: fixed effects Var_FFX_ ∈ {0.1, 0.2, …, 0.9}, family effect (i.e., variance attributed to common environment) Var_Fam_ ∈ {0.1, 0.2, …, 0.9}, subject effect (i.e., variance attributed to repeated measurements) Var_Sub_ ∈ {0.1, 0.2, …, 0.9}, and noise Var_Noise_ ∈ {0.1, 0.2, …, 0.9}. The total phenotypic variance was the sum of these four variances with the total variance being 1; i.e., Var_*y*_ = Var_FFX_ + Var_Fam_ + Var_Sub_ + Var_Noise_ = 1. This resulted in 84 combinations of variances.

For each of these combinations, we simulated data for 5,000 unique individuals with five repeated measurements; these individuals were grouped in families such that a family had no more than five members; then, we randomly subset the data to result in a total of 20,000 observations, reflecting repeated and missing measurements, as well as differing number of individuals across families. We simulated the age at each measurement using the mean and standard deviation of age as {0 ± 0; 60 ± 10; 180 ± 12; 240 ± 14; 365 ± 15} days – this scenario represents repeated measures data in infants with measurements conducted at birth, two months, six months, eight months, and one year.

For each of the 84 combinations of variances, we created an arbitrary 5 × 5 covariance matrix for family effect (V_Fam_) and subject effect (V_Sub_). We simulated 15 covariates– intercept, spline basis functions of the simulated age, and other random *X* variables drawn from a normal distribution. The ground truth beta coefficients were drawn from a uniform distribution *β*^sim^∼𝒰[−0.2, 0.2] such that *y*_FFX_ = *Xβ*^sim^, having mean 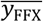 and standard deviation *σ*_FFX_; we then mean centered and scaled *y*_FFX_ to have Var_FFX_ variance. Then, using *V*_Fam_ and *V*_Sub_ covariance matrices, we drew samples from a multivariate normal (MVN) distribution with zero means and scaled them to have 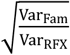 and 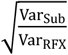 variances each, where Var_RFX_ = Var_Fam_ + Var_Sub_. In other words, 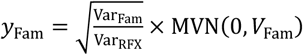) and 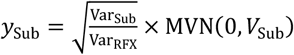. Therefore, 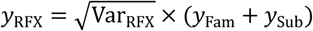. Finally, we simulated noise drawn from a uniform distribution and scaled it to have zero mean and Var_Noise_ variance. Thus, the simulated phenotype was *y* = *y*_FFX_ + *y*_RFX_ + *y*_noise_.

For this simulated phenotype, we fitted the model using FEMA, specifying the covariance type to be unstructured and the random effects to be family and subject effects. As an additional comparator, we fit the same model using the *glmmTMB* toolbox version 1.1.10 (26) in R version 4.2.1 (18). The model in glmmTMB was specified as: *y* ∼ 1 + *X* + us(timepoint + 0|FID) + us(timepoint + 0|IID), dispformula = ∼0, where 1 indicates the intercept, *X* indicates the remaining covariates (i.e., excluding the intercept), us specifics an unstructured covariance matrix for family (FID) and subject (IID) terms, the timepoint denotes a factor variable indicating which visit the observation belonged to, and the dispformula component disables the addition of an additional error term as that would be an overparameterized model (27). After preliminary testing, we noticed convergence issues with glmmTMB; therefore, we changed the default settings to have higher number of iterations, specifically setting iter.max and eval.max to 5,000.

We compared the beta coefficients for the covariates and the estimated variance-covariance matrices for the family and subject effects from both FEMA-Long and glmmTMB with the ground truth and with each other. For the beta coefficients, 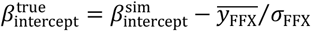, and for the remaining *X* variables, 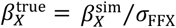; for the family effect variance-covariance matrix, 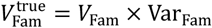; and for the subject effect variance-covariance matrix, 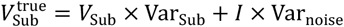, where *I* was an identity matrix of size five (corresponding to number of visits). Then, we calculated the root mean squared error (RMSE) for the fixed effects and the family and subject variance-covariance matrices to assess how far the estimated values were from the true simulated parameters, and additionally to quantify the difference between the estimated parameters from FEMA-Long and glmmTMB. For each of the 84 combinations of variances explained by covariates and random effects, we repeated this entire process 50 times.

#### Simulation 2: false positive rate

To evaluate the false positive rate, we repeated the strategy from simulation one. The covariates included the intercept, age, and other random 98 *X* variables drawn from a random normal distribution, resulting in 100 *X* variables. However, the covariates had zero contribution to the simulated phenotype. The total phenotypic variance was 1, i.e., Var_*y*_ = Var_Fam_ + Var_Sub_ + Var_Noise_ = 1, resulting in 36 different combinations. For each simulation setting, we created 10 *y* variables that have different amount of contribution from family, subject, and noise terms and repeated these simulations 1,000 times. Using these simulated data, we fitted the models using the unstructured covariance in FEMA-Long as well as repeated the model fitting using compound symmetry covariance type. Then, we examined the distribution of the *p*-values from both the models.

#### Simulation 3: computational time and carbon footprint

To assess the computational performance of FEMA-Long with unstructured covariance, we simulated data with varying number of observations and varying number of outcome variables. We fixed Var_Fam_ = 0.20, Var_Sub_ = 0.30, Var_FFX_ = 0.35, and Var_noise_ = 0.15. We set the number of covariates to 50. We created 10 log-spaced values for the number of observations (rounded off), varying between 10^0^ and 10^5^. Similarly, we created 10 log-spaced values for the number of outcome variables (rounded off), varying between 10^0^ and 10^4^. For each of these combinations of number of observations and number of outcome variables, we estimated the time taken by FEMA. We repeated the time measurement five time and then took the median to get a robust estimate of the time taken to run the analysis. We also performed the time measurement using parallel computing with 32 parallel jobs and 2 threads per job. As comparators, we used the glmmTMB toolbox and the *lme4* (28) toolbox (version 1.1-28); since these toolboxes are not designed for multiple outcome variables, we only timed it for one outcome variable (across the 10 values of number of observations) and then scaled the timing for larger number of outcome variables. For fitting the models with *lme4*, we used the *lmer* function, specifying the random effects as timepoint + 0|FID and timepoint + 0|IID; in addition, we specified a *lmer* control object that disabled the check for number of observations being greater than or equal to the number of random effects levels (i.e., we set check.nobs.vs.nRE to ignore). We ran the timing measurements on a compute cluster with two AMD EPYC 7702 64-core processors, with the job specification of 64 CPUs per task and 5GB memory per CPU. In addition, to estimate the carbon footprint of FEMA-Long, we used the formula from (29) and the data from the green-algorithms.org v3.0. We specified the number of cores as 64 and the memory usage as 320 GB; we set the CPU model to “Any” and the location to “World”. For comparison, we also estimated the carbon footprint for the glmmTMB and the lme4 toolboxes.

#### Simulation 4: impact of sample size and missingness on parameter recovery

Since FEMA-Long is designed for large sample sizes and the estimation of the covariance parameters uses pairs of timepoints, we performed an additional analysis to examine the impact of missing observations on parameter recovery. Specifically, we simulated data for 5,000 unique individuals with five repeated measurements; these individuals were assigned family numbers such that a family had no more than five members. The probability of a missing visit was between 0.1 and 0.8, while ensuring that subsequent visits had a higher missingness probability; since there are five repeated measurements, there are 15 possible visit combinations (five variances and ten covariances) – we varied the minimum number of observations between any pairs of visits min_numObs_ between 500 to 800, in increments of 100; the resulting total number of observations *n*_Obs_ were 12,000, 15,000, 18,000, and 20,000. To clarify, there were 16 scenarios: four min_numObs_ and four *n*_Obs_. For simulating arbitrary covariance matrices (family and subject covariances), we allowed the within-visit variance to vary between 0.2 and 0.8, and the correlation between visits (scaled to covariances) to vary between -0.7 and 0.7; after ensuring that the simulated covariance matrix was positive semidefinite, we additionally ensured that the condition number of the matrices did not exceed 1000. Then, for these 16 scenarios, the rest of the simulation strategy was the same as simulation 1. We compared the estimated parameters with the ground truth and with glmmTMB as a comparator for 16 scenarios, each having 84 combinations of variance values, each of them having 15 covariates, one outcome variable, and the process repeated 50 times.

#### Simulation 5: impact of sample size and missingness on false positives

Similar to simulation 4, to examine the impact of missing observations on the number of false positives, we followed the simulation strategy from simulation 4 to create 16 different scenarios with differing min_numObs_ and *n*_Obs_. We examined the distribution of the *p*-values (using unstructured covariance and compound symmetry) for the 16 scenarios, each having 36 combination of variance values (similar to simulation 2), each of them having 100 covariates, 10 outcome variable, and the entire process repeated 1000 times.

### Results

#### Simulation 1: FEMA-Long accurately models unstructured covariances

For the fixed effects, the average (over 50 iterations) RMSE across 84 simulations was 0.0121 ± 0.0018 [0.0073 − 0.0150], *r*_true,FEMA_ = 0.996 (**Figure 1a, Supplementary Figure S3**). For the random effects parameters, the average RMSE for the family effect was 0.0160 ± 0.0058 [0.0064 − 0.0312], *r*_true,FEMA_ = 0.997 (**Figure 1b, Supplementary Figure S4**), and for the subject effect was 0.0122 ± 0.0036 [0.0051 − 0.0218], *r*_true,FEMA_ = 0.999 (**Figure 1c, Supplementary Figure S5**), indicating that the estimates were close to the ground truth.

**Figure 1:**
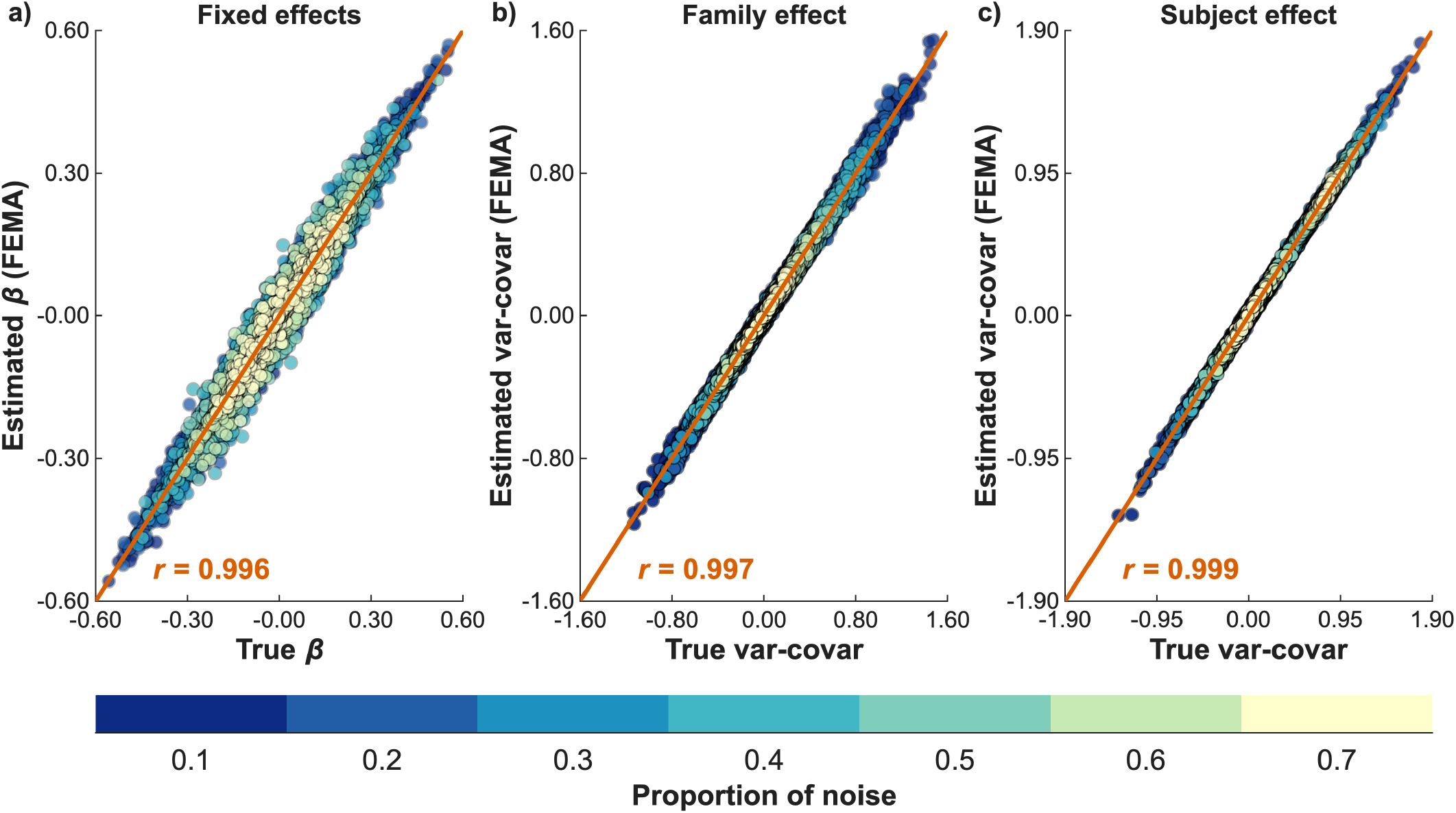
Comparison of estimated parameters from FEMA-Long with ground truth. Each panel shows the scatterplot of the simulated ground truth and the estimated parameters from FEMA-Long across 50 iterations of 84 simulation settings with different amounts of variances explained by fixed, family, and subject effects, as well as noise; the amount of noise in the simulated phenotype is indicated by the color of the points; the orange line indicates the least square fit and the correlation between ground truth and estimated parameters are indicated in orange text; **a)** beta coefficients for the covariates; **b)** variance-covariance components for the family random effect; and **c)** variance-covariance components for the subject effect.

The average RMSE between the estimates from FEMA-Long and from glmmTMB toolbox for the fixed effects was 0.0004 ± 0.0001 [0.0002 − 0.0007], *r*_glmmTMB,FEMA_ = 1.00 (**Supplementary Figure S6**), for the family effect was 0.0076 ± 0.0028 [0.0028 − 0.0151], *r*_glmmTMB,FEMA_ = 0.9996 (**Supplementary Figure S7**), and for the subject effect was 0.0055 ± 0.0016 [0.0020 − 0.0104], *r*_glmmTMB,FEMA_ = 0.9998 (**Supplementary Figure S8**), showing that the FEMA-Long estimates were almost identical to glmmTMB.

#### Simulation 2: FEMA-Long controls for false positives

Across the 36 simulations, when using unstructured covariance, the *p*-values were uniformly distributed under the null (− log_10_ *p*-values following the diagonal; green points in **Figure 2** and **Supplementary Figure S9**). However, for the compound symmetry covariance pattern, in almost all scenarios, the *p*-values had a non-uniform distribution with several of the *p*-values being smaller than expected (leftward deflection of the − log_10_ *p*-values from the diagonal; orange points in **Figure 2** and **Supplementary Figure S9)**. These results show that the unstructured covariances *p*-values are well-calibrated.

**Figure 2:**
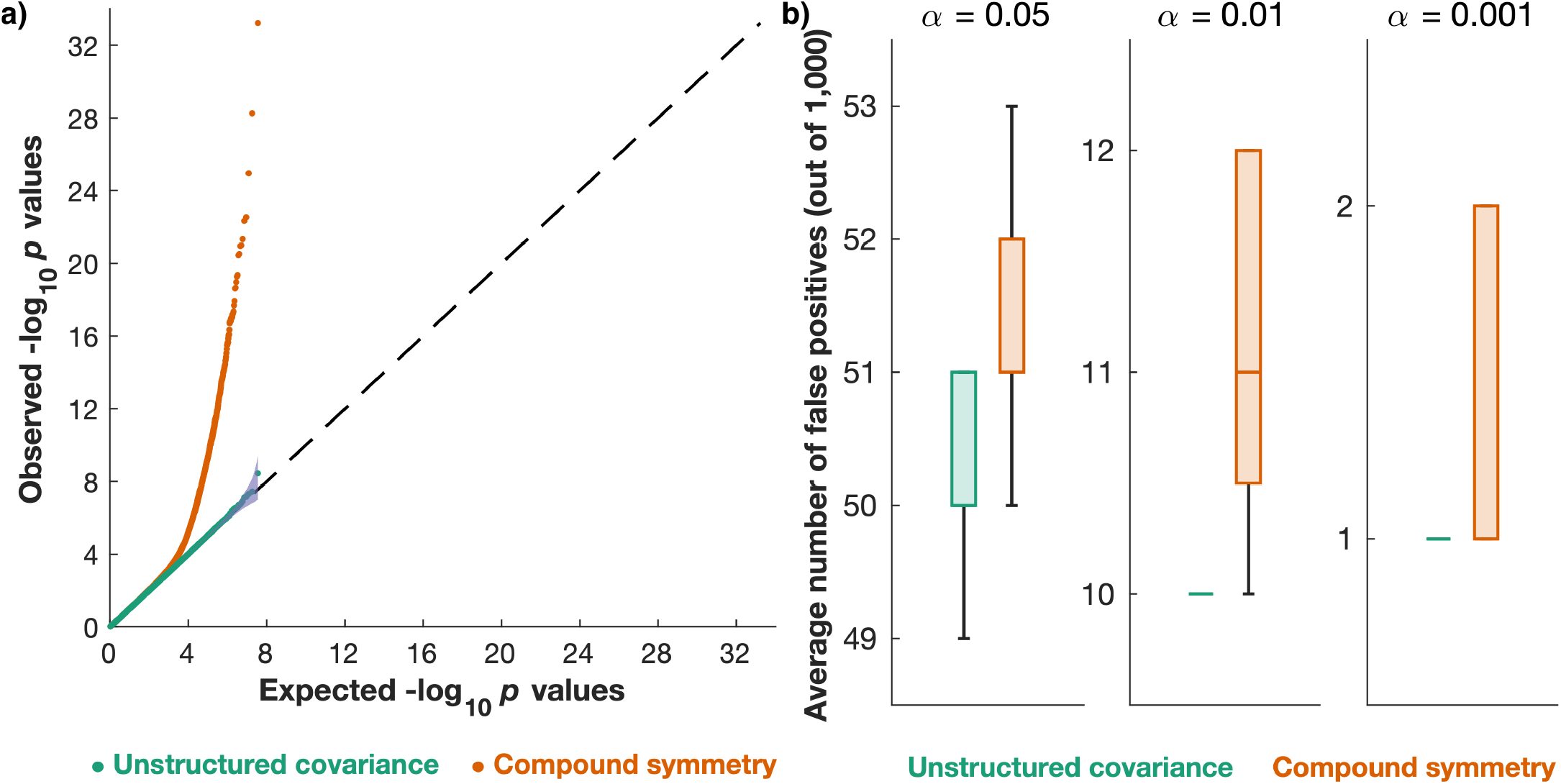
Examining false positives under the null. **a)** Distribution of − log_10_ *p*-values across 1000 iterations of 36 simulation settings with differing amounts of variances explained by family effect, subject effect, and noise for unstructured covariance (green points) and compound symmetry (orange points); each simulation iteration consisted of 100 *X* variables and 10 outcome variables; the purple filled area indicates the 95% confidence interval based on inverse beta distribution; **b)** box-plots showing the average number of false positives (rounded off) at different alpha values across 100 *X* variables and 10 *y* variables for unstructured covariance (green bars) and compound symmetry (orange bars); the whiskers mark the values 1.5 times the interquartile range away from 75^th^ and 25^th^ percentile; under the null, the expected number of false positives are 50 (α = 0.05), 10 (α = 0.01), and 1 (α = 0.001).

#### Simulation 3: FEMA-Long is faster and greener

FEMA-Long was fast across a range of observations and outcomes (**Figure 3**). For a single outcome, for 1,000 observations, FEMA-Long took ∼0.14 seconds; in comparison, glmmTMB took ∼9.76 seconds and lmer took ∼85.66 seconds (i.e., FEMA-Long was ∼71 times faster than glmmTMB and ∼622 times faster than lmer); for 100,000 observations, this time went up to ∼34.8 seconds (FEMA-Long; **Figure 3a**), ∼1013 seconds (glmmTMB; **Supplementary Table S1**), and ∼3105 seconds (lmer; **Supplementary Table S5**); therefore, for a single phenotype, FEMA-Long was 29-106 times faster than glmmTMB and 89-622 times faster than lmer. The carbon footprint for glmmTMB for a single outcome varied between 1.78 gCO_2_e and 185.01 gCO_2_e; for lmer the carbon footprint varied between 8.52 gCO_2_e and 567 gCO_2_e; in comparison, for FEMA-Long, the carbon footprint was between 0.03 gCO_2_e and 6.35 gCO_2_e; therefore, similar to computational timing, FEMA-Long was between 29 and 106 times greener than glmmTMB (**Supplementary Table S7**) and between 89 and 622 times greener than lmer (**Supplementary Table S9**).

**Figure 3:**
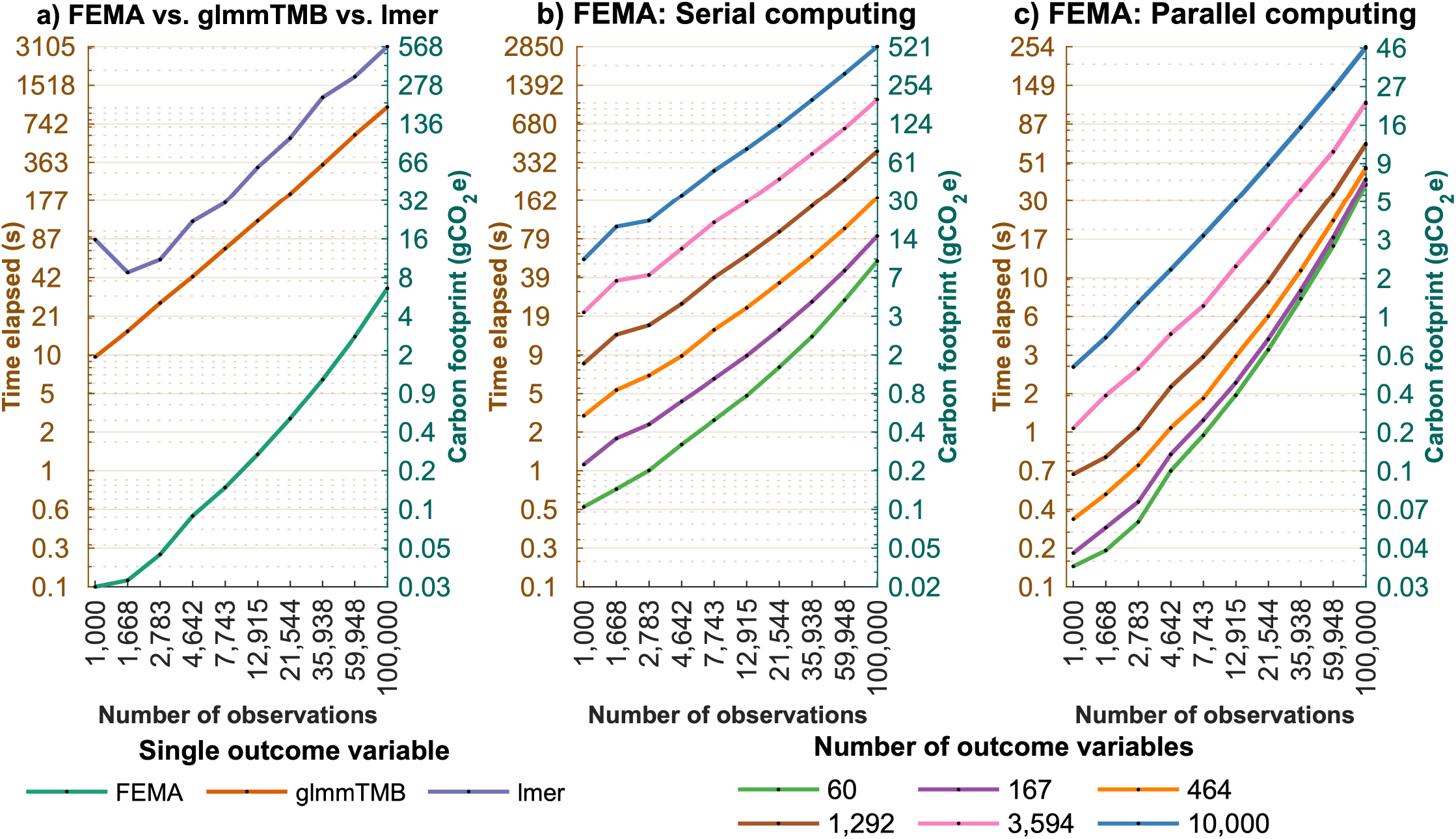
Time taken and carbon footprint for fitting linear mixed-effects models with unstructured covariances. In each panel, the line(s) indicate the time taken (in seconds) on the left *y*-axis and the carbon footprint (in grams of carbon dioxide emission) on the right *y*-axis for fitting linear mixed-effects models with unstructured covariances for family and subject random effects and 50 covariates across a range of number of observations (shown on the *x*-axis). Panel **a)** shows the time taken by and the carbon footprint for FEMA-Long (green), glmmTMB (orange), and lmer (from the lme4 toolbox; purple) for a single outcome variable across a range of number of observations. Panel **b)** shows the time taken by and the carbon footprint of FEMA-Long with serial computing, and panel **c)** shows the time taken by and the carbon footprint of FEMA-Long with parallel computing (32 parallel processes) across a range of increasing number of outcome variables, indicated by the color of the line. In all the panels, both axes are log-spaced and the *y*-axes tick labels are rounded off.

For 100,000 observations and 10,000 outcome variables, FEMA-Long took ∼47.5 minutes and had a carbon footprint of 520.51 gCO_2_e (serial, **Figure 3b** and **Supplementary Tables S2** and **S7**) and ∼4.2 minutes with a carbon footprint of 46.46 gCO_2_e (parallel, **Figure 3c** and **Supplementary Tables S2** and **S7**); in contrast, extrapolating the timing for glmmTMB, the same analysis would have taken ∼117.25 days and would have had a carbon footprint of 1.85 TCO_2_e (**Supplementary Tables S2** and **S7**); similarly, extrapolating the timing for lmer, the same analysis would have taken ∼359.36 days and would have had a carbon footprint of ∼5.67 TCO_2_e (**Supplementary Tables S6** and **S9**). Overall, for more than one phenotype, across a range of observations, FEMA-Long was between ∼86-3,554 times faster and greener than glmmTMB (serial) and between ∼84-42,096 times faster and greener than glmmTMB (parallel; **Supplementary Table S2** and **Supplementary Table S7**); compared to lmer, FEMA-Long was between ∼263-15,785 times faster and greener (serial) and between ∼259-287,190 times faster and greener (parallel; **Supplementary Tables S6** and **S9**). We encountered convergence issues with glmmTMB; increasing the maximum number of iterations and evaluations to 5000 and re-examining the time taken by glmmTMB, we found FEMA-Long to be up to 6,902 (serial) and 93,659 times faster and greener (parallel) than glmmTMB (**Supplementary Table S3, S4**, and **S8**).

#### Simulation 4: FEMA-Long estimates are stable across sample sizes and missingness

When we compared the FEMA-Long estimates with the ground truth across a range of sample size and differing minimum number of observations between pairs of visits, we found the FEMA-Long estimates to be highly correlated to the ground truth (*r*_true,FEMA_ = 0.987 − 0.999; **Supplementary Figures S10 – S25**). The average (over 50 iterations) RMSE across 84 simulations for the fixed effects ranged between 0.0150 ± 0.0025 (*n*_Obs_ = 12,000, min_numObs_ = 500) and 0.0111 ± 0.0018 (*n*_Obs_ = 15,000, min_numObs_ = 800); for the family effect covariance parameters, the average RMSE ranged between 0.0128 ± 0.0033 (*n*_Obs_ = 12,000, min_numObs_ = 500) and 0.0085 ± 0.0025 (*n*_Obs_ = 15,000, min_numObs_ = 800); for the subject effect covariance parameters, the average RMSE ranged between 0.0152 ± 0.0042 (*n*_Obs_ = 12,000, min_numObs_ = 500) and 0.0082 ± 0.0027 (*n*_Obs_ = 15,000, min_numObs_ = 800); see **Supplementary Table S10** for details. For smaller *n*_Obs_ such as 12,000 observations, increasing min_numObs_ resulted in reduced RMSE with the ground truth; at larger *n*_Obs_ such as 20,000 observations, we did not see any differences when increasing min_numObs_, likely indicating a stable performance for large samples.

On comparing the FEMA-Long estimates with glmmTMB across a range of sample sizes and minimum number of observations between pairs of visits, the FEMA-Long parameters were comparable to the glmmTMB estimated parameters (**Supplementary Table S11**). The average (over 50 iterations) RMSE across 84 simulations for the fixed effects ranged between 0.0006 ± 0.0002 (*n*_Obs_ = 12,000, min_numObs_ = 500) and 0.0002 ± 0.0001 (*n*_Obs_ = 15,000, min_numObs_ = 800); for the family effect covariance parameters, the average RMSE ranged between 0.0053 ± 0.0021 (*n*_Obs_ = 12,000, min_numObs_ = 500) and 0.0022 ± 0.0010 (*n*_Obs_ = 15,000, min_numObs_ = 800); for the subject effect covariance parameters, the average RMSE ranged between 0.0064 ± 0.0019 (*n*_Obs_ = 12,000, min_numObs_ = 500) and 0.0024 ± 0.0007 (*n*_Obs_ = 15,000, min_numObs_ = 800). Similar to the comparison with ground truth, we observed that for smaller *n*_Obs_ such as 12,000 observations, increasing min_numObs_ resulted in reduced RMSE, while at larger *n*_Obs_ such as 20,000 observations, increasing min_numObs_ did not make a difference to the average RMSE.

#### Simulation 5: FEMA-Long false positive calibration improves with sample size

We examined the distribution of *p*-values under a range of *n*_Obs_ and min_numObs_; in each of these 16 scenarios, across the 36 simulations, unstructured covariance generally resulted in uniformly distributed *p*-values (**Supplementary Figures S26, S29, S32**, and **S35**), except for two settings: Var_Fam_ = 0.7, Var_Sub_ = 0.2, Var_Noise_ = 0.1 and Var_Fam_ = 0.8, Var_Sub_ = 0.1, Var_Noise_ = 0.1; in these two settings, we observed that there were a few combinations of *n*_Obs_ and min_numObs_, where the distribution of the unstructured covariance *p*-values closely followed the null distribution but deviated away from the null towards the tail of the distribution: *n*_Obs_ = 12,000, min_numObs_ = 500 (**Supplementary Figure S27**), *n*_Obs_ = 12,000, min_numObs_ = 600 (**Supplementary Figure S28**), *n*_Obs_ = 15,000, min_numObs_ = 500 (**Supplementary Figure S30**), *n*_Obs_ = 15,000, min_numObs_ = 600 (**Supplementary Figure S31**), *n*_Obs_ = 18,000, min_numObs_ = 600 (**Supplementary Figure S33**), and *n*_Obs_ = 18,000, min_numObs_ = 700 (**Supplementary Figure S34**). On further examination of these cases, we observed that the miscalibration of the *p*-values was caused by one or few iterations (out of the 1000) where the resulting distribution of visits was extremely imbalanced (**Supplementary Figure S36**), leading to a small number of common observations between the last pairs of visits, possibly leading to an unstable estimation of the covariance components. As the sample size increased, the false positive calibration for the unstructured covariance improved; in contrast, the compound symmetry covariance pattern resulted in inflated Q-Q plots.

## Application: GWAS on anthropometric features in infants

### Study sample

To demonstrate the features of FEMA-Long (unstructured covariance, non-linear modeling, scalability to large datasets, performing GWAS, and discovery of time-dependent genetic effects), we performed a GWAS on the length, weight, and BMI of infants in the first year of life using the large-scale Norwegian Mother, Father, and Child Cohort Study (MoBa) (30–32). The MoBa Study is a population-based pregnancy cohort study conducted by the Norwegian Institute of Public Health. Participants were recruited from all over Norway from 1999-2008. The women consented to participation in 41% of the pregnancies. The cohort includes approximately 114,500 children, 95,200 mothers and 75,200 fathers. The current study is based on version 12 of the quality-assured data files released for research in 2019. The establishment of MoBa and initial data collection was based on a license from the Norwegian Data Protection Agency and approval from The Regional Committees for Medical and Health Research Ethics. The MoBa cohort is currently regulated by the Norwegian Health Registry Act. The current study was approved by The Regional Committees for Medical and Health Research Ethics (2016/1226/REK Sør-Øst C).

### Genetic data

Blood samples were obtained from both parents during pregnancy and from mothers and children (umbilical cord) at birth (33), which were used for genotyping. We used the quality controlled genetic data from the MoBaPsychGen pipeline v1 (34). The release consists of data from 76,577 children, 77,634 mothers, and 53,358 fathers; we restricted ourselves to the data on children who had not withdrawn their consent from MoBa as of November 1, 2024.

### Phenotypes

The phenotypes of interest were the length, weight, and BMI of infants at six timepoints during the first year of life: birth, six weeks, three months, six months, eight months, and twelve months. Phenotypic information was retrieved from questionnaire data, and from the Medical Birth Registry of Norway (MBRN); the MBRN is a national health registry containing information about all births in Norway (see Supplementary Materials for details on data curation and quality checks prior to GWAS). The final sample size for GWAS was 68,273 infants with 299,447 observations having complete data on length, weight, and BMI (i.e., each of the 68,273 subjects had all three measurements); see **Supplementary Figure S10** for distribution of age at each timepoint, and **Supplementary Figures S11-S13** for plots showing the trajectories of length, weight, and BMI.

### Genome-wide association study

For the 68,273 infants, with up to six repeated measurements totaling 299,447 observations on length, weight, and BMI during the first year of life, we performed a longitudinal GWAS using FEMA-Long on the autosomes. We standardized the phenotypes to have zero mean and unit standard deviation. The covariates included the intercept, six spline basis functions of age (after the transformations described in the Methods section), dummy-coded sex, twenty genetic principal components (PC), and 23 dummy-coded genotyping batch variables, resulting in a total of 51 covariates. We created the spline basis functions by placing a knot at the median age for each timepoint: 44, 94, 181, 246, and 369 days, and boundary knots at the extremes (0 and 425 days; see, **Supplementary Figure S14** and **Supplementary Figure S15**).

The random effects included the effect of family or shared environment, the genetic relationship matrix (GRM), and the subject effect (or repeated measurements). We defined families using the mother ID of the participants; in addition, for a subset of participants where the mother ID was different, but the father ID was the same, we assigned those individuals to the same family; this resulted in 46,453 unique families. To calculate GRM, we used a subset of 500,243 SNPs which were directly genotyped in at least one imputation batch or had an imputation INFO score > 0.99785 in all imputation batches; we mean imputed any missing genotyping data and standardized the SNPs (zero mean, unit standard deviation); then, the GRM was calculated as the correlation coefficient between the standardized SNPs. Therefore, the stage-1 model (without GWAS) for each phenotype was:

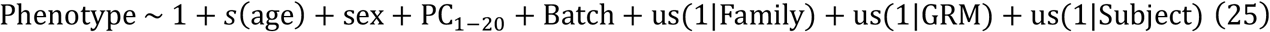

We performed GWAS on 6,981,748 SNPs (34) after mean imputing missing genotyping data, and examined both the main effect of the SNPs as well as the interaction of each SNP with the six basis functions, resulting in seven coefficients per SNP per phenotype. Specifically, after fitting the stage-1 model and residualizing each genotype for the covariates in (25), the model for each residual SNP, *G*_res_, was:

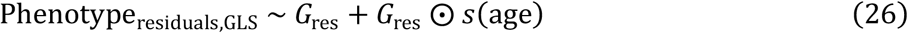

where *G*_res_ ⨀ *s*(age) is the element-wise multiplication of *G*_res_ with every column of the basis function matrix. Then, for each phenotype, and each SNP, we performed a Wald test to determine if any of these coefficients had a non-zero effect. To accelerate computing, we split the genetic data into chunks of 5,000 SNPs, processing eighteen chunks in parallel, with each parallel worker having seven threads, run on a compute cluster with two AMD EPYC 7702 64-core processers; the job specification specified 128 CPUs per task and 30 GB RAM per task. Additionally, we also ran a standard longitudinal GWAS, without including *s*(age) and *G*_res_ ⨀ *s*(age) terms (run on the same hardware with the same specification, but twenty parallel chunks with six threads each).

To quantify genomic inflation factor (*λ*_GC_), we used the Wald test *p* values, converted them to chi-squared statistics (using 1 − *p* values as input to the inverse of the chi-square cumulative distribution function with 1 degree of freedom); then, *λ*_GC_ was calculated as:

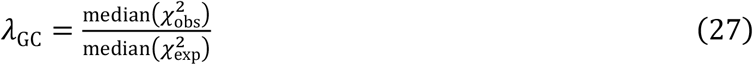

where 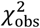 were the observed chi-squared statistics, while 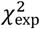 were the expected chi-squared statistics under the null (this simplifies to using the chi-squared statistics for a value of 0.5, since the under the null, the *p* values are uniformly distributed). We quantified *λ*_GC_ for both the scenarios: when allowing SNP effects to vary as smooth function of age, and when performing a standard longitudinal GWAS without any interactions or smooth terms.

### Results: examining unstructured covariances and discovery of age-dependent genetic effects

The stage-1 analysis was completed in ∼3 minutes; examining the random effects unstructured covariance revealed dynamic patterns over time (**Figure 4**). Specifically, the proportion of within-visit variance attributed to genetic effects (within-visit heritability) for each phenotype showed differences across timepoints, ranging between 0.42–0.91 for length, 0.45–0.84 for weight, and 0.55–0.73 for BMI (diagonal entries, middle column, **Figure 4**). Interestingly, genetic correlations between timepoints (off-diagonal entries, middle column, **Figure 4)** also showed fluctuations over time with different patterns across phenotypes. For example, the correlations between birth and other timepoints for length and weight were fairly stable, while BMI showed a decaying pattern. The heritability was relatively lower for the first three timepoints for length and weight compared to other timepoints, while the heritability for BMI was relatively stable from 3m to 12m period. The family effect explained a small amount of variance for the first three timepoints for length and weight, but not as much for BMI. The variances explained by subject effect (and noise) also changed over time: a stronger effect for the first three timepoints for length but not as much for later timepoints, and a similar trend for weight and BMI but with greater variance explained for later timepoints.

**Figure 4:**
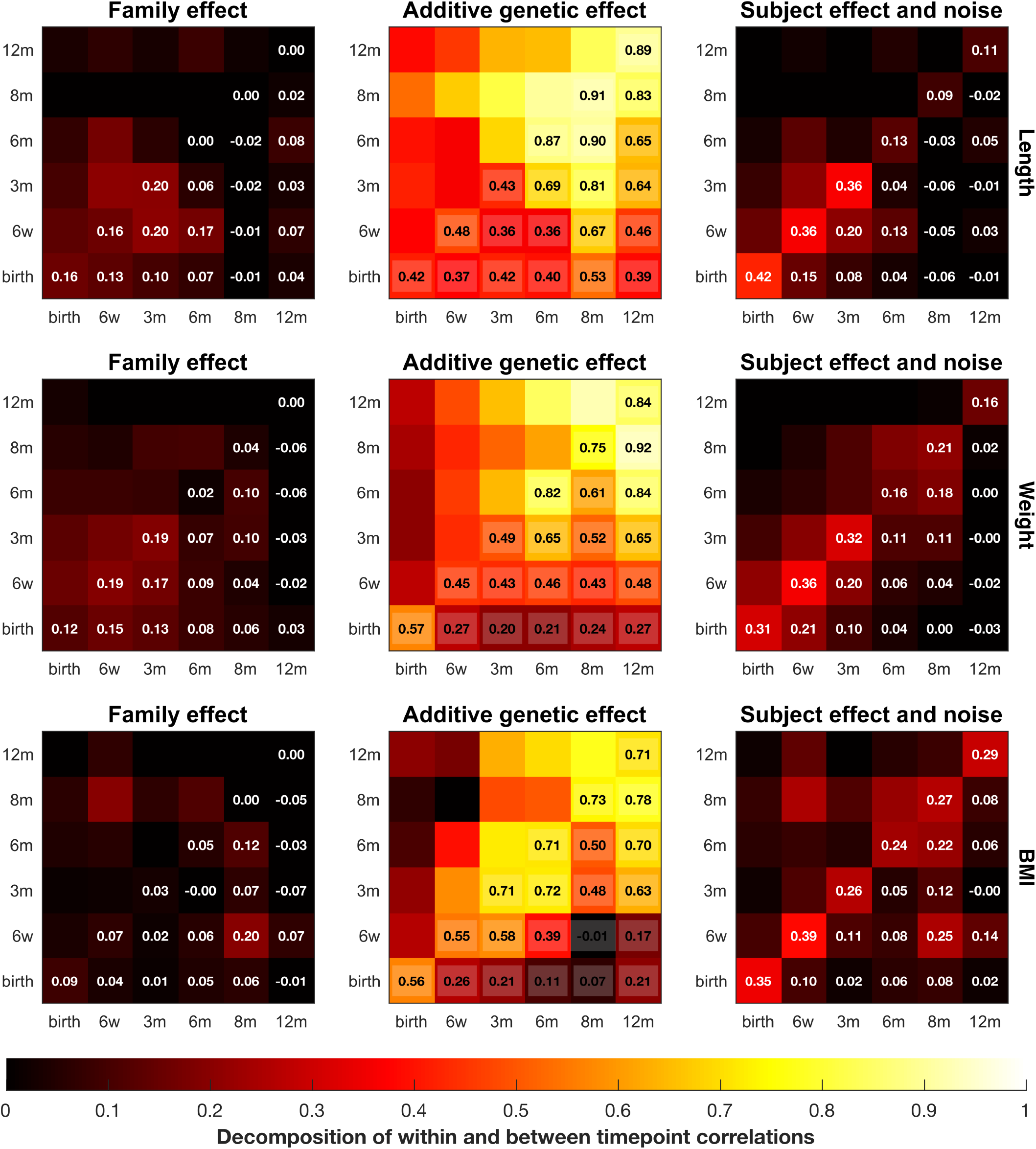
Decomposition of within and between timepoint correlations into family, additive genetic, and subject effects. The panels are the normalized unstructured covariance matrices for length, weight, and BMI in infants in the first year of life; the diagonal values are the variances explained by shared family environment, the additive genetic effect (heritability), and the subject effect (which also includes noise) for each timepoint while the off-diagonals reflect the correlations between time points.

The GWAS analysis, completed in ∼16.88 hours, revealed 2,836, 4,367, and 6,707 SNPs showing a main effect or a time-dependent effect on length (*λ*_GC_ = 1.10), weight (*λ*_GC_ = 0.91), and BMI (*λ*_GC_ = 1.36) respectively at *α* = 5 × 10^−8^; in comparison, longitudinal GWAS (without including splines or interactions; analysis completed in ∼3.13 hours) revealed 2,126, 660, and 2,797 significant SNPs for length (*λ*_GC_ = 1.25), weight (*λ*_GC_ = 1.13), and BMI (*λ*_GC_ = 1.21) respectively (**Figure 5a, Figure 6a**, and **Figure 7a**). Here, the *λ*_GC_ > 1 is in line with other recent large-scale GWAS analyses and likely reflects the high polygenicity of the traits (see **Supplementary Figures S43-S45** for Q-Q plots). We used the SNP2GENE module of FUMA (35) to examine the number of independently significant SNPs and the number of identified genetic loci. For length, FUMA identified 134 independent SNPs and 54 loci when allowing the SNP effect to vary over time, as compared to 126 independent SNPs and 49 loci when modeling only the longitudinal main effect of SNPs; for weight, FUMA identified 201 independent SNPs and 46 loci when allowing the SNP effect to vary over time, as compared to 45 independent SNPs and 23 loci when modeling only the longitudinal main effect of SNPs; for BMI, FUMA identified 242 independent SNPs and 59 loci when allowing the SNP effect to vary over time, as compared to 124 independent SNPs and 44 loci when modeling only the longitudinal main effect of SNPs. We examined the fitted spline trajectories for the top 10 SNPs for each trait, one SNP per chromosome (**Figure 5b, Figure 6b**, and **Figure 7b**. For length, most of the effects were linear; however, the selected SNPs for weight and BMI showed more dynamic patterns with non-linear trajectories. Notably, even the same selected SNPs for weight and BMI showed differing patterns in the direction and timing of the effect changes across the first year of life (e.g., rs2767486 and rs13322435 (3)). These trajectories highlight the importance of modeling the time-dependent effects and reveal how the SNP effects change over time.

**Figure 5:**
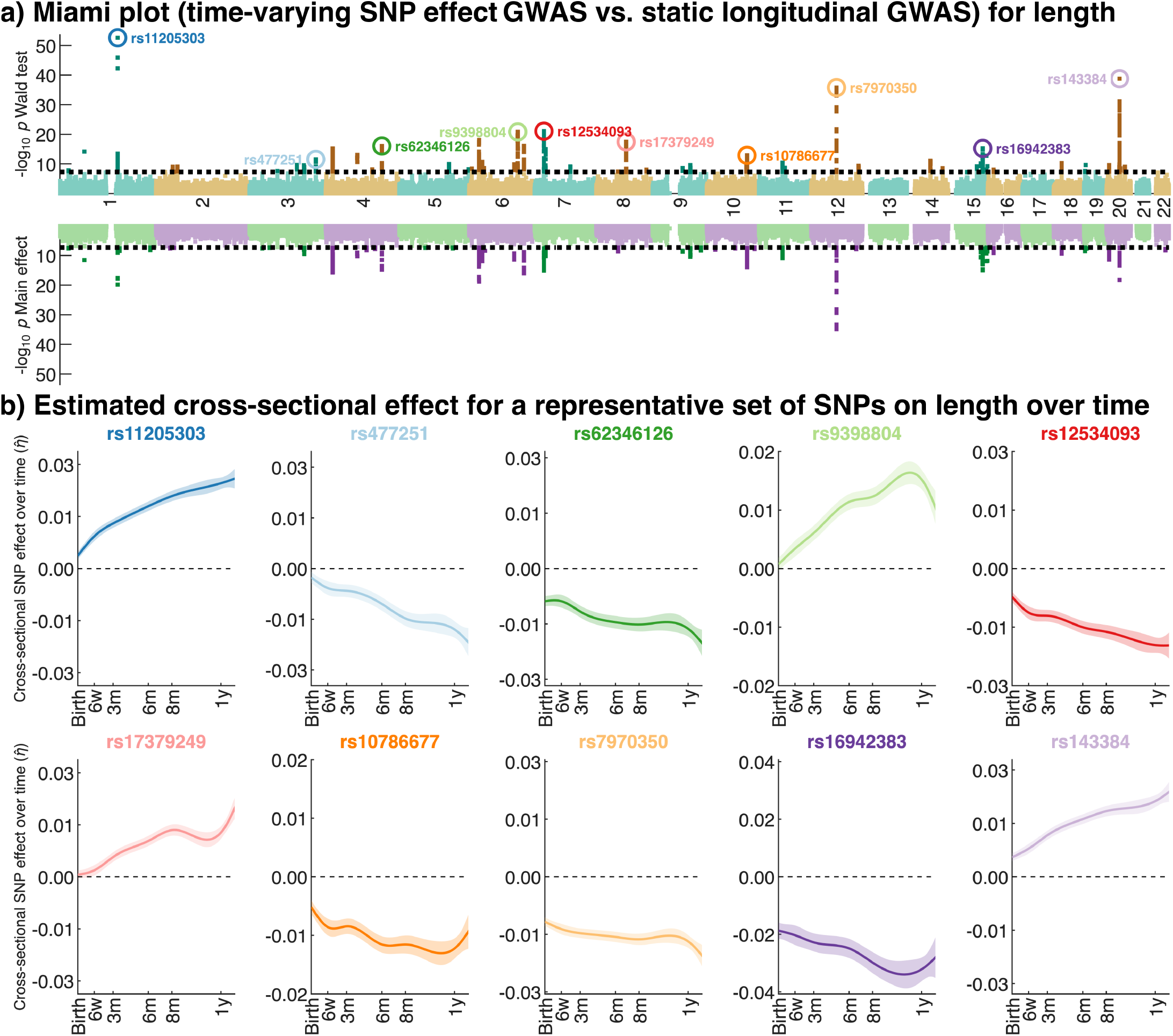
Miami plots comparing the omnibus effect of main and time-dependent effect of SNPs with the longitudinal main effect of SNPs on length of infants in the first year of life, and the estimated SNP effect over time for selected SNPs. **a)** the top part shows the −log_10_(*p*)-values for the Wald test while the lower part shows the −log_10_(*p*)-values for the longitudinal main effect; **b)** the solid lines show the cross-sectional effect of selected SNPs over time (SNPs marked in panel **a)**); the corresponding standard errors are shown by the band surrounding the solid lines.

**Figure 6:**
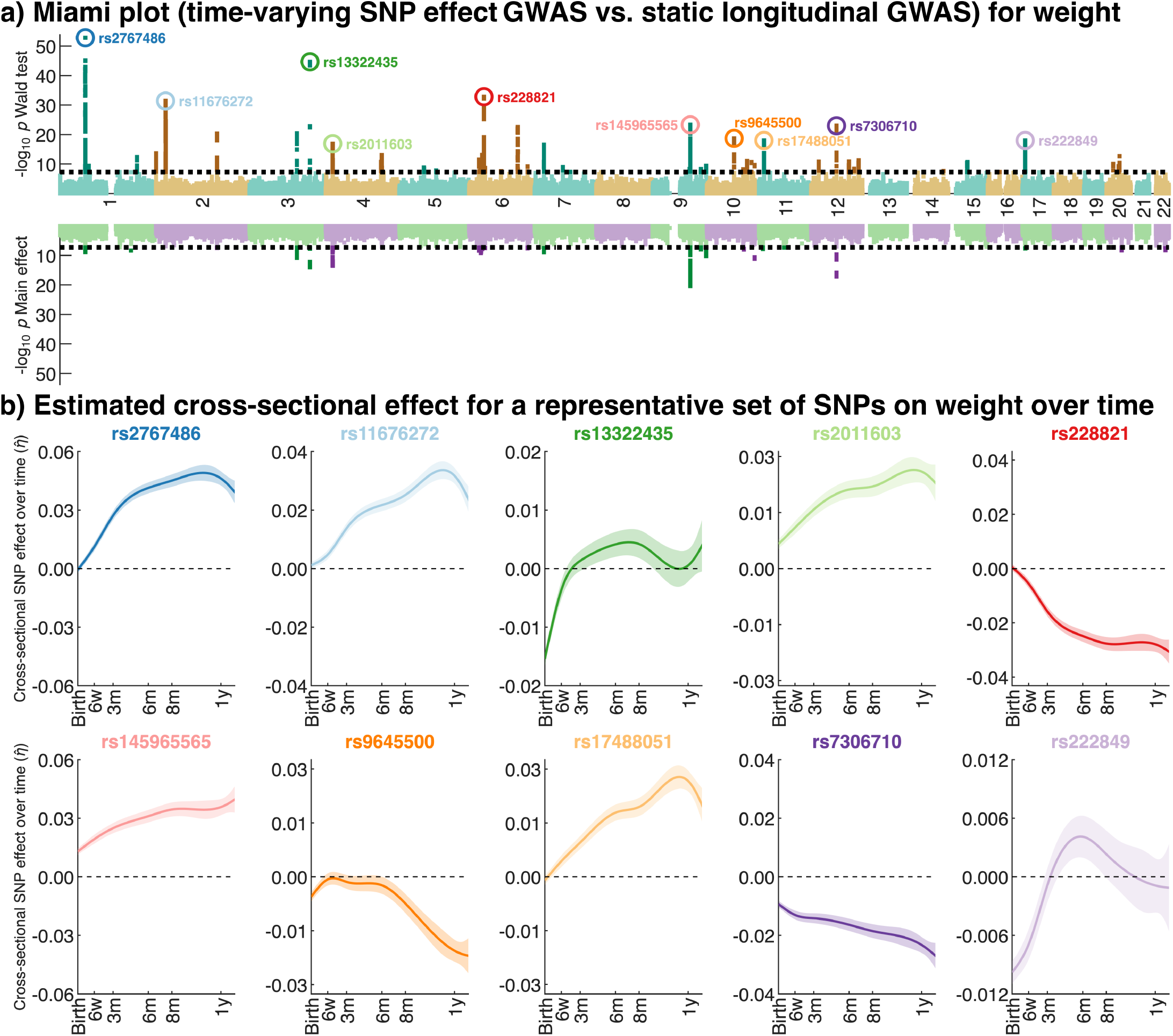
Miami plots comparing the omnibus effect of main and time-dependent effect of SNPs with the longitudinal main effect of SNPs on weight of infants in the first year of life, and the estimated SNP effect over time for selected SNPs. **a)** the top part shows the −log_10_(*p*)-values for the Wald test while the lower part shows the −log_10_(*p*)-values for the longitudinal main effect; **b)** the solid lines show the cross-sectional effect of selected SNPs over time (SNPs marked in panel **a)**); the corresponding standard errors are shown by the band surrounding the solid lines.

**Figure 7:**
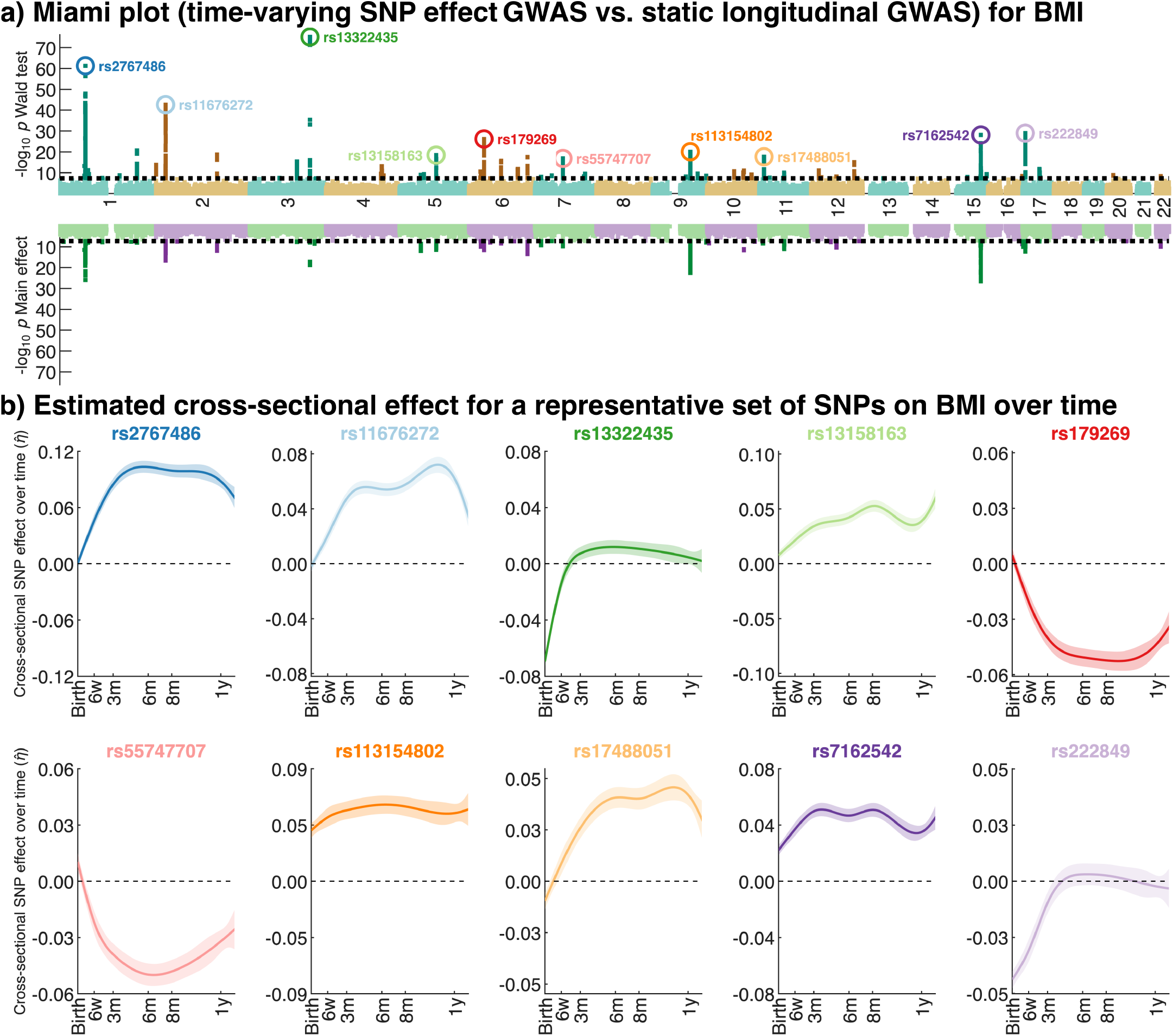
Miami plots comparing the omnibus effect of main and time-dependent effect of SNPs with the longitudinal main effect of SNPs on BMI of infants in the first year of life, and the estimated SNP effect over time for selected SNPs. **a)** the top part shows the −log_10_(*p*)-values for the Wald test while the lower part shows the −log_10_(*p*)-values for the longitudinal main effect; **b)** the solid lines show the cross-sectional effect of selected SNPs over time (SNPs marked in panel **a)**); the corresponding standard errors are shown by the band surrounding the solid lines.

## Discussion

We have adapted and extended our computationally fast linear mixed-effects modeling method FEMA for large-scale high-dimensional longitudinal datasets. The features of FEMA-Long include: i) modeling time-varying random effects using unstructured covariance, ii) incorporating splines for modeling non-linear effects of variables, iii) examining time-dependent effects using spline interactions, and iv) performing GWAS-like analyses by scaling FEMA to estimate marginal effect of millions of variables, including time-dependent effects. Our simulation results show that FEMA-Long can accurately model these parameters, while controlling for false positives, and provides exceptional computational performance, tens to thousands of times faster than a standard LME implementation, and at the same time having a much smaller carbon footprint. We have also demonstrated that FEMA-Long is scalable to scenarios with a large number of observations and outcome variables such as GWAS or whole-brain neuroimaging analyses.

As previously mentioned, random effects like heritability can vary significantly over time (3–8). Similarly, longitudinal data usually shows some form of time-varying correlation. Our application of FEMA-Long to length, weight, and BMI in infants during the first year of life revealed time-varying random effect covariances including heritability. These patterns also differed between similar phenotypes. Generally, increasing heritability for later timepoints (compared to birth) potentially indicates that the phenotypes at earlier timepoints might be more influenced from other sources like the maternal genotype (36,37). Importantly, it underscores the need to model these using unstructured covariance as these covariances cannot be well-approximated with simpler covariance patterns like compound symmetry or autoregressive models.

Additionally, when studying dynamic phenomenon such as development, it is essential to model time-dependent effects. Our time-dependent GWAS results revealed several SNPs showing time-dependent genetic effects. In contrast, the discovery of these SNPs was stunted when only examining the longitudinal main effect of SNPs, most notably for weight and BMI. Charting the effect of these SNPs over time revealed several interesting patterns including non-monotonic and curvilinear patterns. These results highlight the two-fold importance of modeling spline interactions for every SNP: first, allowing discovery of SNPs showing non-linear time-dependent effects; and second, ensuring more accurate predictions when using these estimated effect sizes (e.g., polygenic risk scores). Both these aspects cannot be captured by just the main effect of the SNPs and will likely be missed even with linear interaction.

The use of unstructured covariance has received relatively little attention in fields with high dimensional data. For example, within neuroimaging, there are no software packages for modeling unstructured covariances, besides AFNI’s *3dLME* and *3dLMEr* (38) which supports compound symmetry and order 1 autoregressive process (9). In genetics, prior work on longitudinal GWAS include: i) computing a rate of change between timepoints (39–41); ii) using standard LME packages (42,43); iii) fitting a growth curve-like model to the phenotype and performing GWAS on the model parameters (44,45); and iv) novel tools for longitudinal GWAS like fGWAS (46), GALLOP (47), trajGWAS (48), L-BRAT and RGMMAT (49), RVMMAT (50), and SPA_GRM_ (51). Despite the availability of these tools, they have one or more limitations including inability to: i) handle related individuals, ii) model additional sources of variances, iii) model linear and/or non-linear SNP interactions, iv) estimating effect sizes, and v) scale to a large samples and/or large-number of phenotypes. Importantly, it is rare to see a longitudinal GWAS study specifying a time-varying covariance pattern. The closest such recent approach (52) used a range of LMEs with linear or cubic smoothing splines, or cubic slopes for age, with and without a continuous AR(1) covariance pattern. However, this work still relied on collapsing repeated observations into a single measurement (followed by a meta-analysis) as well as removing related observations. In addition, the authors highlighted the significant computational burden when scaling to large cohorts.

FEMA-Long addresses all the above-mentioned limitations, providing a unified framework for performing large-scale analyses, modeling unstructured covariances, non-linear effects, and discovery of time-dependent effects, including time-dependent genetic effects. There are, however, some limitations to our work. First, the MoM estimator is not as efficient as maximum likelihood estimator. Second, currently FEMA-Long only supports continuous outcome variables, assuming approximately normally distributed errors. Third, for GWAS, we assume that the individual contribution of SNPs is small and unlikely to affect the covariance estimation of the random effect. Fourth, the computational time, especially for GWAS, could be further improved. Fifth, when performing the visit-wise estimation of the covariance pattern, for certain scenarios where the number of shared observations between the visits are too low, the method of moments estimation could be unstable. While FEMA and FEMA-Long are designed for large samples, we envision improving the stability of the covariance estimation by updating our method to perform a joint estimation of all parameters instead of visit-wise estimation; other possibilities include introducing a shrinkage or regularization on the estimation of the covariance parameters and/or enforcing smoothness during the estimation; additionally, we could interpolate the visit-wise estimated covariances, resulting in smoothly changing covariances, and further extending the scope of time-varying random effects. Future work will explore these and incorporate additional features like binning the phenotypes, computation of age-specific risk scores, extension to non-continuous phenotypes such as Bernoulli distribution, allowing statistical inference on the random effects estimates, using longitudinal trajectories for prediction, and the application of FEMA-Long to longitudinal complex trait phenotypes such as mental health.

To conclude, FEMA-Long is a powerful novel method that will enable researchers to apply large-scale LME models with unstructured covariances as well as discover time-dependent effects. The unified framework of FEMA-Long not only allows inclusion of subjects with a differing number of timepoints but also handles scenarios like sample relatedness and other sources of variance while providing a computationally fast and environmentally green solution for application to genomics, neuroimaging, and other fields. We expect FEMA-Long to facilitate discoveries in the growing number of large longitudinal datasets becoming available.

## Acknowledgements

The Norwegian Mother, Father and Child Cohort Study is supported by the Norwegian Ministry of Health and Care Services and the Ministry of Education and Research. We are grateful to all the participating families in Norway who take part in this on-going cohort study. We thank the Norwegian Institute of Public Health (NIPH) for generating high-quality genomic data. This research is part of the HARVEST collaboration, supported by the Research Council of Norway (#229624). We also thank the NORMENT Centre for providing genotype data, funded by the Research Council of Norway (#223273), South East Norway Health Authorities and Stiftelsen Kristian Gerhard Jebsen. We further thank the Center for Diabetes Research, the University of Bergen for providing genotype data and performing quality control and imputation of the data funded by the ERC AdG project SELECTionPREDISPOSED, Stiftelsen Kristian Gerhard Jebsen, Trond Mohn Foundation, the Research Council of Norway, the Novo Nordisk Foundation, the University of Bergen, and the Western Norway Health Authorities.

We acknowledge helpful suggestions from Drs. Gleda Kutrolli and Dominic Holland. We thank Dr. Elizabeth Corfield for leading the MoBaPsychGen quality control and imputation pipeline.

Some of the color schemes used for various figures were taken from ColorBrewer (https://colorbrewer2.org/). Some of the figures use the tight_subplot function (Pekka Kumpulainen (2023). tight_subplot(Nh, Nw, gap, marg_h, marg_w) (https://www.mathworks.com/matlabcentral/fileexchange/27991-tight_subplot-nh-nw-gap-marg_h-marg_w), MATLAB Central File Exchange. Retrieved April 5, 2023).

## Author contribution

**Conceptualization**: PP, NP, OF, AAS, TEN, OAA, AMD

**Data curation**: PP, EF, MV, PJ, VB, NRB, OF

**Formal analysis**: PP

**Funding acquisition**: PP, OAA, AMD

**Investigation**: PP, EF

**Methodology**: PP, NP, DP, WKT, OF, AAS, TEN, AMD

**Project administration**: OAA, AMD

**Resources**: IES, TLJ, OAA, AMD

**Software**: PP, DP, EF, DMS, AR, CCF, JK, OF, AAS, TEN, AMD

**Supervision**: OF, AAS, TEN, OAA, AMD

**Visualization**: PP, NP, PJ, TEN, AMD

**Writing – original draft**: PP

**Writing – review & editing**: PP, NP, DP, EF, MV, DMS, AR, PJ, IES, VB, NRB, CCF, CM, JK, RL, DJH, DM, SJ, PRN, TLJ, WKT, OF, AAS, TEN, OAA, AMD

## Data and code availability

FEMA-Long is available on GitHub at: https://github.com/cmig-research-group/cmig_tools. Data from the Norwegian Mother, Father and Child Cohort Study is managed by the Norwegian Institute of Public Health. Access requires approval from the Regional Committees for Medical and Health Research Ethics (REC), compliance with GDPR, and data owner approval. Participant consent does not allow individual-level data storage in repositories or journals. Researchers seeking access for replication must apply via www.helsedata.no. The synthetic data used for simulations can be generated using the code provided here: https://github.com/parekhpravesh/FEMA-Long.

