## Supplementary Materials for "FEMA-Long: Modeling unstructured covariances for discovery of time-dependent effects in large-scale longitudinal datasets"

**List of authors:**

Pravesh Parekh[1,2,*], Nadine Parker[1], Diliana Pecheva[2], Evgeniia Frei[1], Marc Vaudel[4,5,6], Diana M. Smith[7,8], Alison Rigby[9], Piotr Jahołkowski[1], Ida Elken Sønderby[1,10], Viktoria Birkenæs[1,11], Nora Refsum Bakken[1,11], Chun Chieh Fan[12], Carolina Makowski[13], Jakub Kopal[1], Robert Loughnan[2,14], Donald J. Hagler Jr.[2], Dennis van der Meer[1], Stefan Johansson[4,15], Pål Rasmus Njølstad[4,15], Terry L. Jernigan[3,7,13,16], Wesley K. Thompson[12], Oleksandr Frei[1,17], Alexey A. Shadrin[1], Thomas E. Nichols[18,19], Ole A. Andreassen[1], Anders M. Dale[2,20,21*]

**Affiliations:**

[1]Centre for Precision Psychiatry, Division of Mental Health and Addiction, University of Oslo and Oslo University Hospital, Oslo, Norway.

[2]Center for Multimodal Imaging and Genetics, J. Craig Venter Institute, La Jolla, CA, USA.

[3]Department of Radiology, School of Medicine, University of California San Diego, La Jolla, CA, USA.

[4]Mohn Center for Diabetes Precision Medicine, Department of Clinical Science, University of Bergen, Bergen, Norway.

[5]Computational Biology Unit, Department of Informatics, University of Bergen, Bergen, Norway.

[6]Department of Genetics and Bioinformatics, Norwegian Institute of Public Health, Bergen, Norway.

[7]Center for Human Development, University of California San Diego, La Jolla, CA, USA.

[8]Medical Scientist Training Program, University of California San Diego, La Jolla, CA, USA.

[9]Neuroscience Graduate Program, University of California San Diego, La Jolla, CA, USA.

[10]Department of Medical Genetics, Oslo University Hospital & University of Oslo, Oslo, Norway.

[11]PsychGen Center for Genetic Epidemiology and Mental Health, Norwegian Institute of Public Health, Oslo, Norway.

[12]Center for Population Neuroscience and Genetics, Laureate Institute for Brain Research, Tulsa, OK, USA.

[13]Department of Psychiatry, School of Medicine, University of California San Diego, La Jolla, CA, USA.

[14]Population Neuroscience and Genetics Lab, University of California San Diego, La Jolla, CA, USA.

[15]Department of Pediatrics and Adolescents, Haukeland University Hospital, Bergen, Norway.

[16]Department of Cognitive Science, University of California San Diego, La Jolla, CA, USA.

[17]Department of Pharmacy, Section for Pharmacology and Pharmaceutical Biosciences, University of Oslo, Oslo, Norway.

[18]Big Data Institute, Li Ka Shing Centre for Health Information and Discovery, Nuffield Department of Population Health, University of Oxford, Oxford, UK.

[19]Oxford Centre for Integrative Neuroimaging, FMRIB, Nuffield Department of Clinical Neurosciences, University of Oxford, Oxford, UK.

[20]University of California San Diego, La Jolla, CA, USA.

[21]University of Oslo, Oslo, Norway.

*Correspondence should be addressed to:

Dr. Pravesh Parekh: (PP)

Prof. Anders M. Dale: (AMD)

### Table of Contents


**Methods** - 1 -

**Identifiability of random effects parameters: variance** - 1 -

**Identifiability of random effects parameters: covariance** - 2 -

**Application of FEMA-Long** - 3 -

Phenotype for GWAS - 3 -

**Results** - 5 -

**Simulation 1: comparison of estimated parameters with ground truth** - 5 -

Fixed effects - 5 -

Family effect - 6 -

Subject effect - 7 -

**Simulation 1: comparison of estimated parameters with glmmTMB** - 8 -

Fixed effects - 8 -

Family effect - 9 -

Subject effect - 10 -

**Simulation 2: evaluation of false positives under the null** - 11 -

**Simulation 3: comparison of computational timing with glmmTMB** - 12 -

Single outcome variable - 12 -

Multiple outcome variable - 13 -

Single outcome variable with non-default settings - 16 -

Multiple outcome variables with non-default settings - 17 -

**Simulation 3: comparison of computational timing with lmer** - 20 -

Single outcome variable - 20 -

Multiple outcome variable - 21 -

**Simulation 3: comparison of carbon footprint with glmmTMB** - 24 -

Using default settings for glmmTMB - 24 -

Using non-default settings for glmmTMB - 27 -

**Simulation 3: comparison of carbon footprint with lmer** - 30 -

**Simulation 4: comparison of estimated parameters with ground truth** - 33 -

Sample size: 12,000 - 33 -

Sample size: 15,000 - 36 -

Sample size: 18,000 - 39 -

Sample size: 20,000 - 42 -

**Simulation 4: summary of comparison of estimated parameters** - 45 -

Comparison with ground truth - 45 -

Comparison with glmmTMB - 46 -

**Simulation 5: false positive calibration across sample sizes** - 47 -

Sample size: 12,000 - 47 -

Sample size 15,000 - 50 -

Sample size 18,000 - 53 -

Sample size 20,000 - 56 -

Examples of scenarios where the unstructured covariance resulted in inflated $\boldsymbol{p}$-values - 56 -

**Phenotype - 57 -**

Age - 57 -
Length - 58 -
Weight - 59 -
BMI - 60 -
Knot placement for creating spline basis functions - 61 -
Basis functions used for analysis - 62 -

**Q-Q plots - 63 -**

Length - 63 -
Weight - 63 -
BMI - 64 -

**Comparison between two-stage regression and full regression - 65 -**

Comparison of beta coefficients - 65 -
Comparison of standard error - 66 -
Comparison of *T* statistics - 67 -
Comparison of $\mathbf{-log10}\boldsymbol{p}$ values - 68 -
Comparison of Wald *F* - 69 -
Comparison of $\mathbf{-log10}$Wald $\boldsymbol{p}$ values - 70 -

**References - 71 -**

### Methods

#### Identifiability of random effects parameters: variance

**a)** Subjects with repeated measurements, nested within families

| Entry | Family | Subject | Visit | $y^{res}$ |
|---|---|---|---|---|
| $y_{F01S01r1}$ | F01 | S01 | r1 | −30 |
| $y_{F01S01r2}$ | F01 | S01 | r2 | −25 |
| $y_{F01S02r1}$ | F01 | S02 | r1 | −20 |
| $y_{F01S02r2}$ | F01 | S02 | r2 | −15 |
| $y_{F01S03r1}$ | F01 | S03 | r1 | −25 |
| $y_{F01S03r2}$ | F01 | S03 | r2 | −20 |
| $y_{F01S04r1}$ | F01 | S04 | r1 | −15 |
| $y_{F01S04r2}$ | F01 | S04 | r2 | −10 |
| $y_{F02S05r1}$ | F02 | S05 | r1 | 10 |
| $y_{F02S05r2}$ | F02 | S05 | r2 | 20 |
| $y_{F02S06r1}$ | F02 | S06 | r1 | 15 |
| $y_{F02S06r2}$ | F02 | S06 | r2 | 25 |
| $y_{F02S07r1}$ | F02 | S07 | r1 | 20 |
| $y_{F02S07r2}$ | F02 | S07 | r2 | 30 |
| $y_{F02S08r1}$ | F02 | S08 | r1 | 15 |
| $y_{F02S08r2}$ | F02 | S08 | r2 | 25 |

**c)** For within visit 1 variance, using entries from b), create a system of equations to solve: note that *S* and *E* are colinear

| Products of entries | $y_i^{res} \times y_{i'}^{res}$ | F | S | E | Term |
|---|---|---|---|---|---|
| $y_{F01S01r1} \times y_{F01S01r1}$ | −30 × −30 = 900 | 1 | 1 | 1 | F + S + E |
| $y_{F01S01r1} \times y_{F01S02r1}$ | −30 × −20 = 600 | 1 | 0 | 0 | F |
| $y_{F01S01r1} \times y_{F01S03r1}$ | −30 × −25 = 750 | 1 | 0 | 0 | F |
| $y_{F01S01r1} \times y_{F01S04r1}$ | −30 × −15 = 450 | 1 | 0 | 0 | F |
| $y_{F01S02r1} \times y_{F01S01r1}$ | −20 × −30 = 600 | 1 | 0 | 0 | F |
| $y_{F01S02r1} \times y_{F01S02r1}$ | −20 × −20 = 400 | 1 | 1 | 1 | F + S + E |
| $y_{F01S02r1} \times y_{F01S03r1}$ | −20 × −25 = 500 | 1 | 0 | 0 | F |
| $y_{F01S02r1} \times y_{F01S04r1}$ | −20 × −15 = 300 | 1 | 0 | 0 | F |
| $y_{F01S03r1} \times y_{F01S01r1}$ | −25 × −30 = 750 | 1 | 0 | 0 | F |
| $y_{F01S03r1} \times y_{F01S02r1}$ | −25 × −20 = 500 | 1 | 0 | 0 | F |
| $y_{F01S03r1} \times y_{F01S03r1}$ | −25 × −25 = 625 | 1 | 1 | 1 | F + S + E |
| $y_{F01S03r1} \times y_{F01S04r1}$ | −25 × −15 = 375 | 1 | 0 | 0 | F |
| $y_{F01S04r1} \times y_{F01S01r1}$ | −15 × −30 = 450 | 1 | 0 | 0 | F |
| $y_{F01S04r1} \times y_{F01S02r1}$ | −15 × −20 = 300 | 1 | 0 | 0 | F |
| $y_{F01S04r1} \times y_{F01S03r1}$ | −15 × −25 = 375 | 1 | 0 | 0 | F |
| $y_{F01S04r1} \times y_{F01S04r1}$ | −15 × −15 = 225 | 1 | 1 | 1 | F + S + E |
| $y_{F02S05r1} \times y_{F02S05r1}$ | 10 × 10 = 100 | 1 | 1 | 1 | F + S + E |
| $y_{F02S05r1} \times y_{F02S06r1}$ | 10 × 15 = 150 | 1 | 0 | 0 | F |
| ⋮ | ⋮ | ⋮ | ⋮ | ⋮ | ⋮ |
| $y_{F02S08r1} \times y_{F02S08r1}$ | 15 × 15 = 225 | 1 | 1 | 1 | F + S + E |

**b)** Create $Z_F Z_F^{\mathsf{T}}$, $Z_S Z_S^{\mathsf{T}}$, and error term $I$

| $i,i'$ | 1 | 2 | 3 | 4 | 5 | 6 | 7 | 8 | 9 | 10 | 11 | 12 | 13 | 14 | 15 | 16 |
|---|---|---|---|---|---|---|---|---|---|---|---|---|---|---|---|---|
| 1 | 1 | 1 | 1 | 1 | 1 | 1 | 1 | 1 | | | | | | | | |
| 2 | 1 | 1 | 1 | 1 | 1 | 1 | 1 | 1 | | | | | | | | |
| 3 | 1 | 1 | 1 | 1 | 1 | 1 | 1 | 1 | | | | | | | | |
| 4 | 1 | 1 | 1 | 1 | 1 | 1 | 1 | 1 | | | | | | | | |
| 5 | 1 | 1 | 1 | 1 | 1 | 1 | 1 | 1 | | | | | | | | |
| 6 | 1 | 1 | 1 | 1 | 1 | 1 | 1 | 1 | | | | | | | | |
| 7 | 1 | 1 | 1 | 1 | 1 | 1 | 1 | 1 | | | | | | | | |
| 8 | 1 | 1 | 1 | 1 | 1 | 1 | 1 | 1 | | | | | | | | |
| 9 | | | | | | | | | 1 | 1 | 1 | 1 | 1 | 1 | 1 | 1 |
| 10 | | | | | | | | | 1 | 1 | 1 | 1 | 1 | 1 | 1 | 1 |
| 11 | | | | | | | | | 1 | 1 | 1 | 1 | 1 | 1 | 1 | 1 |
| 12 | | | | | | | | | 1 | 1 | 1 | 1 | 1 | 1 | 1 | 1 |
| 13 | | | | | | | | | 1 | 1 | 1 | 1 | 1 | 1 | 1 | 1 |
| 14 | | | | | | | | | 1 | 1 | 1 | 1 | 1 | 1 | 1 | 1 |
| 15 | | | | | | | | | 1 | 1 | 1 | 1 | 1 | 1 | 1 | 1 |
| 16 | | | | | | | | | 1 | 1 | 1 | 1 | 1 | 1 | 1 | 1 |

| $i,i'$ | 1 | 2 | 3 | 4 | 5 | 6 | 7 | 8 | 9 | 10 | 11 | 12 | 13 | 14 | 15 | 16 |
|---|---|---|---|---|---|---|---|---|---|---|---|---|---|---|---|---|
| 1 | 1 | 1 | | | | | | | | | | | | | | |
| 2 | 1 | 1 | | | | | | | | | | | | | | |
| 3 | | | 1 | 1 | | | | | | | | | | | | |
| 4 | | | 1 | 1 | | | | | | | | | | | | |
| 5 | | | | | 1 | 1 | | | | | | | | | | |
| 6 | | | | | 1 | 1 | | | | | | | | | | |
| 7 | | | | | | | 1 | 1 | | | | | | | | |
| 8 | | | | | | | 1 | 1 | | | | | | | | |
| 9 | | | | | | | | | 1 | 1 | | | | | | |
| 10 | | | | | | | | | 1 | 1 | | | | | | |
| 11 | | | | | | | | | | | 1 | 1 | | | | |
| 12 | | | | | | | | | | | 1 | 1 | | | | |
| 13 | | | | | | | | | | | | | 1 | 1 | | |
| 14 | | | | | | | | | | | | | 1 | 1 | | |
| 15 | | | | | | | | | | | | | | | 1 | 1 |
| 16 | | | | | | | | | | | | | | | 1 | 1 |

| $i,i'$ | 1 | 2 | 3 | 4 | 5 | 6 | 7 | 8 | 9 | 10 | 11 | 12 | 13 | 14 | 15 | 16 |
|---|---|---|---|---|---|---|---|---|---|---|---|---|---|---|---|---|
| 1 | 1 | | | | | | | | | | | | | | | |
| 2 | | 1 | | | | | | | | | | | | | | |
| 3 | | | 1 | | | | | | | | | | | | | |
| 4 | | | | 1 | | | | | | | | | | | | |
| 5 | | | | | 1 | | | | | | | | | | | |
| 6 | | | | | | 1 | | | | | | | | | | |
| 7 | | | | | | | 1 | | | | | | | | | |
| 8 | | | | | | | | 1 | | | | | | | | |
| 9 | | | | | | | | | 1 | | | | | | | |
| 10 | | | | | | | | | | 1 | | | | | | |
| 11 | | | | | | | | | | | 1 | | | | | |
| 12 | | | | | | | | | | | | 1 | | | | |
| 13 | | | | | | | | | | | | | 1 | | | |
| 14 | | | | | | | | | | | | | | 1 | | |
| 15 | | | | | | | | | | | | | | | 1 | |
| 16 | | | | | | | | | | | | | | | | 1 |

**Figure S1**: Demonstration of random effects parameter identifiability for variance terms. a) consider a dataset with eight subjects across two families, each having two repeated measurements; b) dummy-coding families $Z_F$ and subjects$Z_S$, followed by creating the $Z_F Z_F^{\mathsf{T}}$, $Z_S Z_S^{\mathsf{T}}$ terms and the independent error term denoted by $I$; c) following equation 11 in the manuscript, create a system of equations to solve for visit 1: the terms $S$ and $E$ are colinear, and cannot be uniquely identified; therefore, these are estimated together. A similar set of equations can be constructed for visit 2.

### Identifiability of random effects parameters: covariance

**a)** Subjects with repeated measurements, nested within families

| Entry | Family | Subject | Visit | $y^{res}$ |
|---|---|---|---|---|
| $y_{F01S01r1}$ | F01 | S01 | r1 | −30 |
| $y_{F01S01r2}$ | F01 | S01 | r2 | −25 |
| $y_{F01S02r1}$ | F01 | S02 | r1 | −20 |
| $y_{F01S02r2}$ | F01 | S02 | r2 | −15 |
| $y_{F01S03r1}$ | F01 | S03 | r1 | −25 |
| $y_{F01S03r2}$ | F01 | S03 | r2 | −20 |
| $y_{F01S04r1}$ | F01 | S04 | r1 | −15 |
| $y_{F01S04r2}$ | F01 | S04 | r2 | −10 |
| $y_{F02S05r1}$ | F02 | S05 | r1 | 10 |
| $y_{F02S05r2}$ | F02 | S05 | r2 | 20 |
| $y_{F02S06r1}$ | F02 | S06 | r1 | 15 |
| $y_{F02S06r2}$ | F02 | S06 | r2 | 25 |
| $y_{F02S07r1}$ | F02 | S07 | r1 | 20 |
| $y_{F02S07r2}$ | F02 | S07 | r2 | 30 |
| $y_{F02S08r1}$ | F02 | S08 | r1 | 15 |
| $y_{F02S08r2}$ | F02 | S08 | r2 | 25 |

**c)** For visit 1 – visit 2 covariance, using entries from b), create a system of equations to solve: note that $E$ is unidentifiable

| Products of entries | $y_i^{res} \times y_{i'}^{res}$ | F | S | E | Term |
|---|---|---|---|---|---|
| $y_{F01S01r1} \times y_{F01S01r2}$ | −30 ×−25 = 750 | 1 | 1 | 0 | F + S |
| $y_{F01S01r1} \times y_{F01S02r2}$ | −30 × −15 = 450 | 1 | 0 | 0 | F |
| $y_{F01S01r1} \times y_{F01S03r2}$ | −30 × −20 = 600 | 1 | 0 | 0 | F |
| $y_{F01S01r1} \times y_{F01S04r2}$ | −30 × −10 = 300 | 1 | 0 | 0 | F |
| $y_{F01S02r1} \times y_{F01S01r2}$ | −20 × −25 = 500 | 1 | 0 | 0 | F |
| $y_{F01S02r1} \times y_{F01S02r2}$ | −20 × −15 = 300 | 1 | 1 | 0 | F + S |
| $y_{F01S02r1} \times y_{F01S03r2}$ | −20 × −20 = 400 | 1 | 0 | 0 | F |
| $y_{F01S02r1} \times y_{F01S04r2}$ | −20 × −10 = 200 | 1 | 0 | 0 | F |
| $y_{F01S03r1} \times y_{F01S01r2}$ | −25 × −25 = 625 | 1 | 0 | 0 | F |
| $y_{F01S03r1} \times y_{F01S02r2}$ | −25 × −15 = 375 | 1 | 0 | 0 | F |
| $y_{F01S03r1} \times y_{F01S03r2}$ | −25 × −20 = 500 | 1 | 1 | 0 | F + S |
| $y_{F01S03r1} \times y_{F01S04r2}$ | −25 × −10 = 250 | 1 | 0 | 0 | F |
| $y_{F01S04r1} \times y_{F01S01r2}$ | −15 × −25 = 375 | 1 | 0 | 0 | F |
| $y_{F01S04r1} \times y_{F01S02r2}$ | −15 × −15 = 225 | 1 | 0 | 0 | F |
| $y_{F01S04r1} \times y_{F01S03r2}$ | −15 × −20 = 300 | 1 | 0 | 0 | F |
| $y_{F01S04r1} \times y_{F01S04r2}$ | −15 × −10 = 150 | 1 | 1 | 0 | F + S |
| $y_{F02S05r1} \times y_{F02S05r2}$ | 10 × 20 = 200 | 1 | 1 | 0 | F + S |
| $y_{F02S05r1} \times y_{F02S06r2}$ | 10 × 25 = 250 | 1 | 0 | 0 | F |
| ⋮ | ⋮ | ⋮ | ⋮ | ⋮ | ⋮ |
| $y_{F02S08r1} \times y_{F02S08r2}$ | 15 × 25 = 375 | 1 | 1 | 0 | F + S |

**b)** Create $Z_F Z_F^{\top}$, $Z_S Z_S^{\top}$, and error term $I$

| $i,i'$ | 1 | 2 | 3 | 4 | 5 | 6 | 7 | 8 | 9 | 10 | 11 | 12 | 13 | 14 | 15 | 16 |
|---|---|---|---|---|---|---|---|---|---|---|---|---|---|---|---|---|
| 1 | 1 | 1 | 1 | 1 | 1 | 1 | 1 | 1 | | | | | | | | |
| 2 | 1 | 1 | 1 | 1 | 1 | 1 | 1 | 1 | | | | | | | | |
| 3 | 1 | 1 | 1 | 1 | 1 | 1 | 1 | 1 | | | | | | | | |
| 4 | 1 | 1 | 1 | 1 | 1 | 1 | 1 | 1 | | | | | | | | |
| 5 | 1 | 1 | 1 | 1 | 1 | 1 | 1 | 1 | | | | | | | | |
| 6 | 1 | 1 | 1 | 1 | 1 | 1 | 1 | 1 | | | | | | | | |
| 7 | 1 | 1 | 1 | 1 | 1 | 1 | 1 | 1 | | | | | | | | |
| 8 | 1 | 1 | 1 | 1 | 1 | 1 | 1 | 1 | | | | | | | | |
| 9 | | | | | | | | | 1 | 1 | 1 | 1 | 1 | 1 | 1 | 1 |
| 10 | | | | | | | | | 1 | 1 | 1 | 1 | 1 | 1 | 1 | 1 |
| 11 | | | | | | | | | 1 | 1 | 1 | 1 | 1 | 1 | 1 | 1 |
| 12 | | | | | | | | | 1 | 1 | 1 | 1 | 1 | 1 | 1 | 1 |
| 13 | | | | | | | | | 1 | 1 | 1 | 1 | 1 | 1 | 1 | 1 |
| 14 | | | | | | | | | 1 | 1 | 1 | 1 | 1 | 1 | 1 | 1 |
| 15 | | | | | | | | | 1 | 1 | 1 | 1 | 1 | 1 | 1 | 1 |
| 16 | | | | | | | | | 1 | 1 | 1 | 1 | 1 | 1 | 1 | 1 |

| $i,i'$ | 1 | 2 | 3 | 4 | 5 | 6 | 7 | 8 | 9 | 10 | 11 | 12 | 13 | 14 | 15 | 16 |
|---|---|---|---|---|---|---|---|---|---|---|---|---|---|---|---|---|
| 1 | 1 | 1 | | | | | | | | | | | | | | |
| 2 | 1 | 1 | | | | | | | | | | | | | | |
| 3 | | | 1 | 1 | | | | | | | | | | | | |
| 4 | | | 1 | 1 | | | | | | | | | | | | |
| 5 | | | | | 1 | 1 | | | | | | | | | | |
| 6 | | | | | 1 | 1 | | | | | | | | | | |
| 7 | | | | | | | 1 | 1 | | | | | | | | |
| 8 | | | | | | | 1 | 1 | | | | | | | | |
| 9 | | | | | | | | | 1 | 1 | | | | | | |
| 10 | | | | | | | | | 1 | 1 | | | | | | |
| 11 | | | | | | | | | | | 1 | 1 | | | | |
| 12 | | | | | | | | | | | 1 | 1 | | | | |
| 13 | | | | | | | | | | | | | 1 | 1 | | |
| 14 | | | | | | | | | | | | | 1 | 1 | | |
| 15 | | | | | | | | | | | | | | | 1 | 1 |
| 16 | | | | | | | | | | | | | | | 1 | 1 |

| $i,i'$ | 1 | 2 | 3 | 4 | 5 | 6 | 7 | 8 | 9 | 10 | 11 | 12 | 13 | 14 | 15 | 16 |
|---|---|---|---|---|---|---|---|---|---|---|---|---|---|---|---|---|
| 1 | 1 | | | | | | | | | | | | | | | |
| 2 | | 1 | | | | | | | | | | | | | | |
| 3 | | | 1 | | | | | | | | | | | | | |
| 4 | | | | 1 | | | | | | | | | | | | |
| 5 | | | | | 1 | | | | | | | | | | | |
| 6 | | | | | | 1 | | | | | | | | | | |
| 7 | | | | | | | 1 | | | | | | | | | |
| 8 | | | | | | | | 1 | | | | | | | | |
| 9 | | | | | | | | | 1 | | | | | | | |
| 10 | | | | | | | | | | 1 | | | | | | |
| 11 | | | | | | | | | | | 1 | | | | | |
| 12 | | | | | | | | | | | | 1 | | | | |
| 13 | | | | | | | | | | | | | 1 | | | |
| 14 | | | | | | | | | | | | | | 1 | | |
| 15 | | | | | | | | | | | | | | | 1 | |
| 16 | | | | | | | | | | | | | | | | 1 |

**Figure S2**: Demonstration of random effects parameter identifiability for covariance terms. a) consider a dataset with eight subjects across two families, each having two repeated measurements; b) dummy-coding families $Z_F$ and subjects $Z_S$, followed by creating the $Z_F Z_F^{\top}$, $Z_S Z_S^{\top}$ terms and the independent error term denoted by $I$; c) following equation 11 in the manuscript, create a system of equations to solve for visit 1 – visit 2: the term $E$ becomes unidentifiable in this case.

#### Application of FEMA-Long

##### Phenotype for GWAS

As mentioned in the main manuscript, the phenotype of interest was the length, weight, and BMI of infants at six time points during the first year of life: birth, length at six weeks, three months, six months, eight months, and twelve months. The starting sample size was data on 113,342 infants. The measurements were derived from a combination of Medical Birth Registry of Norway (MBRN) and questionnaires filled by the mothers six and eighteen months after birth. Within the questionnaire filled by mothers when their child was six months old, the questions included a record of the length and weight of the child at birth, six weeks, three months, and six months. For the latter three, the mother was asked to refer to the child's health card. Within the questionnaire filled by mothers when their child was eighteen months old, the questions included the length and weight of the child at eight months, around one year, and between fifteen to eighteen months.

*Consolidating measurements*

First, we ensured that we had information regarding birth year for every child. Additionally, we ensured that there were no duplicated records. Then, we removed the data on 214 individuals where the sex of the child was unknown. Next, we removed 646 individuals where the pregnancy duration was less than 22 weeks or more than 44 weeks and 7,343 individuals where the gestational period was less than 37 weeks. Next, using the MBRN records, we removed the data on 4,582 infants with congenital malformations; we additionally checked the records for chromosomal abnormalities and Down's syndrome but at this point no such records were in the dataset.

Next, for each timepoint, we ensured that the age at that timepoint was greater than the mode of the age at its preceding timepoint and less than the mode of the age at the next timepoint. For example, age of infants at three months measurement ought to be greater than the mode of age at six weeks and less than the mode of age at six months. For the measurement at twelve months, we used the upper cut-off of 425 days. This resulted in removing the data of 1,127 individuals (six weeks), 1,009 individuals (three months), 1,019 individuals (six months), 2,223 individuals (eight months), and 1,286 individuals (twelve months). Next, for any individual where the birth length or weight was missing, we replaced the measurement with the questionnaire data; for 1,611 individuals, birth length or weight record was missing in both the MBRN and the questionnaire data – these entries were removed. Finally, we removed the data on fifteen individuals where the six-week age was less than seven days, and the weight gain (since birth) was more than 400 grams.

*Examining growth charts*

We calculated sex-specific percentiles of length and weight measurements at each timepoint. Specifically, we removed any observation that was less than $0.5^{th}$ percentile or greater than $99.5^{th}$ percentile at each timepoint. After this step, we were left with the data on 98,946 individuals. Then, we examined growth curves and flagged data points as outliers using a strategy described in (1): for each successive pair of timepoints (birth and six weeks, six weeks and three months, and so forth), we computed the $\log_2$ of the ratios (for both length and weight; for example, $\log_2\left(\frac{3m}{6w}\right)$). Using this value, we computed the median and the 0.0013 and 0.9986 quantiles of the data (normal cumulative distribution function at values -3 and 3 respectively). Using these three values, we derived a signed ratio which identified whether the timepoint value was an outlier (see (1) for more details). We ran this growth curve inspection stage only once (rather than iteratively) and marked these outliers. At this point, we also calculated the BMI for the infants and merged the dataset with the subjects who had been genotyped and who's genetic data had passed quality control (2). We removed the records of three individuals where there was a mismatch between genotyping sex and the information retrieved from MBRN/questionnaire. The steps till this point were carried out in Python 3.9.6 using libraries Pandas 2.2.3 (3), NumPy 2.0.2 (4), and SciPy 1.13.1 (5).

*Final quality check*

At this point, we had data on 68,514 unique individuals and a total of 411,084 observations. Out of these, we removed 93,024 observations where the age was missing, followed by removing 16,524 observations where either length or weight information was missing. These 301,536 observations with full information were subjected to a few additional quality check steps. We identified 1,065 individuals where there was one or more duplicated age; out of these, there were 244 observations which had different length or weight values at the same age (which we removed). There were 66 observations where there was a discrepancy between the timepoint and the ordering of age (for example, age at eight months timepoint was smaller than the age at six months timepoint).

In the remaining dataset of 300,283 observations, we found 109 individuals where there was one instance of duplicated BMI value (i.e., observations with the same value of length and weight but different age value) – since it was not possible to ascertain which of these observations were artifactual, we removed 218 observations of duplicated BMI. Next, we checked for scenarios where the length of the individual decreased with age. We found 114 individuals where this was so; for these individuals, we removed 228 measurements where length had decreased. Finally, we removed the data for 85 monozygotic twin pairs (one of the twins). At the end of this process, we were left with 299,447 complete length, weight, and BMI observations on 68,273 infants between birth and first year of life. The final sample size for GWAS was 68,273 infants with 299,447 observations having complete data on length, weight, and BMI (i.e., each of the 68,273 subjects had all three measurements).

### Results

#### Simulation 1: comparison of estimated parameters with ground truth

##### Fixed effects

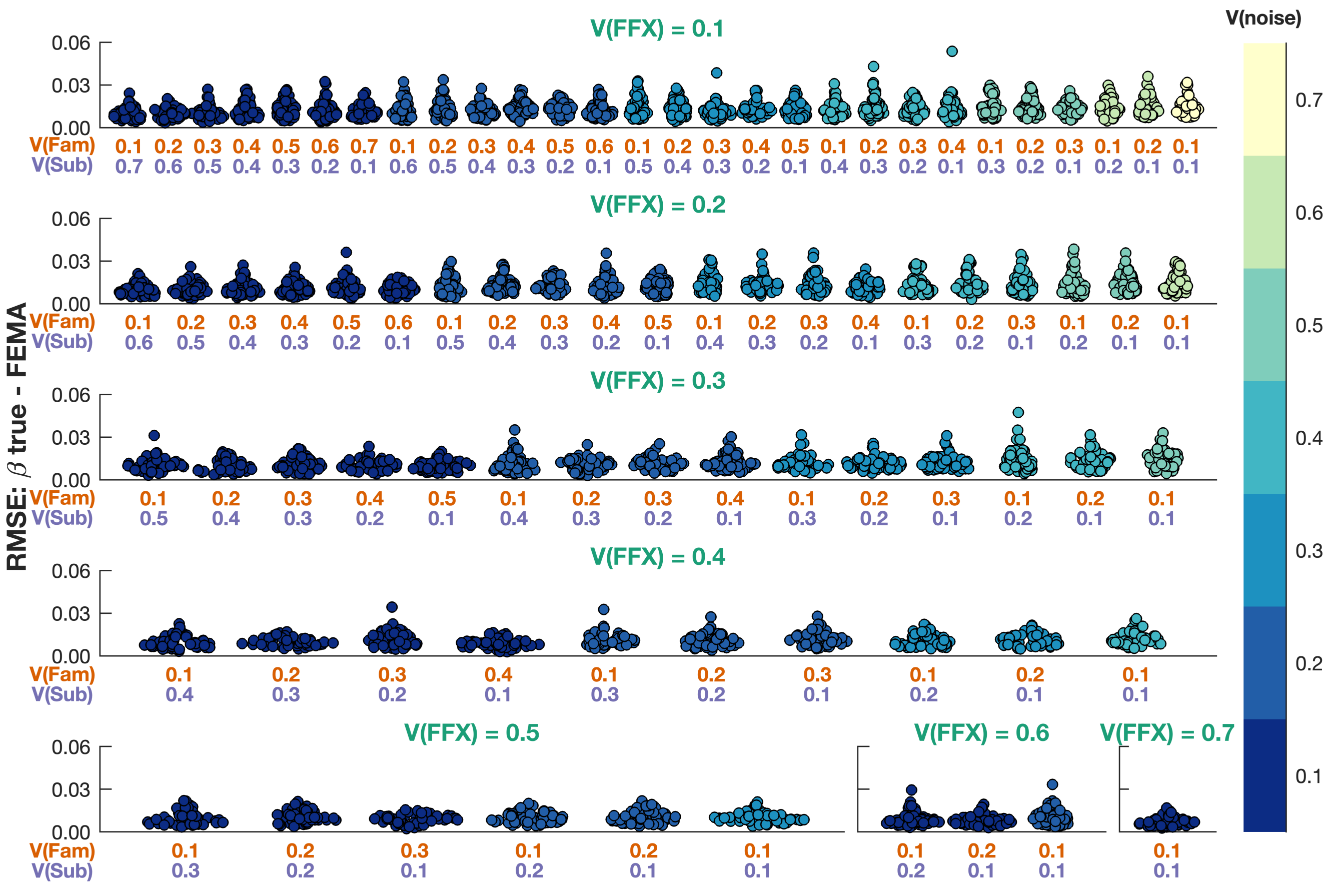


**Figure S3: Comparison of estimated beta coefficients from FEMA with ground truth for the fixed effect**. Each panel shows a simulation condition with the amount of variance in the phenotypes explained by the fixed effect V(FFX) shown on the top and the amounts of variances explained by family and subject effects V(Fam) and V(Sub) labeled on the *x*-axis. Each point shows the root mean squared error (RMSE) between the simulated ground truth and the estimates from FEMA, repeated 50 times for each simulation scenario, color-coded by the amount of noise in the phenotype.

#### Family effect

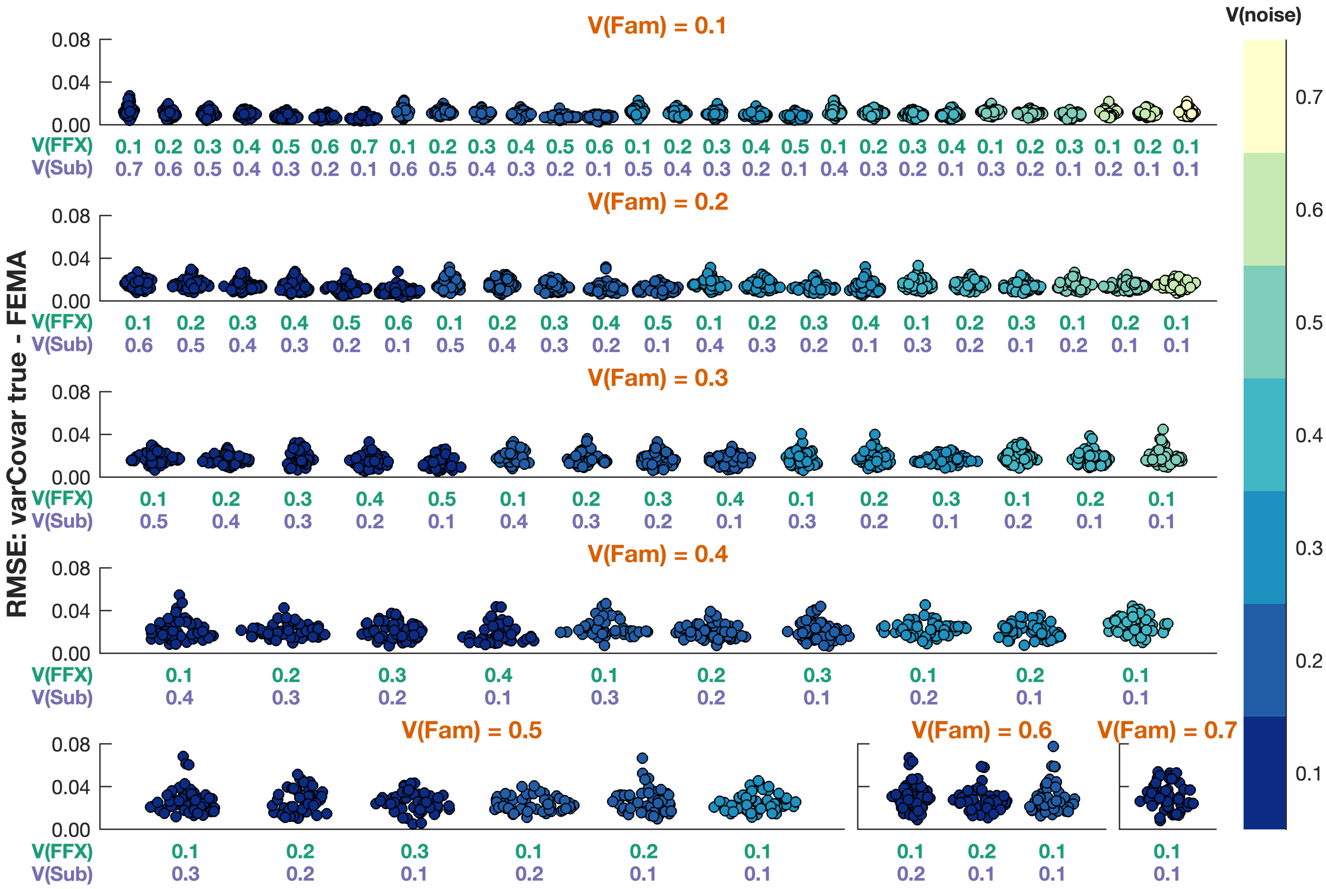


**Figure S4: Comparison of estimated variance covariance matrix coefficients from FEMA with ground truth for family effect**. Each panel shows a simulation condition with the amount of variance in the phenotypes explained by the family effect V(Fam) shown on the top and the amounts of variances explained by fixed effects and subject effects V(FFX) and V(Sub) labeled on the *x*-axis. Each point shows the root mean squared error (RMSE) between the simulated ground truth and the estimates from FEMA, repeated 50 times for each simulation scenario, color-coded by the amount of noise in the phenotype.

#### Subject effect

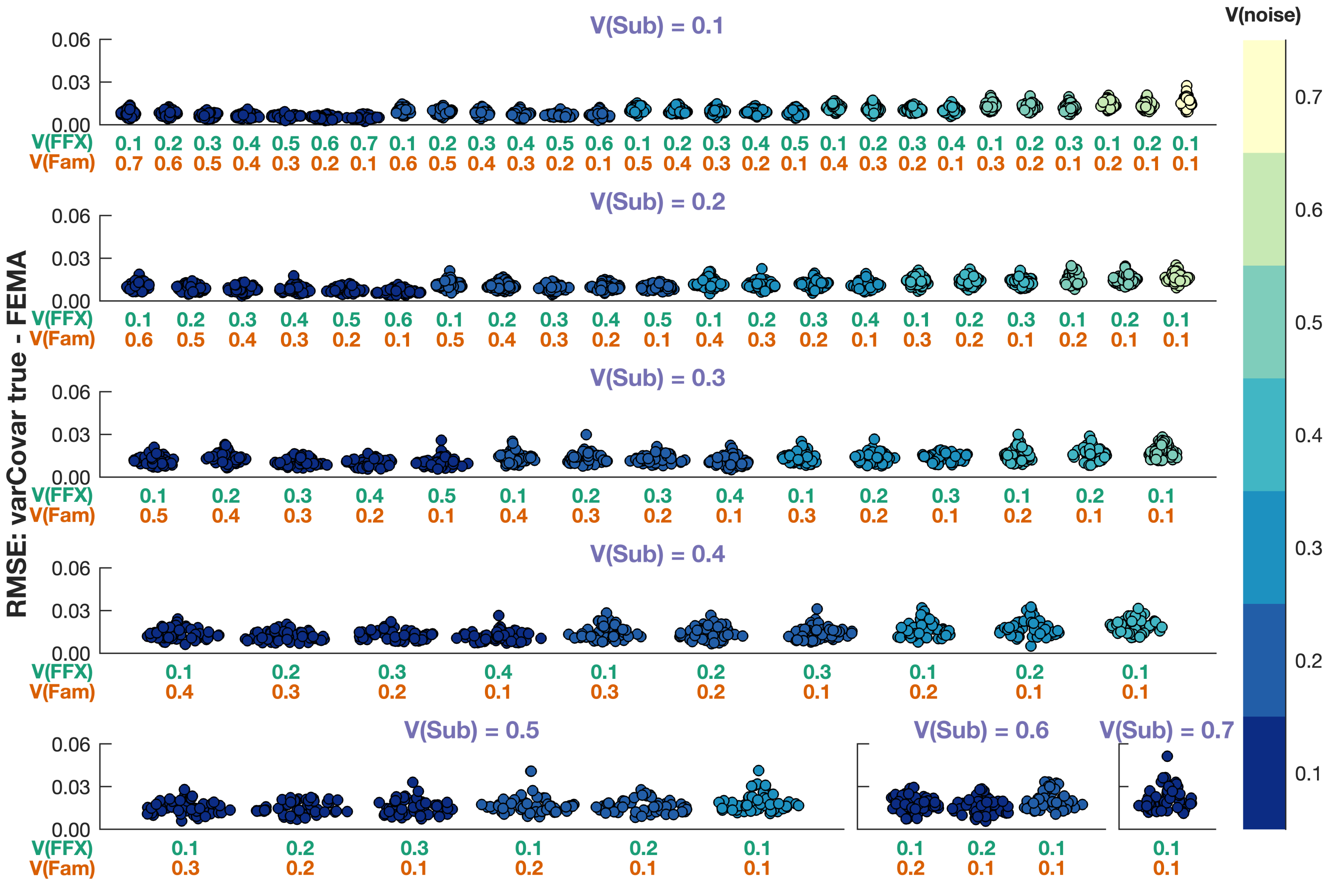


**Figure S5: Comparison of estimated variance covariance matrix coefficients from FEMA with ground truth for subject effect**. Each panel shows a simulation condition with the amount of variance in the phenotypes explained by the subject effect V(Sub) shown on the top and the amounts of variances explained by fixed effects and family effects V(FFX) and V(Fam) labeled on the *x*-axis. Each point shows the root mean squared error (RMSE) between the simulated ground truth and the estimates from FEMA, repeated 50 times for each simulation scenario, color-coded by the amount of noise in the phenotype.

### Simulation 1: comparison of estimated parameters with glmmTMB

#### Fixed effects

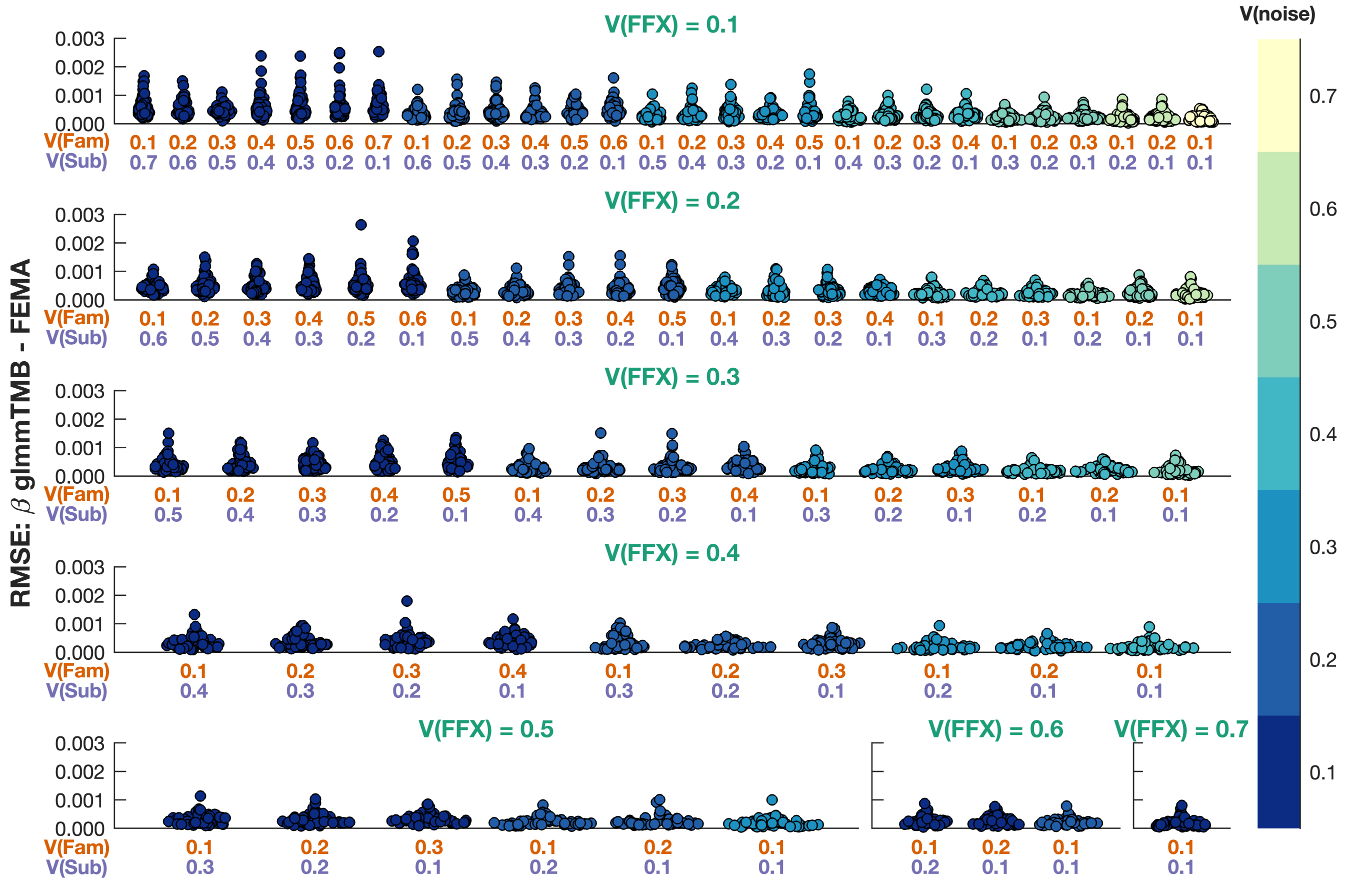


**Figure S6: Comparison of estimated beta coefficients from FEMA with estimates from glmmTMB for the fixed effects**. Each panel shows a simulation condition with the amount of variance in the phenotypes explained by the fixed effects V(FFX) shown on the top and the amounts of variances explained by family effects and subject effects V(Fam) and V(Sub) labeled on the *x*-axis. Each point shows the root mean squared error (RMSE) between the estimates from glmmTMB and the estimates from FEMA, repeated 50 times for each simulation scenario, color-coded by the amount of noise in the phenotype.

#### Family effect

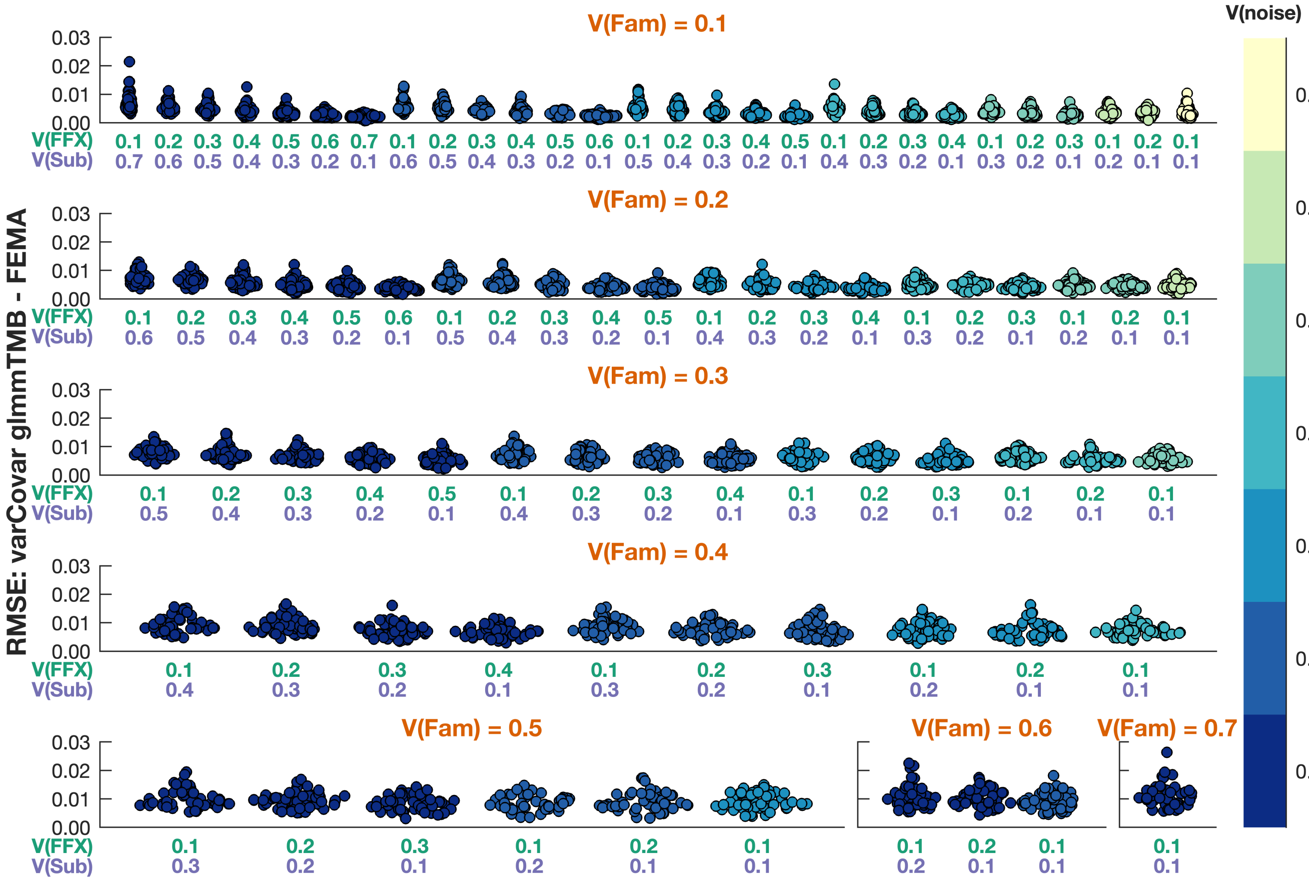


**Figure S7: Comparison of estimated variance covariance matrix coefficients from FEMA with estimates from glmmTMB for family effect**. Each panel shows a simulation condition with the amount of variance in the phenotypes explained by the family effects V(Fam) shown on the top and the amounts of variances explained by fixed effects and subject effects V(FFX) and V(Sub) labeled on the *x*-axis. Each point shows the root mean squared error (RMSE) between the estimates from glmmTMB and the estimates from FEMA, repeated 50 times for each simulation scenario, color-coded by the amount of noise in the phenotype.

#### Subject effect

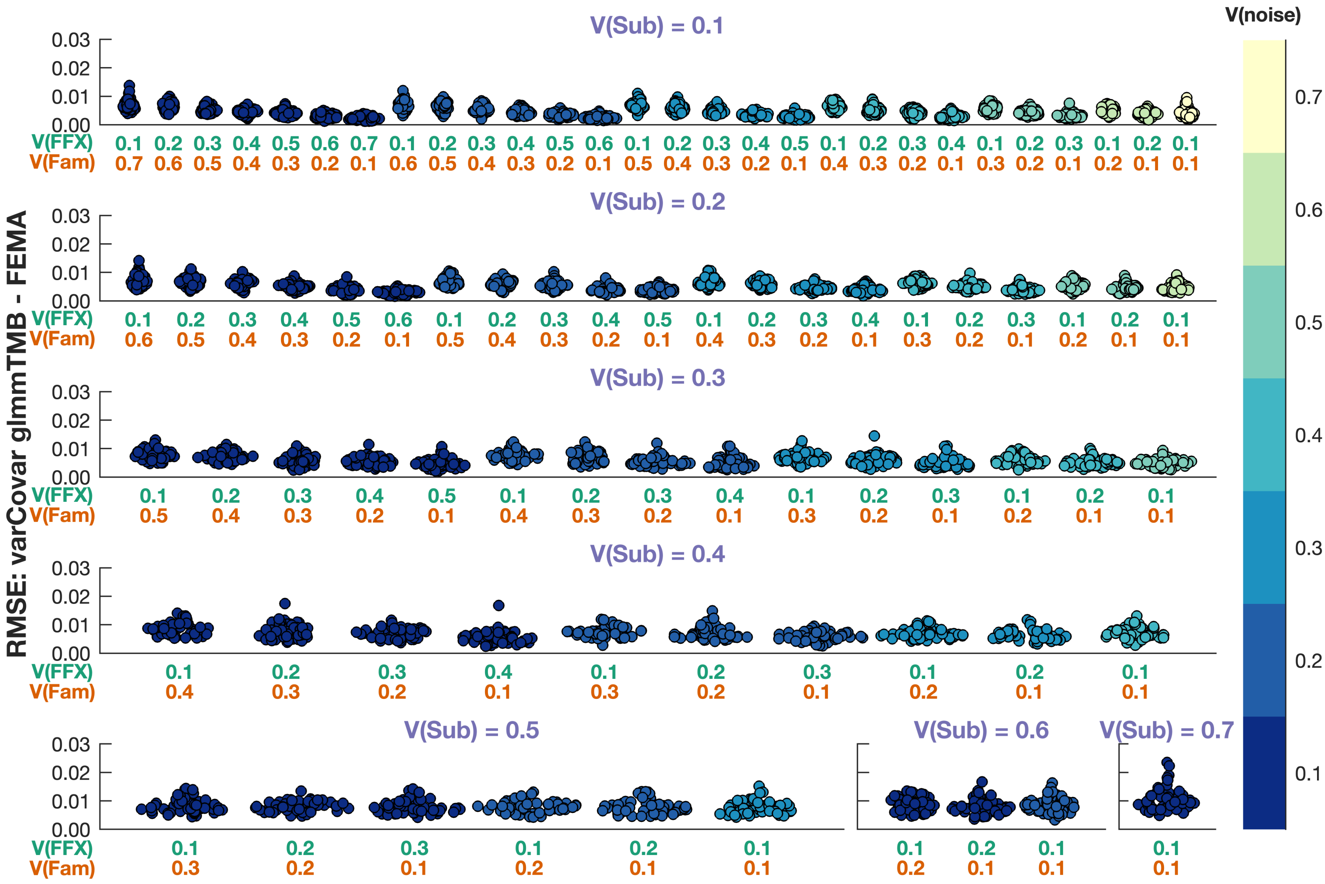


**Figure S8: Comparison of estimated variance covariance matrix coefficients from FEMA with estimates from glmmTMB for subject effect**. Each panel shows a simulation condition with the amount of variance in the phenotypes explained by the subject effects V(Sub) shown on the top and the amounts of variances explained by fixed effects and family effects V(FFX) and V(Fam) labeled on the *x*-axis. Each point shows the root mean squared error (RMSE) between the estimates from glmmTMB and the estimates from FEMA, repeated 50 times for each simulation scenario, color-coded by the amount of noise in the phenotype.

#### Simulation 2: evaluation of false positives under the null

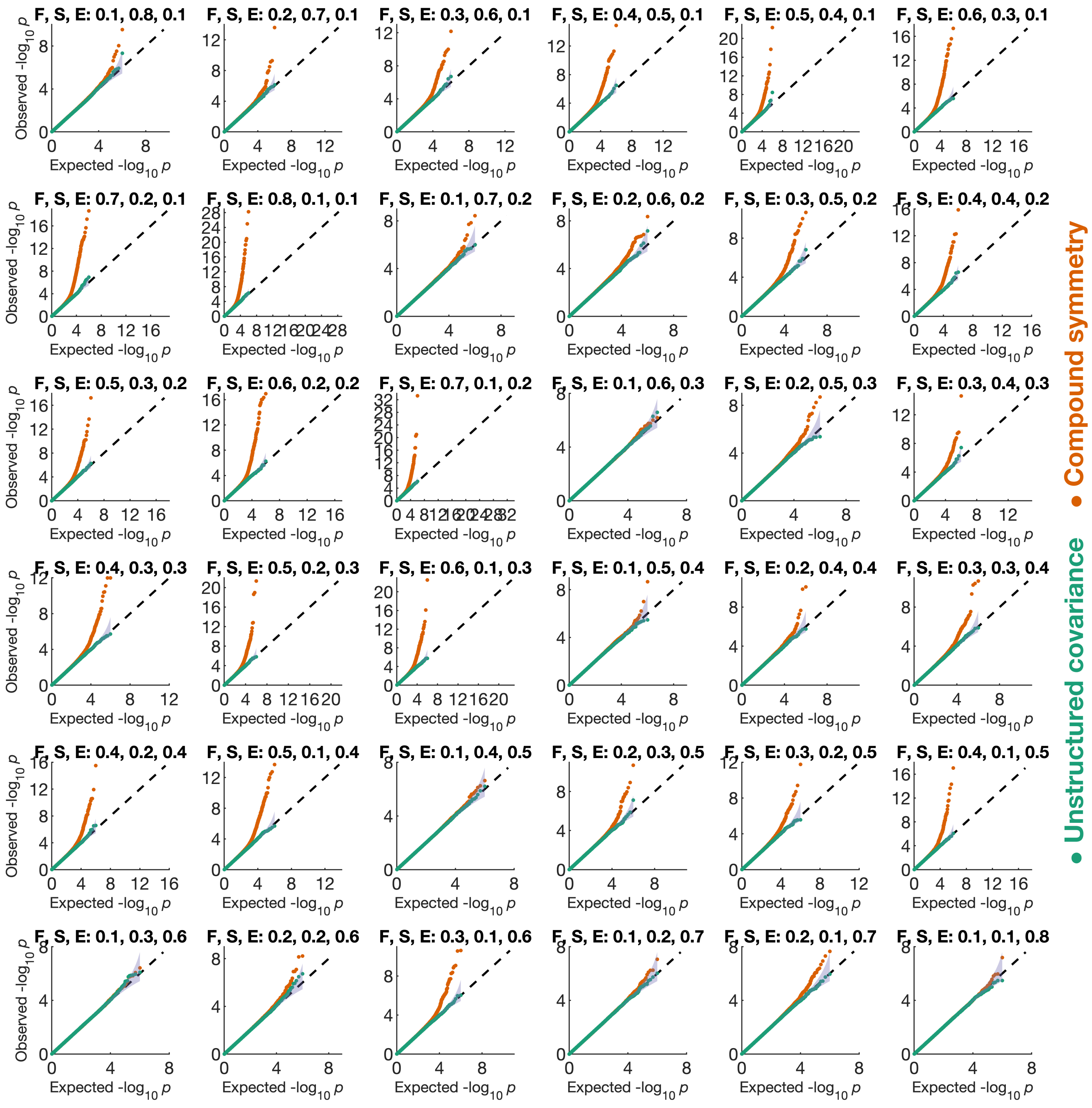


**Figure S9: Q-Q plots showing the distribution of $-\log_{10}(p)$ values under different simulation scenarios**. The simulation setting is indicated on the top of each Q-Q plot indicating the amounts of variances (in the phenotype) explained by family (F), subject (S), and noise (E); the $x$-axes indicate the expected $-\log_{10}(p)$ values under the null hypothesis while the $y$-axes show the observed $-\log_{10}(p)$ values across 1000 repeats, 100 $X$ variables, and 10 $y$ variables. The purple filled area indicates the 95% confidence interval based on inverse beta distribution.

### Simulation 3: comparison of computational timing with glmmTMB

#### Single outcome variable

**Table S1: Comparison of time taken (in seconds) by glmmTMB and FEMA for a single outcome variable**

| Number of observations | Median time glmmTMB | Median time FEMA (serial) | FEMA speed-up factor (serial) |
|---|---|---|---|
| 1000 | 9.76 | 0.14 | 70.80 |
| 1668 | 15.70 | 0.16 | 101.13 |
| 2783 | 26.51 | 0.25 | 105.67 |
| 4642 | 43.20 | 0.51 | 84.49 |
| 7743 | 72.52 | 0.86 | 83.87 |
| 12915 | 121.42 | 1.60 | 75.94 |
| 21544 | 202.02 | 3.10 | 65.11 |
| 35938 | 348.22 | 6.42 | 54.27 |
| 59948 | 605.86 | 14.21 | 42.63 |
| 100000 | 1013.11 | 34.80 | 29.11 |

#### Multiple outcome variable

**Table S2: Comparison of time taken (in seconds) by glmmTMB and FEMA for multiple outcome variables.** The glmmTMB timings are extrapolated from the time taken for a single outcome variable

| No. of observations | No. of *y* vars. | Median time glmmTMB | Median time FEMA (serial) | Median time FEMA (parallel) | FEMA speed-up factor (serial) | FEMA speed-up factor (parallel) |
|---|---|---|---|---|---|---|
| 1000 | 3 | 29.27 | 0.13 | 0.14 | 233.65 | 203.42 |
| 1668 | 3 | 47.09 | 0.19 | 0.18 | 253.57 | 265.22 |
| 2783 | 3 | 79.54 | 0.28 | 0.28 | 281.49 | 280.55 |
| 4642 | 3 | 129.61 | 0.59 | 0.60 | 219.75 | 215.68 |
| 7743 | 3 | 217.55 | 0.98 | 0.97 | 222.01 | 224.34 |
| 12915 | 3 | 364.26 | 1.72 | 1.75 | 212.39 | 207.98 |
| 21544 | 3 | 606.06 | 3.37 | 3.42 | 179.84 | 177.09 |
| 35938 | 3 | 1044.65 | 6.78 | 7.02 | 153.98 | 148.85 |
| 59948 | 3 | 1817.57 | 15.68 | 15.98 | 115.93 | 113.75 |
| 100000 | 3 | 3039.34 | 35.37 | 35.97 | 85.92 | 84.49 |
| 1000 | 8 | 78.06 | 0.17 | 0.14 | 453.93 | 555.78 |
| 1668 | 8 | 125.58 | 0.25 | 0.19 | 502.42 | 647.59 |
| 2783 | 8 | 212.11 | 0.34 | 0.28 | 625.71 | 744.43 |
| 4642 | 8 | 345.62 | 0.75 | 0.64 | 458.14 | 543.77 |
| 7743 | 8 | 580.13 | 1.13 | 0.99 | 512.45 | 586.34 |
| 12915 | 8 | 971.36 | 1.98 | 1.77 | 490.12 | 550.27 |
| 21544 | 8 | 1616.17 | 3.70 | 3.36 | 436.47 | 481.32 |
| 35938 | 8 | 2785.72 | 7.30 | 7.02 | 381.79 | 396.60 |
| 59948 | 8 | 4846.86 | 15.52 | 15.13 | 312.38 | 320.43 |
| 100000 | 8 | 8104.90 | 36.96 | 34.73 | 219.28 | 233.35 |
| 1000 | 22 | 214.65 | 0.25 | 0.15 | 842.75 | 1447.73 |
| 1668 | 22 | 345.36 | 0.37 | 0.21 | 931.43 | 1651.86 |
| 2783 | 22 | 583.31 | 0.57 | 0.30 | 1028.30 | 1966.79 |
| 4642 | 22 | 950.47 | 1.01 | 0.66 | 945.18 | 1450.26 |
| 7743 | 22 | 1595.35 | 1.57 | 1.06 | 1016.45 | 1500.45 |
| 12915 | 22 | 2671.24 | 2.62 | 1.84 | 1018.63 | 1451.61 |
| 21544 | 22 | 4444.46 | 4.65 | 3.52 | 955.39 | 1262.86 |
| 35938 | 22 | 7660.73 | 8.89 | 7.21 | 861.74 | 1062.13 |
| 59948 | 22 | 13328.88 | 19.06 | 15.43 | 699.20 | 863.57 |
| 100000 | 22 | 22288.46 | 39.72 | 36.47 | 561.13 | 611.09 |
| 1000 | 60 | 585.42 | 0.55 | 0.19 | 1063.57 | 3140.32 |
| 1668 | 60 | 941.88 | 0.77 | 0.23 | 1230.37 | 4052.02 |
| 2783 | 60 | 1590.84 | 1.08 | 0.35 | 1471.76 | 4599.61 |
| 4642 | 60 | 2592.18 | 1.75 | 0.70 | 1481.59 | 3689.30 |

| | | | | | | |
|---|---|---|---|---|---|---|
| 7743 | 60 | 4350.96 | 2.74 | 1.15 | 1586.30 | 3781.51 |
| 12915 | 60 | 7285.20 | 4.34 | 2.02 | 1678.22 | 3613.11 |
| 21544 | 60 | 12121.26 | 7.38 | 3.79 | 1643.47 | 3196.46 |
| 35938 | 60 | 20892.90 | 13.00 | 7.72 | 1607.61 | 2707.35 |
| 59948 | 60 | 36351.48 | 25.51 | 16.06 | 1424.72 | 2262.95 |
| 100000 | 60 | 60786.72 | 52.54 | 37.94 | 1156.91 | 1602.07 |
| 1000 | 167 | 1629.42 | 1.21 | 0.22 | 1350.77 | 7270.34 |
| 1668 | 167 | 2621.57 | 1.96 | 0.32 | 1340.62 | 8206.47 |
| 2783 | 167 | 4427.84 | 2.54 | 0.46 | 1743.91 | 9692.38 |
| 4642 | 167 | 7214.90 | 3.89 | 0.88 | 1854.74 | 8166.37 |
| 7743 | 167 | 12110.17 | 5.94 | 1.42 | 2039.88 | 8538.95 |
| 12915 | 167 | 20277.14 | 9.11 | 2.40 | 2226.30 | 8452.50 |
| 21544 | 167 | 33737.51 | 14.78 | 4.39 | 2282.31 | 7682.55 |
| 35938 | 167 | 58151.91 | 24.77 | 8.62 | 2347.64 | 6745.65 |
| 59948 | 167 | 101178.29 | 44.14 | 18.06 | 2292.43 | 5603.54 |
| 100000 | 167 | 169189.70 | 83.87 | 40.63 | 2017.26 | 4163.68 |
| 1000 | 464 | 4527.25 | 2.97 | 0.36 | 1522.39 | 12595.18 |
| 1668 | 464 | 7283.87 | 4.83 | 0.51 | 1508.38 | 14346.75 |
| 2783 | 464 | 12302.50 | 6.32 | 0.76 | 1947.06 | 16192.13 |
| 4642 | 464 | 20046.19 | 9.07 | 1.27 | 2210.69 | 15724.64 |
| 7743 | 464 | 33647.42 | 14.70 | 1.93 | 2288.93 | 17463.99 |
| 12915 | 464 | 56338.88 | 22.12 | 3.46 | 2547.40 | 16288.74 |
| 21544 | 464 | 93737.74 | 35.16 | 6.04 | 2665.75 | 15511.15 |
| 35938 | 464 | 161571.76 | 56.99 | 11.40 | 2835.28 | 14172.13 |
| 59948 | 464 | 281118.11 | 96.46 | 22.87 | 2914.44 | 12291.16 |
| 100000 | 464 | 470083.97 | 171.67 | 47.54 | 2738.30 | 9887.54 |
| 1000 | 1292 | 12606.04 | 7.89 | 0.67 | 1598.07 | 18806.45 |
| 1668 | 1292 | 20281.82 | 13.46 | 0.85 | 1507.20 | 23859.68 |
| 2783 | 1292 | 34256.09 | 16.06 | 1.27 | 2133.64 | 27078.40 |
| 4642 | 1292 | 55818.28 | 23.85 | 2.26 | 2340.71 | 24670.18 |
| 7743 | 1292 | 93690.67 | 38.81 | 3.43 | 2414.26 | 27280.06 |
| 12915 | 1292 | 156874.64 | 58.35 | 5.67 | 2688.67 | 27666.18 |
| 21544 | 1292 | 261011.13 | 90.71 | 9.72 | 2877.32 | 26855.63 |
| 35938 | 1292 | 449893.78 | 147.07 | 18.43 | 3059.00 | 24406.48 |
| 59948 | 1292 | 782768.54 | 240.67 | 33.27 | 3252.46 | 23525.73 |
| 100000 | 1292 | 1308940.70 | 410.20 | 66.55 | 3191.00 | 19669.97 |
| 1000 | 3594 | 35066.66 | 20.35 | 1.27 | 1723.54 | 27652.51 |
| 1668 | 3594 | 56418.61 | 36.53 | 2.01 | 1544.57 | 28102.33 |
| 2783 | 3594 | 95291.32 | 40.66 | 2.91 | 2343.68 | 32743.53 |

| | | | | | | |
|---|---|---|---|---|---|---|
| 4642 | 3594 | 155271.58 | 66.23 | 4.72 | 2344.33 | 32911.60 |
| 7743 | 3594 | 260622.50 | 108.15 | 6.96 | 2409.88 | 37424.77 |
| 12915 | 3594 | 436383.48 | 159.23 | 12.07 | 2740.66 | 36149.66 |
| 21544 | 3594 | 726063.47 | 244.26 | 20.30 | 2972.54 | 35765.93 |
| 35938 | 3594 | 1251484.71 | 389.79 | 35.25 | 3210.67 | 35499.33 |
| 59948 | 3594 | 2177453.65 | 624.35 | 60.16 | 3487.55 | 36191.97 |
| 100000 | 3594 | 3641124.53 | 1067.67 | 117.83 | 3410.36 | 30901.16 |
| 1000 | 10000 | 97570.00 | 54.27 | 2.98 | 1797.85 | 32710.94 |
| 1668 | 10000 | 156980.00 | 99.82 | 4.49 | 1572.57 | 34926.95 |
| 2783 | 10000 | 265140.00 | 111.76 | 7.29 | 2372.46 | 36366.34 |
| 4642 | 10000 | 432030.00 | 179.45 | 11.54 | 2407.50 | 37432.32 |
| 7743 | 10000 | 725160.00 | 285.13 | 18.49 | 2543.26 | 39223.00 |
| 12915 | 10000 | 1214200.00 | 426.80 | 30.16 | 2844.89 | 40259.04 |
| 21544 | 10000 | 2020210.00 | 658.69 | 50.26 | 3067.02 | 40198.00 |
| 35938 | 10000 | 3482150.00 | 1060.28 | 84.43 | 3284.19 | 41242.15 |
| 59948 | 10000 | 6058580.00 | 1722.00 | 143.92 | 3518.34 | 42096.52 |
| 100000 | 10000 | 10131120.00 | 2850.32 | 254.42 | 3554.37 | 39820.70 |

#### Single outcome variable with non-default settings

**Table S3: Comparison of time taken (in seconds) by glmmTMB and FEMA for a single outcome variable using non-default settings.** For this analysis, we set iter.max=5000 and eval.max=5000 for glmmTMB

| Number of observations | Median time glmmTMB | Median time FEMA (serial) | FEMA speed-up factor (serial) |
|---|---|---|---|
| 1000 | 26.23 | 0.14 | 190.34 |
| 1668 | 40.55 | 0.16 | 261.19 |
| 2783 | 68.29 | 0.25 | 272.15 |
| 4642 | 78.48 | 0.51 | 153.48 |
| 7743 | 129.06 | 0.86 | 149.26 |
| 12915 | 222.16 | 1.60 | 138.95 |
| 21544 | 367.77 | 3.10 | 118.53 |
| 35938 | 675.97 | 6.42 | 105.35 |
| 59948 | 1188.62 | 14.21 | 83.64 |
| 100000 | 1894.25 | 34.80 | 54.44 |

#### Multiple outcome variables with non-default settings

**Table S4: Comparison of time taken (in seconds) by glmmTMB and FEMA for multiple outcome variables using non-default settings.** The glmmTMB timings are extrapolated from the time taken for a single outcome variable with iter.max=5000 and eval.max=5000 settings for glmmTMB

| No. of observations | No. of *y* vars. | Median time glmmTMB | Median time FEMA (serial) | Median time FEMA (parallel) | FEMA speed-up factor (serial) | FEMA speed-up factor (parallel) |
|---|---|---|---|---|---|---|
| 1000 | 3 | 78.69 | 0.13 | 0.14 | 628.16 | 546.88 |
| 1668 | 3 | 121.64 | 0.19 | 0.18 | 654.92 | 685.00 |
| 2783 | 3 | 204.86 | 0.28 | 0.28 | 724.95 | 722.53 |
| 4642 | 3 | 235.44 | 0.59 | 0.60 | 399.18 | 391.79 |
| 7743 | 3 | 387.17 | 0.98 | 0.97 | 395.11 | 399.26 |
| 12915 | 3 | 666.48 | 1.72 | 1.75 | 388.61 | 380.53 |
| 21544 | 3 | 1103.31 | 3.37 | 3.42 | 327.39 | 322.39 |
| 35938 | 3 | 2027.91 | 6.78 | 7.02 | 298.92 | 288.95 |
| 59948 | 3 | 3565.85 | 15.68 | 15.98 | 227.44 | 223.16 |
| 100000 | 3 | 5682.75 | 35.37 | 35.97 | 160.65 | 157.98 |
| 1000 | 8 | 209.85 | 0.17 | 0.14 | 1220.36 | 1494.18 |
| 1668 | 8 | 324.36 | 0.25 | 0.19 | 1297.66 | 1672.62 |
| 2783 | 8 | 546.28 | 0.34 | 0.28 | 1611.47 | 1917.24 |
| 4642 | 8 | 627.83 | 0.75 | 0.64 | 832.21 | 987.77 |
| 7743 | 8 | 1032.46 | 1.13 | 0.99 | 912.02 | 1043.51 |
| 12915 | 8 | 1777.29 | 1.98 | 1.77 | 896.77 | 1006.83 |
| 21544 | 8 | 2942.17 | 3.70 | 3.36 | 794.57 | 876.22 |
| 35938 | 8 | 5407.76 | 7.30 | 7.02 | 741.14 | 769.89 |
| 59948 | 8 | 9508.94 | 15.52 | 15.13 | 612.85 | 628.64 |
| 100000 | 8 | 15154.01 | 36.96 | 34.73 | 410.00 | 436.30 |
| 1000 | 22 | 577.08 | 0.25 | 0.15 | 2265.68 | 3892.12 |
| 1668 | 22 | 891.99 | 0.37 | 0.21 | 2405.72 | 4266.44 |
| 2783 | 22 | 1502.27 | 0.57 | 0.30 | 2648.31 | 5065.34 |
| 4642 | 22 | 1726.54 | 1.01 | 0.66 | 1716.93 | 2634.42 |
| 7743 | 22 | 2839.25 | 1.57 | 1.06 | 1808.98 | 2670.35 |
| 12915 | 22 | 4887.54 | 2.62 | 1.84 | 1863.78 | 2655.99 |
| 21544 | 22 | 8090.96 | 4.65 | 3.52 | 1739.25 | 2298.99 |
| 35938 | 22 | 14871.34 | 8.89 | 7.21 | 1672.84 | 2061.85 |
| 59948 | 22 | 26149.60 | 19.06 | 15.43 | 1371.74 | 1694.21 |
| 100000 | 22 | 41673.52 | 39.72 | 36.47 | 1049.17 | 1142.57 |
| 1000 | 60 | 1573.86 | 0.55 | 0.19 | 2859.33 | 8442.54 |
| 1668 | 60 | 2432.70 | 0.77 | 0.23 | 3177.82 | 10465.61 |
| 2783 | 60 | 4097.10 | 1.08 | 0.35 | 3790.41 | 11845.98 |

| | | | | | | |
|---|---|---|---|---|---|---|
| 4642 | 60 | 4708.74 | 1.75 | 0.70 | 2691.33 | 6701.67 |
| 7743 | 60 | 7743.42 | 2.74 | 1.15 | 2823.14 | 6729.97 |
| 12915 | 60 | 13329.66 | 4.34 | 2.02 | 3070.63 | 6610.87 |
| 21544 | 60 | 22066.26 | 7.38 | 3.79 | 2991.88 | 5819.03 |
| 35938 | 60 | 40558.20 | 13.00 | 7.72 | 3120.76 | 5255.62 |
| 59948 | 60 | 71317.08 | 25.51 | 16.06 | 2795.13 | 4439.64 |
| 100000 | 60 | 113655.06 | 52.54 | 37.94 | 2163.11 | 2995.45 |
| 1000 | 167 | 4380.58 | 1.21 | 0.22 | 3631.45 | 19545.79 |
| 1668 | 167 | 6771.02 | 1.96 | 0.32 | 3462.56 | 21195.78 |
| 2783 | 167 | 11403.60 | 2.54 | 0.46 | 4491.31 | 24962.05 |
| 4642 | 167 | 13105.99 | 3.89 | 0.88 | 3369.18 | 14834.35 |
| 7743 | 167 | 21552.52 | 5.94 | 1.42 | 3630.39 | 15196.79 |
| 12915 | 167 | 37100.89 | 9.11 | 2.40 | 4073.43 | 15465.46 |
| 21544 | 167 | 61417.76 | 14.78 | 4.39 | 4154.85 | 13985.77 |
| 35938 | 167 | 112886.99 | 24.77 | 8.62 | 4557.33 | 13094.95 |
| 59948 | 167 | 198499.21 | 44.14 | 18.06 | 4497.47 | 10993.45 |
| 100000 | 167 | 316339.92 | 83.87 | 40.63 | 3771.74 | 7784.98 |
| 1000 | 464 | 12171.18 | 2.97 | 0.36 | 4092.83 | 33861.25 |
| 1668 | 464 | 18812.88 | 4.83 | 0.51 | 3895.85 | 37054.97 |
| 2783 | 464 | 31684.24 | 6.32 | 0.76 | 5014.51 | 41701.72 |
| 4642 | 464 | 36414.26 | 9.07 | 1.27 | 4015.76 | 28564.07 |
| 7743 | 464 | 59882.45 | 14.70 | 1.93 | 4073.61 | 31080.73 |
| 12915 | 464 | 103082.70 | 22.12 | 3.46 | 4660.96 | 29803.35 |
| 21544 | 464 | 170645.74 | 35.16 | 6.04 | 4852.89 | 28237.42 |
| 35938 | 464 | 313650.08 | 56.99 | 11.40 | 5503.97 | 27511.54 |
| 59948 | 464 | 551518.75 | 96.46 | 22.87 | 5717.78 | 24113.72 |
| 100000 | 464 | 878932.46 | 171.67 | 47.54 | 5119.90 | 18487.07 |
| 1000 | 1292 | 33890.45 | 7.89 | 0.67 | 4296.30 | 50559.81 |
| 1668 | 1292 | 52384.14 | 13.46 | 0.85 | 3892.82 | 61625.09 |
| 2783 | 1292 | 88224.22 | 16.06 | 1.27 | 5495.04 | 69738.57 |
| 4642 | 1292 | 101394.87 | 23.85 | 2.26 | 4251.95 | 44813.81 |
| 7743 | 1292 | 166741.64 | 38.81 | 3.43 | 4296.67 | 48550.43 |
| 12915 | 1292 | 287032.01 | 58.35 | 5.67 | 4919.44 | 50620.55 |
| 21544 | 1292 | 475160.13 | 90.71 | 9.72 | 5238.05 | 48889.58 |
| 35938 | 1292 | 873353.24 | 147.07 | 18.43 | 5938.26 | 47378.91 |
| 59948 | 1292 | 1535694.46 | 240.67 | 33.27 | 6380.93 | 46154.55 |
| 100000 | 1292 | 2447372.29 | 410.20 | 66.55 | 5966.33 | 36777.63 |
| 1000 | 3594 | 94274.21 | 20.35 | 1.27 | 4633.61 | 74341.82 |
| 1668 | 3594 | 145718.73 | 36.53 | 2.01 | 3989.33 | 72583.07 |

| | | | | | | |
|---|---|---|---|---|---|---|
| 2783 | 3594 | 245416.29 | 40.66 | 2.91 | 6036.00 | 84328.72 |
| 4642 | 3594 | 282053.53 | 66.23 | 4.72 | 4258.52 | 59784.49 |
| 7743 | 3594 | 463830.86 | 108.15 | 6.96 | 4288.88 | 66605.01 |
| 12915 | 3594 | 798446.63 | 159.23 | 12.07 | 5014.57 | 66142.68 |
| 21544 | 3594 | 1321768.97 | 244.26 | 20.30 | 5411.39 | 65110.42 |
| 35938 | 3594 | 2429436.18 | 389.79 | 35.25 | 6232.68 | 68912.84 |
| 59948 | 3594 | 4271893.09 | 624.35 | 60.16 | 6842.13 | 71004.13 |
| 100000 | 3594 | 6807938.09 | 1067.67 | 117.83 | 6376.46 | 57776.98 |
| 1000 | 10000 | 262310.00 | 54.27 | 2.98 | 4833.40 | 87941.03 |
| 1668 | 10000 | 405450.00 | 99.82 | 4.49 | 4061.66 | 90209.78 |
| 2783 | 10000 | 682850.00 | 111.76 | 7.29 | 6110.11 | 93659.04 |
| 4642 | 10000 | 784790.00 | 179.45 | 11.54 | 4373.27 | 67996.47 |
| 7743 | 10000 | 1290570.00 | 285.13 | 18.49 | 4526.25 | 69805.32 |
| 12915 | 10000 | 2221610.00 | 426.80 | 30.16 | 5205.26 | 73661.58 |
| 21544 | 10000 | 3677710.00 | 658.69 | 50.26 | 5583.38 | 73178.83 |
| 35938 | 10000 | 6759700.00 | 1060.28 | 84.43 | 6375.42 | 80061.05 |
| 59948 | 10000 | 11886180.00 | 1722.00 | 143.92 | 6902.55 | 82588.13 |
| 100000 | 10000 | 18942510.00 | 2850.32 | 254.42 | 6645.74 | 74454.15 |

### Simulation 3: comparison of computational timing with lmer

#### Single outcome variable

**Table S5: Comparison of time taken (in seconds) by lmer and FEMA for a single outcome variable**

| Number of observations | Median time lmer | Median time FEMA (serial) | FEMA speed-up factor (serial) |
|---|---|---|---|
| 1000 | 85.66 | 0.14 | 622 |
| 1668 | 46.64 | 0.16 | 300 |
| 2783 | 59.26 | 0.25 | 236 |
| 4642 | 120.54 | 0.51 | 236 |
| 7743 | 171.11 | 0.86 | 198 |
| 12915 | 331.33 | 1.60 | 207 |
| 21544 | 569.95 | 3.10 | 184 |
| 35938 | 1212.30 | 6.42 | 189 |
| 59948 | 1778.05 | 14.21 | 125 |
| 100000 | 3104.90 | 34.80 | 89 |

#### Multiple outcome variable

**Table S6**: **Comparison of time taken (in seconds) by lmer and FEMA for multiple outcome variables.** The lmer timings are extrapolated from the time taken for a single outcome variable

| No. of observations | No. of *y* vars. | Median time lmer | Median time FEMA (serial) | Median time FEMA (parallel) | FEMA speed-up factor (serial) | FEMA speed-up factor (parallel) |
|---|---|---|---|---|---|---|
| 1000 | 3 | 256.99 | 0.13 | 0.14 | 2051.38 | 1785.96 |
| 1668 | 3 | 139.93 | 0.19 | 0.18 | 753.43 | 788.04 |
| 2783 | 3 | 177.77 | 0.28 | 0.28 | 629.08 | 626.99 |
| 4642 | 3 | 361.61 | 0.59 | 0.60 | 613.11 | 601.76 |
| 7743 | 3 | 513.34 | 0.98 | 0.97 | 523.86 | 529.37 |
| 12915 | 3 | 993.98 | 1.72 | 1.75 | 579.56 | 567.52 |
| 21544 | 3 | 1709.85 | 3.37 | 3.42 | 507.37 | 499.62 |
| 35938 | 3 | 3636.90 | 6.78 | 7.02 | 536.09 | 518.20 |
| 59948 | 3 | 5334.14 | 15.68 | 15.98 | 340.22 | 333.82 |
| 100000 | 3 | 9314.71 | 35.37 | 35.97 | 263.33 | 258.95 |
| 1000 | 8 | 685.30 | 0.17 | 0.14 | 3985.34 | 4879.56 |
| 1668 | 8 | 373.15 | 0.25 | 0.19 | 1492.86 | 1924.22 |
| 2783 | 8 | 474.04 | 0.34 | 0.28 | 1398.37 | 1663.70 |
| 4642 | 8 | 964.30 | 0.75 | 0.64 | 1278.21 | 1517.14 |
| 7743 | 8 | 1368.91 | 1.13 | 0.99 | 1209.22 | 1383.57 |
| 12915 | 8 | 2650.61 | 1.98 | 1.77 | 1337.42 | 1501.57 |
| 21544 | 8 | 4559.60 | 3.70 | 3.36 | 1231.38 | 1357.92 |
| 35938 | 8 | 9698.39 | 7.30 | 7.02 | 1329.18 | 1380.74 |
| 59948 | 8 | 14224.36 | 15.52 | 15.13 | 916.76 | 940.37 |
| 100000 | 8 | 24839.22 | 36.96 | 34.73 | 672.04 | 715.14 |
| 1000 | 22 | 1884.59 | 0.25 | 0.15 | 7399.07 | 12710.57 |
| 1668 | 22 | 1026.17 | 0.37 | 0.21 | 2767.60 | 4908.22 |
| 2783 | 22 | 1303.61 | 0.57 | 0.30 | 2298.09 | 4395.50 |
| 4642 | 22 | 2651.81 | 1.01 | 0.66 | 2637.05 | 4046.24 |
| 7743 | 22 | 3764.51 | 1.57 | 1.06 | 2398.49 | 3540.56 |
| 12915 | 22 | 7289.17 | 2.62 | 1.84 | 2779.61 | 3961.09 |
| 21544 | 22 | 12538.90 | 4.65 | 3.52 | 2695.39 | 3562.84 |
| 35938 | 22 | 26670.58 | 8.89 | 7.21 | 3000.11 | 3697.76 |
| 59948 | 22 | 39116.99 | 19.06 | 15.43 | 2051.97 | 2534.35 |
| 100000 | 22 | 68307.84 | 39.72 | 36.47 | 1719.72 | 1872.81 |
| 1000 | 60 | 5139.78 | 0.55 | 0.19 | 9337.76 | 27570.93 |
| 1668 | 60 | 2798.64 | 0.77 | 0.23 | 3655.84 | 12039.91 |
| 2783 | 60 | 3555.30 | 1.08 | 0.35 | 3289.16 | 10279.47 |
| 4642 | 60 | 7232.22 | 1.75 | 0.70 | 4133.65 | 10293.20 |

| | | | | | | |
|---|---|---|---|---|---|---|
| 7743 | 60 | 10266.84 | 2.74 | 1.15 | 3743.14 | 8923.13 |
| 12915 | 60 | 19879.56 | 4.34 | 2.02 | 4579.46 | 9859.31 |
| 21544 | 60 | 34197.00 | 7.38 | 3.79 | 4636.64 | 9018.00 |
| 35938 | 60 | 72737.94 | 13.00 | 7.72 | 5596.83 | 9425.53 |
| 59948 | 60 | 106682.70 | 25.51 | 16.06 | 4181.21 | 6641.22 |
| 100000 | 60 | 186294.12 | 52.54 | 37.94 | 3545.60 | 4909.89 |
| 1000 | 167 | 14305.72 | 1.21 | 0.22 | 11859.27 | 63831.01 |
| 1668 | 167 | 7789.55 | 1.96 | 0.32 | 3983.42 | 24384.16 |
| 2783 | 167 | 9895.59 | 2.54 | 0.46 | 3897.38 | 21661.08 |
| 4642 | 167 | 20129.68 | 3.89 | 0.88 | 5174.76 | 22784.29 |
| 7743 | 167 | 28576.04 | 5.94 | 1.42 | 4813.45 | 20149.11 |
| 12915 | 167 | 55331.44 | 9.11 | 2.40 | 6075.03 | 23064.85 |
| 21544 | 167 | 95181.65 | 14.78 | 4.39 | 6438.94 | 21674.33 |
| 35938 | 167 | 202453.93 | 24.77 | 8.62 | 8173.22 | 23484.76 |
| 59948 | 167 | 296933.52 | 44.14 | 18.06 | 6727.73 | 16445.03 |
| 100000 | 167 | 518518.63 | 83.87 | 40.63 | 6182.32 | 12760.50 |
| 1000 | 464 | 39747.63 | 2.97 | 0.36 | 13366.02 | 110581.23 |
| 1668 | 464 | 21642.82 | 4.83 | 0.51 | 4481.89 | 42628.98 |
| 2783 | 464 | 27494.32 | 6.32 | 0.76 | 4351.39 | 36187.09 |
| 4642 | 464 | 55929.17 | 9.07 | 1.27 | 6167.86 | 43871.96 |
| 7743 | 464 | 79396.90 | 14.70 | 1.93 | 5401.12 | 41209.30 |
| 12915 | 464 | 153735.26 | 22.12 | 3.46 | 6951.25 | 44448.06 |
| 21544 | 464 | 264456.80 | 35.16 | 6.04 | 7520.73 | 43760.70 |
| 35938 | 464 | 562506.74 | 56.99 | 11.40 | 9870.94 | 49339.79 |
| 59948 | 464 | 825012.88 | 96.46 | 22.87 | 8553.18 | 36071.54 |
| 100000 | 464 | 1440674.53 | 171.67 | 47.54 | 8392.12 | 30302.51 |
| 1000 | 1292 | 110676.60 | 7.89 | 0.67 | 14030.49 | 165114.00 |
| 1668 | 1292 | 60264.05 | 13.46 | 0.85 | 4478.40 | 70895.07 |
| 2783 | 1292 | 76557.46 | 16.06 | 1.27 | 4768.37 | 60516.35 |
| 4642 | 1292 | 155733.80 | 23.85 | 2.26 | 6530.62 | 68830.16 |
| 7743 | 1292 | 221079.29 | 38.81 | 3.43 | 5696.87 | 64372.01 |
| 12915 | 1292 | 428073.19 | 58.35 | 5.67 | 7336.74 | 75494.36 |
| 21544 | 1292 | 736375.40 | 90.71 | 9.72 | 8117.62 | 75766.21 |
| 35938 | 1292 | 1566290.31 | 147.07 | 18.43 | 10649.81 | 84970.35 |
| 59948 | 1292 | 2297234.14 | 240.67 | 33.27 | 9545.19 | 69042.25 |
| 100000 | 1292 | 4011533.38 | 410.20 | 66.55 | 9779.52 | 60282.90 |
| 1000 | 3594 | 307872.82 | 20.35 | 1.27 | 15132.06 | 242779.27 |
| 1668 | 3594 | 167638.54 | 36.53 | 2.01 | 4589.43 | 83501.41 |
| 2783 | 3594 | 212962.47 | 40.66 | 2.91 | 5237.80 | 73177.10 |

| | | | | | | |
|---|---|---|---|---|---|---|
| 4642 | 3594 | 433209.98 | 66.23 | 4.72 | 6540.72 | 91823.83 |
| 7743 | 3594 | 614983.72 | 108.15 | 6.96 | 5686.53 | 88310.21 |
| 12915 | 3594 | 1190785.64 | 159.23 | 12.07 | 7478.62 | 98643.73 |
| 21544 | 3594 | 2048400.30 | 244.26 | 20.30 | 8386.26 | 100904.32 |
| 35938 | 3594 | 4357002.61 | 389.79 | 35.25 | 11177.82 | 123589.76 |
| 59948 | 3594 | 6390293.73 | 624.35 | 60.16 | 10235.10 | 106214.56 |
| 100000 | 3594 | 11159017.79 | 1067.67 | 117.83 | 10451.78 | 94703.33 |
| 1000 | 10000 | 856630.00 | 54.27 | 2.98 | 15784.50 | 287190.45 |
| 1668 | 10000 | 466440.00 | 99.82 | 4.49 | 4672.64 | 103779.62 |
| 2783 | 10000 | 592550.00 | 111.76 | 7.29 | 5302.11 | 81273.58 |
| 4642 | 10000 | 1205370.00 | 179.45 | 11.54 | 6716.96 | 104436.73 |
| 7743 | 10000 | 1711140.00 | 285.13 | 18.49 | 6001.27 | 92553.43 |
| 12915 | 10000 | 3313260.00 | 426.80 | 30.16 | 7763.02 | 109857.25 |
| 21544 | 10000 | 5699500.00 | 658.69 | 50.26 | 8652.80 | 113408.27 |
| 35938 | 10000 | 12122990.00 | 1060.28 | 84.43 | 11433.81 | 143583.19 |
| 59948 | 10000 | 17780450.00 | 1722.00 | 143.92 | 10325.47 | 123542.98 |
| 100000 | 10000 | 31049020.00 | 2850.32 | 254.42 | 10893.15 | 122039.18 |

#### Simulation 3: comparison of carbon footprint with glmmTMB

##### Using default settings for glmmTMB

**Table S7: Comparison of carbon footprint (in grams of carbon dioxide emission) of glmmTMB and FEMA.** The glmmTMB carbon footprint are extrapolated from the carbon footprint for a single outcome variable; the green ratio is the ratio of carbon footprint of glmmTMB to the carbon footprint of FEMA

| No. of observations | No. of *y* vars. | Carbon footprint glmmTMB ($gCO_2e$) | Carbon footprint FEMA (serial; $gCO_2e$) | Carbon footprint FEMA (parallel; $gCO_2e$) | FEMA green ratio (serial) | FEMA green ratio (parallel) |
|---|---|---|---|---|---|---|
| 1000 | 1 | 1.78 | 0.03 | 0.03 | 70.80 | 52.13 |
| 1668 | 1 | 2.87 | 0.03 | 0.03 | 101.13 | 97.87 |
| 2783 | 1 | 4.84 | 0.05 | 0.05 | 105.67 | 105.33 |
| 4642 | 1 | 7.89 | 0.09 | 0.10 | 84.49 | 76.87 |
| 7743 | 1 | 13.24 | 0.16 | 0.18 | 83.87 | 74.73 |
| 12915 | 1 | 22.17 | 0.29 | 0.32 | 75.94 | 70.08 |
| 21544 | 1 | 36.89 | 0.57 | 0.60 | 65.11 | 61.34 |
| 35938 | 1 | 63.59 | 1.17 | 1.25 | 54.27 | 50.88 |
| 59948 | 1 | 110.64 | 2.60 | 2.74 | 42.63 | 40.38 |
| 100000 | 1 | 185.01 | 6.35 | 6.58 | 29.11 | 28.11 |
| 1000 | 3 | 5.35 | 0.02 | 0.03 | 233.65 | 203.42 |
| 1668 | 3 | 8.60 | 0.03 | 0.03 | 253.57 | 265.22 |
| 2783 | 3 | 14.53 | 0.05 | 0.05 | 281.49 | 280.55 |
| 4642 | 3 | 23.67 | 0.11 | 0.11 | 219.75 | 215.68 |
| 7743 | 3 | 39.73 | 0.18 | 0.18 | 222.01 | 224.34 |
| 12915 | 3 | 66.52 | 0.31 | 0.32 | 212.39 | 207.98 |
| 21544 | 3 | 110.68 | 0.62 | 0.62 | 179.84 | 177.09 |
| 35938 | 3 | 190.77 | 1.24 | 1.28 | 153.98 | 148.85 |
| 59948 | 3 | 331.92 | 2.86 | 2.92 | 115.93 | 113.75 |
| 100000 | 3 | 555.03 | 6.46 | 6.57 | 85.92 | 84.49 |
| 1000 | 8 | 14.25 | 0.03 | 0.03 | 453.93 | 555.78 |
| 1668 | 8 | 22.93 | 0.05 | 0.04 | 502.42 | 647.59 |
| 2783 | 8 | 38.73 | 0.06 | 0.05 | 625.71 | 744.43 |
| 4642 | 8 | 63.12 | 0.14 | 0.12 | 458.14 | 543.77 |
| 7743 | 8 | 105.94 | 0.21 | 0.18 | 512.45 | 586.34 |
| 12915 | 8 | 177.39 | 0.36 | 0.32 | 490.12 | 550.27 |
| 21544 | 8 | 295.14 | 0.68 | 0.61 | 436.47 | 481.32 |
| 35938 | 8 | 508.72 | 1.33 | 1.28 | 381.79 | 396.60 |
| 59948 | 8 | 885.11 | 2.83 | 2.76 | 312.38 | 320.43 |
| 100000 | 8 | 1480.08 | 6.75 | 6.34 | 219.28 | 233.35 |
| 1000 | 22 | 39.20 | 0.05 | 0.03 | 842.75 | 1447.73 |

| | | | | | | |
|---|---|---|---|---|---|---|
| 1668 | 22 | 63.07 | 0.07 | 0.04 | 931.43 | 1651.86 |
| 2783 | 22 | 106.52 | 0.10 | 0.05 | 1028.30 | 1966.79 |
| 4642 | 22 | 173.57 | 0.18 | 0.12 | 945.18 | 1450.26 |
| 7743 | 22 | 291.34 | 0.29 | 0.19 | 1016.45 | 1500.45 |
| 12915 | 22 | 487.81 | 0.48 | 0.34 | 1018.63 | 1451.61 |
| 21544 | 22 | 811.63 | 0.85 | 0.64 | 955.39 | 1262.86 |
| 35938 | 22 | 1398.97 | 1.62 | 1.32 | 861.74 | 1062.13 |
| 59948 | 22 | 2434.06 | 3.48 | 2.82 | 699.20 | 863.57 |
| 100000 | 22 | 4070.22 | 7.25 | 6.66 | 561.13 | 611.09 |
| 1000 | 60 | 106.91 | 0.10 | 0.03 | 1063.57 | 3140.32 |
| 1668 | 60 | 172.00 | 0.14 | 0.04 | 1230.37 | 4052.02 |
| 2783 | 60 | 290.51 | 0.20 | 0.06 | 1471.76 | 4599.61 |
| 4642 | 60 | 473.37 | 0.32 | 0.13 | 1481.59 | 3689.30 |
| 7743 | 60 | 794.55 | 0.50 | 0.21 | 1586.30 | 3781.51 |
| 12915 | 60 | 1330.39 | 0.79 | 0.37 | 1678.22 | 3613.11 |
| 21544 | 60 | 2213.53 | 1.35 | 0.69 | 1643.47 | 3196.46 |
| 35938 | 60 | 3815.36 | 2.37 | 1.41 | 1607.61 | 2707.35 |
| 59948 | 60 | 6638.34 | 4.66 | 2.93 | 1424.72 | 2262.95 |
| 100000 | 60 | 11100.59 | 9.60 | 6.93 | 1156.91 | 1602.07 |
| 1000 | 167 | 297.56 | 0.22 | 0.04 | 1350.77 | 7270.34 |
| 1668 | 167 | 478.74 | 0.36 | 0.06 | 1340.62 | 8206.47 |
| 2783 | 167 | 808.59 | 0.46 | 0.08 | 1743.91 | 9692.38 |
| 4642 | 167 | 1317.55 | 0.71 | 0.16 | 1854.74 | 8166.37 |
| 7743 | 167 | 2211.50 | 1.08 | 0.26 | 2039.88 | 8538.95 |
| 12915 | 167 | 3702.92 | 1.66 | 0.44 | 2226.30 | 8452.50 |
| 21544 | 167 | 6160.99 | 2.70 | 0.80 | 2282.31 | 7682.55 |
| 35938 | 167 | 10619.43 | 4.52 | 1.57 | 2347.64 | 6745.65 |
| 59948 | 167 | 18476.71 | 8.06 | 3.30 | 2292.43 | 5603.54 |
| 100000 | 167 | 30896.63 | 15.32 | 7.42 | 2017.26 | 4163.68 |
| 1000 | 464 | 826.74 | 0.54 | 0.07 | 1522.39 | 12595.18 |
| 1668 | 464 | 1330.15 | 0.88 | 0.09 | 1508.38 | 14346.75 |
| 2783 | 464 | 2246.62 | 1.15 | 0.14 | 1947.06 | 16192.13 |
| 4642 | 464 | 3660.74 | 1.66 | 0.23 | 2210.69 | 15724.64 |
| 7743 | 464 | 6144.54 | 2.68 | 0.35 | 2288.93 | 17463.99 |
| 12915 | 464 | 10288.34 | 4.04 | 0.63 | 2547.40 | 16288.74 |
| 21544 | 464 | 17117.95 | 6.42 | 1.10 | 2665.75 | 15511.15 |
| 35938 | 464 | 29505.48 | 10.41 | 2.08 | 2835.28 | 14172.13 |
| 59948 | 464 | 51336.48 | 17.61 | 4.18 | 2914.44 | 12291.16 |
| 100000 | 464 | 85844.54 | 31.35 | 8.68 | 2738.30 | 9887.54 |

| | | | | | | |
|---|---|---|---|---|---|---|
| 1000 | 1292 | 2302.06 | 1.44 | 0.12 | 1598.07 | 18806.45 |
| 1668 | 1292 | 3703.77 | 2.46 | 0.16 | 1507.20 | 23859.68 |
| 2783 | 1292 | 6255.69 | 2.93 | 0.23 | 2133.64 | 27078.40 |
| 4642 | 1292 | 10193.27 | 4.35 | 0.41 | 2340.71 | 24670.18 |
| 7743 | 1292 | 17109.35 | 7.09 | 0.63 | 2414.26 | 27280.06 |
| 12915 | 1292 | 28647.71 | 10.65 | 1.04 | 2688.67 | 27666.18 |
| 21544 | 1292 | 47664.63 | 16.57 | 1.77 | 2877.32 | 26855.63 |
| 35938 | 1292 | 82157.50 | 26.86 | 3.37 | 3059.00 | 24406.48 |
| 59948 | 1292 | 142945.54 | 43.95 | 6.08 | 3252.46 | 23525.73 |
| 100000 | 1292 | 239032.64 | 74.91 | 12.15 | 3191.00 | 19669.97 |
| 1000 | 3594 | 6403.71 | 3.72 | 0.23 | 1723.54 | 27652.51 |
| 1668 | 3594 | 10302.90 | 6.67 | 0.37 | 1544.57 | 28102.33 |
| 2783 | 3594 | 17401.66 | 7.42 | 0.53 | 2343.68 | 32743.53 |
| 4642 | 3594 | 28354.97 | 12.10 | 0.86 | 2344.33 | 32911.60 |
| 7743 | 3594 | 47593.67 | 19.75 | 1.27 | 2409.88 | 37424.77 |
| 12915 | 3594 | 79690.31 | 29.08 | 2.20 | 2740.66 | 36149.66 |
| 21544 | 3594 | 132590.32 | 44.61 | 3.71 | 2972.54 | 35765.93 |
| 35938 | 3594 | 228540.30 | 71.18 | 6.44 | 3210.67 | 35499.33 |
| 59948 | 3594 | 397636.42 | 114.02 | 10.99 | 3487.55 | 36191.97 |
| 100000 | 3594 | 664925.17 | 194.97 | 21.52 | 3410.36 | 30901.16 |
| 1000 | 10000 | 17817.78 | 9.91 | 0.54 | 1797.85 | 32710.94 |
| 1668 | 10000 | 28666.96 | 18.23 | 0.82 | 1572.57 | 34926.95 |
| 2783 | 10000 | 48418.63 | 20.41 | 1.33 | 2372.46 | 36366.34 |
| 4642 | 10000 | 78895.30 | 32.77 | 2.11 | 2407.50 | 37432.32 |
| 7743 | 10000 | 132425.34 | 52.07 | 3.38 | 2543.26 | 39223.00 |
| 12915 | 10000 | 221731.54 | 77.94 | 5.51 | 2844.89 | 40259.04 |
| 21544 | 10000 | 368921.32 | 120.29 | 9.18 | 3067.02 | 40198.00 |
| 35938 | 10000 | 635893.98 | 193.62 | 15.42 | 3284.19 | 41242.15 |
| 59948 | 10000 | 1106389.61 | 314.46 | 26.28 | 3518.34 | 42096.52 |
| 100000 | 10000 | 1850097.86 | 520.51 | 46.46 | 3554.37 | 39820.70 |

#### Using non-default settings for glmmTMB

**Table S8: Comparison of carbon footprint (in grams of carbon dioxide emission) of glmmTMB and FEMA.** The glmmTMB carbon footprint are extrapolated from the carbon footprint for a single outcome variable with iter.max=5000 and eval.max=5000 settings for glmmTMB; the green ratio is the ratio of carbon footprint of glmmTMB to the carbon footprint of FEMA

| No. of observations | No. of *y* vars. | Carbon footprint glmmTMB ($gCO_2e$) | Carbon footprint FEMA (serial; $gCO_2e$) | Carbon footprint FEMA (parallel; $gCO_2e$) | FEMA green ratio (serial) | FEMA green ratio (parallel) |
|---|---|---|---|---|---|---|
| 1000 | 1 | 4.79 | 0.03 | 0.03 | 190.34 | 140.14 |
| 1668 | 1 | 7.40 | 0.03 | 0.03 | 261.19 | 252.79 |
| 2783 | 1 | 12.47 | 0.05 | 0.05 | 272.15 | 271.28 |
| 4642 | 1 | 14.33 | 0.09 | 0.10 | 153.48 | 139.63 |
| 7743 | 1 | 23.57 | 0.16 | 0.18 | 149.26 | 132.99 |
| 12915 | 1 | 40.57 | 0.29 | 0.32 | 138.95 | 128.22 |
| 21544 | 1 | 67.16 | 0.57 | 0.60 | 118.53 | 111.67 |
| 35938 | 1 | 123.44 | 1.17 | 1.25 | 105.35 | 98.78 |
| 59948 | 1 | 217.06 | 2.60 | 2.74 | 83.64 | 79.22 |
| 100000 | 1 | 345.92 | 6.35 | 6.58 | 54.44 | 52.55 |
| 1000 | 3 | 14.37 | 0.02 | 0.03 | 628.16 | 546.88 |
| 1668 | 3 | 22.21 | 0.03 | 0.03 | 654.92 | 685.00 |
| 2783 | 3 | 37.41 | 0.05 | 0.05 | 724.95 | 722.53 |
| 4642 | 3 | 42.99 | 0.11 | 0.11 | 399.18 | 391.79 |
| 7743 | 3 | 70.70 | 0.18 | 0.18 | 395.11 | 399.26 |
| 12915 | 3 | 121.71 | 0.31 | 0.32 | 388.61 | 380.53 |
| 21544 | 3 | 201.48 | 0.62 | 0.62 | 327.39 | 322.39 |
| 35938 | 3 | 370.33 | 1.24 | 1.28 | 298.92 | 288.95 |
| 59948 | 3 | 651.18 | 2.86 | 2.92 | 227.44 | 223.16 |
| 100000 | 3 | 1037.76 | 6.46 | 6.57 | 160.65 | 157.98 |
| 1000 | 8 | 38.32 | 0.03 | 0.03 | 1220.36 | 1494.18 |
| 1668 | 8 | 59.23 | 0.05 | 0.04 | 1297.66 | 1672.62 |
| 2783 | 8 | 99.76 | 0.06 | 0.05 | 1611.47 | 1917.24 |
| 4642 | 8 | 114.65 | 0.14 | 0.12 | 832.21 | 987.77 |
| 7743 | 8 | 188.54 | 0.21 | 0.18 | 912.02 | 1043.51 |
| 12915 | 8 | 324.56 | 0.36 | 0.32 | 896.77 | 1006.83 |
| 21544 | 8 | 537.28 | 0.68 | 0.61 | 794.57 | 876.22 |
| 35938 | 8 | 987.54 | 1.33 | 1.28 | 741.14 | 769.89 |
| 59948 | 8 | 1736.48 | 2.83 | 2.76 | 612.85 | 628.64 |
| 100000 | 8 | 2767.35 | 6.75 | 6.34 | 410.00 | 436.30 |
| 1000 | 22 | 105.38 | 0.05 | 0.03 | 2265.68 | 3892.12 |
| 1668 | 22 | 162.89 | 0.07 | 0.04 | 2405.72 | 4266.44 |

| | | | | | | |
|---|---|---|---|---|---|---|
| 2783 | 22 | 274.34 | 0.10 | 0.05 | 2648.31 | 5065.34 |
| 4642 | 22 | 315.29 | 0.18 | 0.12 | 1716.93 | 2634.42 |
| 7743 | 22 | 518.49 | 0.29 | 0.19 | 1808.98 | 2670.35 |
| 12915 | 22 | 892.54 | 0.48 | 0.34 | 1863.78 | 2655.99 |
| 21544 | 22 | 1477.53 | 0.85 | 0.64 | 1739.25 | 2298.99 |
| 35938 | 22 | 2715.73 | 1.62 | 1.32 | 1672.84 | 2061.85 |
| 59948 | 22 | 4775.32 | 3.48 | 2.82 | 1371.74 | 1694.21 |
| 100000 | 22 | 7610.22 | 7.25 | 6.66 | 1049.17 | 1142.57 |
| 1000 | 60 | 287.41 | 0.10 | 0.03 | 2859.33 | 8442.54 |
| 1668 | 60 | 444.25 | 0.14 | 0.04 | 3177.82 | 10465.61 |
| 2783 | 60 | 748.19 | 0.20 | 0.06 | 3790.41 | 11845.98 |
| 4642 | 60 | 859.89 | 0.32 | 0.13 | 2691.33 | 6701.67 |
| 7743 | 60 | 1414.07 | 0.50 | 0.21 | 2823.14 | 6729.97 |
| 12915 | 60 | 2434.20 | 0.79 | 0.37 | 3070.63 | 6610.87 |
| 21544 | 60 | 4029.64 | 1.35 | 0.69 | 2991.88 | 5819.03 |
| 35938 | 60 | 7406.55 | 2.37 | 1.41 | 3120.76 | 5255.62 |
| 59948 | 60 | 13023.59 | 4.66 | 2.93 | 2795.13 | 4439.64 |
| 100000 | 60 | 20755.16 | 9.60 | 6.93 | 2163.11 | 2995.45 |
| 1000 | 167 | 799.96 | 0.22 | 0.04 | 3631.45 | 19545.79 |
| 1668 | 167 | 1236.49 | 0.36 | 0.06 | 3462.56 | 21195.78 |
| 2783 | 167 | 2082.47 | 0.46 | 0.08 | 4491.31 | 24962.05 |
| 4642 | 167 | 2393.36 | 0.71 | 0.16 | 3369.18 | 14834.35 |
| 7743 | 167 | 3935.82 | 1.08 | 0.26 | 3630.39 | 15196.79 |
| 12915 | 167 | 6775.19 | 1.66 | 0.44 | 4073.43 | 15465.46 |
| 21544 | 167 | 11215.82 | 2.70 | 0.80 | 4154.85 | 13985.77 |
| 35938 | 167 | 20614.90 | 4.52 | 1.57 | 4557.33 | 13094.95 |
| 59948 | 167 | 36249.00 | 8.06 | 3.30 | 4497.47 | 10993.45 |
| 100000 | 167 | 57768.52 | 15.32 | 7.42 | 3771.74 | 7784.98 |
| 1000 | 464 | 2222.64 | 0.54 | 0.07 | 4092.83 | 33861.25 |
| 1668 | 464 | 3435.52 | 0.88 | 0.09 | 3895.85 | 37054.97 |
| 2783 | 464 | 5786.03 | 1.15 | 0.14 | 5014.51 | 41701.72 |
| 4642 | 464 | 6649.80 | 1.66 | 0.23 | 4015.76 | 28564.07 |
| 7743 | 464 | 10935.45 | 2.68 | 0.35 | 4073.61 | 31080.73 |
| 12915 | 464 | 18824.48 | 4.04 | 0.63 | 4660.96 | 29803.35 |
| 21544 | 464 | 31162.53 | 6.42 | 1.10 | 4852.89 | 28237.42 |
| 35938 | 464 | 57277.31 | 10.41 | 2.08 | 5503.97 | 27511.54 |
| 59948 | 464 | 100715.78 | 17.61 | 4.18 | 5717.78 | 24113.72 |
| 100000 | 464 | 160506.54 | 31.35 | 8.68 | 5119.90 | 18487.07 |
| 1000 | 1292 | 6188.92 | 1.44 | 0.12 | 4296.30 | 50559.81 |

| | | | | | | |
|---|---|---|---|---|---|---|
| 1668 | 1292 | 9566.15 | 2.46 | 0.16 | 3892.82 | 61625.09 |
| 2783 | 1292 | 16111.10 | 2.93 | 0.23 | 5495.04 | 69738.57 |
| 4642 | 1292 | 18516.26 | 4.35 | 0.41 | 4251.95 | 44813.81 |
| 7743 | 1292 | 30449.58 | 7.09 | 0.63 | 4296.67 | 48550.43 |
| 12915 | 1292 | 52416.45 | 10.65 | 1.04 | 4919.44 | 50620.55 |
| 21544 | 1292 | 86771.53 | 16.57 | 1.77 | 5238.05 | 48889.58 |
| 35938 | 1292 | 159487.69 | 26.86 | 3.37 | 5938.26 | 47378.91 |
| 59948 | 1292 | 280441.35 | 43.95 | 6.08 | 6380.93 | 46154.55 |
| 100000 | 1292 | 446927.71 | 74.91 | 12.15 | 5966.33 | 36777.63 |
| 1000 | 3594 | 17215.92 | 3.72 | 0.23 | 4633.61 | 74341.82 |
| 1668 | 3594 | 26610.47 | 6.67 | 0.37 | 3989.33 | 72583.07 |
| 2783 | 3594 | 44816.78 | 7.42 | 0.53 | 6036.00 | 84328.72 |
| 4642 | 3594 | 51507.30 | 12.10 | 0.86 | 4258.52 | 59784.49 |
| 7743 | 3594 | 84702.63 | 19.75 | 1.27 | 4288.88 | 66605.01 |
| 12915 | 3594 | 145808.60 | 29.08 | 2.20 | 5014.57 | 66142.68 |
| 21544 | 3594 | 241375.28 | 44.61 | 3.71 | 5411.39 | 65110.42 |
| 35938 | 3594 | 443652.30 | 71.18 | 6.44 | 6232.68 | 68912.84 |
| 59948 | 3594 | 780113.18 | 114.02 | 10.99 | 6842.13 | 71004.13 |
| 100000 | 3594 | 1243233.88 | 194.97 | 21.52 | 6376.46 | 57776.98 |
| 1000 | 10000 | 47901.83 | 9.91 | 0.54 | 4833.40 | 87941.03 |
| 1668 | 10000 | 74041.39 | 18.23 | 0.82 | 4061.66 | 90209.78 |
| 2783 | 10000 | 124698.88 | 20.41 | 1.33 | 6110.11 | 93659.04 |
| 4642 | 10000 | 143314.69 | 32.77 | 2.11 | 4373.27 | 67996.47 |
| 7743 | 10000 | 235677.87 | 52.07 | 3.38 | 4526.25 | 69805.32 |
| 12915 | 10000 | 405700.05 | 77.94 | 5.51 | 5205.26 | 73661.58 |
| 21544 | 10000 | 671606.24 | 120.29 | 9.18 | 5583.38 | 73178.83 |
| 35938 | 10000 | 1234424.87 | 193.62 | 15.42 | 6375.42 | 80061.05 |
| 59948 | 10000 | 2170598.72 | 314.46 | 26.28 | 6902.55 | 82588.13 |
| 100000 | 10000 | 3459192.78 | 520.51 | 46.46 | 6645.74 | 74454.15 |

#### Simulation 3: comparison of carbon footprint with lmer

**Table S9: Comparison of carbon footprint (in grams of carbon dioxide emission) of lmer and FEMA.** The lmer carbon footprint are extrapolated from the carbon footprint for a single outcome variable; the green ratio is the ratio of carbon footprint of lmer to the carbon footprint of FEMA

| No. of observations | No. of *y* vars. | Carbon footprint lmer ($gCO_2e$) | Carbon footprint FEMA (serial; $gCO_2e$) | Carbon footprint FEMA (parallel; $gCO_2e$) | FEMA green ratio (serial) | FEMA green ratio (parallel) |
|---|---|---|---|---|---|---|
| 1000 | 1 | 15.64 | 0.03 | 0.03 | 621.61 | 457.66 |
| 1668 | 1 | 8.52 | 0.03 | 0.03 | 300.48 | 290.82 |
| 2783 | 1 | 10.82 | 0.05 | 0.05 | 236.16 | 235.40 |
| 4642 | 1 | 22.01 | 0.09 | 0.10 | 235.73 | 214.46 |
| 7743 | 1 | 31.25 | 0.16 | 0.18 | 197.90 | 176.33 |
| 12915 | 1 | 60.51 | 0.29 | 0.32 | 207.23 | 191.22 |
| 21544 | 1 | 104.08 | 0.57 | 0.60 | 183.70 | 173.06 |
| 35938 | 1 | 221.38 | 1.17 | 1.25 | 188.93 | 177.15 |
| 59948 | 1 | 324.70 | 2.60 | 2.74 | 125.11 | 118.51 |
| 100000 | 1 | 567.00 | 6.35 | 6.58 | 89.23 | 86.14 |
| 1000 | 3 | 46.93 | 0.02 | 0.03 | 2051.38 | 1785.96 |
| 1668 | 3 | 25.55 | 0.03 | 0.03 | 753.43 | 788.04 |
| 2783 | 3 | 32.46 | 0.05 | 0.05 | 629.08 | 626.99 |
| 4642 | 3 | 66.04 | 0.11 | 0.11 | 613.11 | 601.76 |
| 7743 | 3 | 93.74 | 0.18 | 0.18 | 523.86 | 529.37 |
| 12915 | 3 | 181.52 | 0.31 | 0.32 | 579.56 | 567.52 |
| 21544 | 3 | 312.24 | 0.62 | 0.62 | 507.37 | 499.62 |
| 35938 | 3 | 664.15 | 1.24 | 1.28 | 536.09 | 518.20 |
| 59948 | 3 | 974.09 | 2.86 | 2.92 | 340.22 | 333.82 |
| 100000 | 3 | 1701.01 | 6.46 | 6.57 | 263.33 | 258.95 |
| 1000 | 8 | 125.15 | 0.03 | 0.03 | 3985.34 | 4879.56 |
| 1668 | 8 | 68.14 | 0.05 | 0.04 | 1492.86 | 1924.22 |
| 2783 | 8 | 86.57 | 0.06 | 0.05 | 1398.37 | 1663.70 |
| 4642 | 8 | 176.10 | 0.14 | 0.12 | 1278.21 | 1517.14 |
| 7743 | 8 | 249.98 | 0.21 | 0.18 | 1209.22 | 1383.57 |
| 12915 | 8 | 484.04 | 0.36 | 0.32 | 1337.42 | 1501.57 |
| 21544 | 8 | 832.65 | 0.68 | 0.61 | 1231.38 | 1357.92 |
| 35938 | 8 | 1771.08 | 1.33 | 1.28 | 1329.18 | 1380.74 |
| 59948 | 8 | 2597.59 | 2.83 | 2.76 | 916.76 | 940.37 |
| 100000 | 8 | 4536.02 | 6.75 | 6.34 | 672.04 | 715.14 |
| 1000 | 22 | 344.15 | 0.05 | 0.03 | 7399.07 | 12710.57 |
| 1668 | 22 | 187.39 | 0.07 | 0.04 | 2767.60 | 4908.22 |
| 2783 | 22 | 238.06 | 0.10 | 0.05 | 2298.09 | 4395.50 |

| | | | | | | |
|---|---|---|---|---|---|---|
| 4642 | 22 | 484.26 | 0.18 | 0.12 | 2637.05 | 4046.24 |
| 7743 | 22 | 687.46 | 0.29 | 0.19 | 2398.49 | 3540.56 |
| 12915 | 22 | 1331.11 | 0.48 | 0.34 | 2779.61 | 3961.09 |
| 21544 | 22 | 2289.80 | 0.85 | 0.64 | 2695.39 | 3562.84 |
| 35938 | 22 | 4870.46 | 1.62 | 1.32 | 3000.11 | 3697.76 |
| 59948 | 22 | 7143.36 | 3.48 | 2.82 | 2051.97 | 2534.35 |
| 100000 | 22 | 12474.06 | 7.25 | 6.66 | 1719.72 | 1872.81 |
| 1000 | 60 | 938.60 | 0.10 | 0.03 | 9337.76 | 27570.93 |
| 1668 | 60 | 511.07 | 0.14 | 0.04 | 3655.84 | 12039.91 |
| 2783 | 60 | 649.25 | 0.20 | 0.06 | 3289.16 | 10279.47 |
| 4642 | 60 | 1320.71 | 0.32 | 0.13 | 4133.65 | 10293.20 |
| 7743 | 60 | 1874.88 | 0.50 | 0.21 | 3743.14 | 8923.13 |
| 12915 | 60 | 3630.31 | 0.79 | 0.37 | 4579.46 | 9859.31 |
| 21544 | 60 | 6244.90 | 1.35 | 0.69 | 4636.64 | 9018.00 |
| 35938 | 60 | 13283.06 | 2.37 | 1.41 | 5596.83 | 9425.53 |
| 59948 | 60 | 19481.90 | 4.66 | 2.93 | 4181.21 | 6641.22 |
| 100000 | 60 | 34020.16 | 9.60 | 6.93 | 3545.60 | 4909.89 |
| 1000 | 167 | 2612.44 | 0.22 | 0.04 | 11859.27 | 63831.01 |
| 1668 | 167 | 1422.49 | 0.36 | 0.06 | 3983.42 | 24384.16 |
| 2783 | 167 | 1807.09 | 0.46 | 0.08 | 3897.38 | 21661.08 |
| 4642 | 167 | 3675.99 | 0.71 | 0.16 | 5174.76 | 22784.29 |
| 7743 | 167 | 5218.42 | 1.08 | 0.26 | 4813.45 | 20149.11 |
| 12915 | 167 | 10104.37 | 1.66 | 0.44 | 6075.03 | 23064.85 |
| 21544 | 167 | 17381.63 | 2.70 | 0.80 | 6438.94 | 21674.33 |
| 35938 | 167 | 36971.19 | 4.52 | 1.57 | 8173.22 | 23484.76 |
| 59948 | 167 | 54224.61 | 8.06 | 3.30 | 6727.73 | 16445.03 |
| 100000 | 167 | 94689.45 | 15.32 | 7.42 | 6182.32 | 12760.50 |
| 1000 | 464 | 7258.53 | 0.54 | 0.07 | 13366.02 | 110581.23 |
| 1668 | 464 | 3952.31 | 0.88 | 0.09 | 4481.89 | 42628.98 |
| 2783 | 464 | 5020.88 | 1.15 | 0.14 | 4351.39 | 36187.09 |
| 4642 | 464 | 10213.52 | 1.66 | 0.23 | 6167.86 | 43871.96 |
| 7743 | 464 | 14499.09 | 2.68 | 0.35 | 5401.12 | 41209.30 |
| 12915 | 464 | 28074.42 | 4.04 | 0.63 | 6951.25 | 44448.06 |
| 21544 | 464 | 48293.87 | 6.42 | 1.10 | 7520.73 | 43760.70 |
| 35938 | 464 | 102722.36 | 10.41 | 2.08 | 9870.94 | 49339.79 |
| 59948 | 464 | 150660.00 | 17.61 | 4.18 | 8553.18 | 36071.54 |
| 100000 | 464 | 263089.26 | 31.35 | 8.68 | 8392.12 | 30302.51 |
| 1000 | 1292 | 20211.24 | 1.44 | 0.12 | 14030.49 | 165114.00 |
| 1668 | 1292 | 11005.14 | 2.46 | 0.16 | 4478.40 | 70895.07 |

| | | | | | | |
|---|---|---|---|---|---|---|
| 2783 | 1292 | 13980.57 | 2.93 | 0.23 | 4768.37 | 60516.35 |
| 4642 | 1292 | 28439.38 | 4.35 | 0.41 | 6530.62 | 68830.16 |
| 7743 | 1292 | 40372.47 | 7.09 | 0.63 | 5696.87 | 64372.01 |
| 12915 | 1292 | 78172.73 | 10.65 | 1.04 | 7336.74 | 75494.36 |
| 21544 | 1292 | 134473.44 | 16.57 | 1.77 | 8117.62 | 75766.21 |
| 35938 | 1292 | 286028.63 | 26.86 | 3.37 | 10649.81 | 84970.35 |
| 59948 | 1292 | 419510.18 | 43.95 | 6.08 | 9545.19 | 69042.25 |
| 100000 | 1292 | 732567.51 | 74.91 | 12.15 | 9779.52 | 60282.90 |
| 1000 | 3594 | 56222.30 | 3.72 | 0.23 | 15132.06 | 242779.27 |
| 1668 | 3594 | 30613.37 | 6.67 | 0.37 | 4589.43 | 83501.41 |
| 2783 | 3594 | 38890.21 | 7.42 | 0.53 | 5237.80 | 73177.10 |
| 4642 | 3594 | 79110.78 | 12.10 | 0.86 | 6540.72 | 91823.83 |
| 7743 | 3594 | 112305.46 | 19.75 | 1.27 | 5686.53 | 88310.21 |
| 12915 | 3594 | 217455.72 | 29.08 | 2.20 | 7478.62 | 98643.73 |
| 21544 | 3594 | 374069.30 | 44.61 | 3.71 | 8386.26 | 100904.32 |
| 35938 | 3594 | 795655.48 | 71.18 | 6.44 | 11177.82 | 123589.76 |
| 59948 | 3594 | 1166965.62 | 114.02 | 10.99 | 10235.10 | 106214.56 |
| 100000 | 3594 | 2037807.75 | 194.97 | 21.52 | 10451.78 | 94703.33 |
| 1000 | 10000 | 156433.77 | 9.91 | 0.54 | 15784.50 | 287190.45 |
| 1668 | 10000 | 85179.10 | 18.23 | 0.82 | 4672.64 | 103779.62 |
| 2783 | 10000 | 108208.72 | 20.41 | 1.33 | 5302.11 | 81273.58 |
| 4642 | 10000 | 220119.04 | 32.77 | 2.11 | 6716.96 | 104436.73 |
| 7743 | 10000 | 312480.40 | 52.07 | 3.38 | 6001.27 | 92553.43 |
| 12915 | 10000 | 605052.08 | 77.94 | 5.51 | 7763.02 | 109857.25 |
| 21544 | 10000 | 1040816.09 | 120.29 | 9.18 | 8652.80 | 113408.27 |
| 35938 | 10000 | 2213843.86 | 193.62 | 15.42 | 11433.81 | 143583.19 |
| 59948 | 10000 | 3246982.80 | 314.46 | 26.28 | 10325.47 | 123542.98 |
| 100000 | 10000 | 5670027.14 | 520.51 | 46.46 | 10893.15 | 122039.18 |

#### Simulation 4: comparison of estimated parameters with ground truth

##### Sample size: 12,000

*Minimum number of observations: 500*

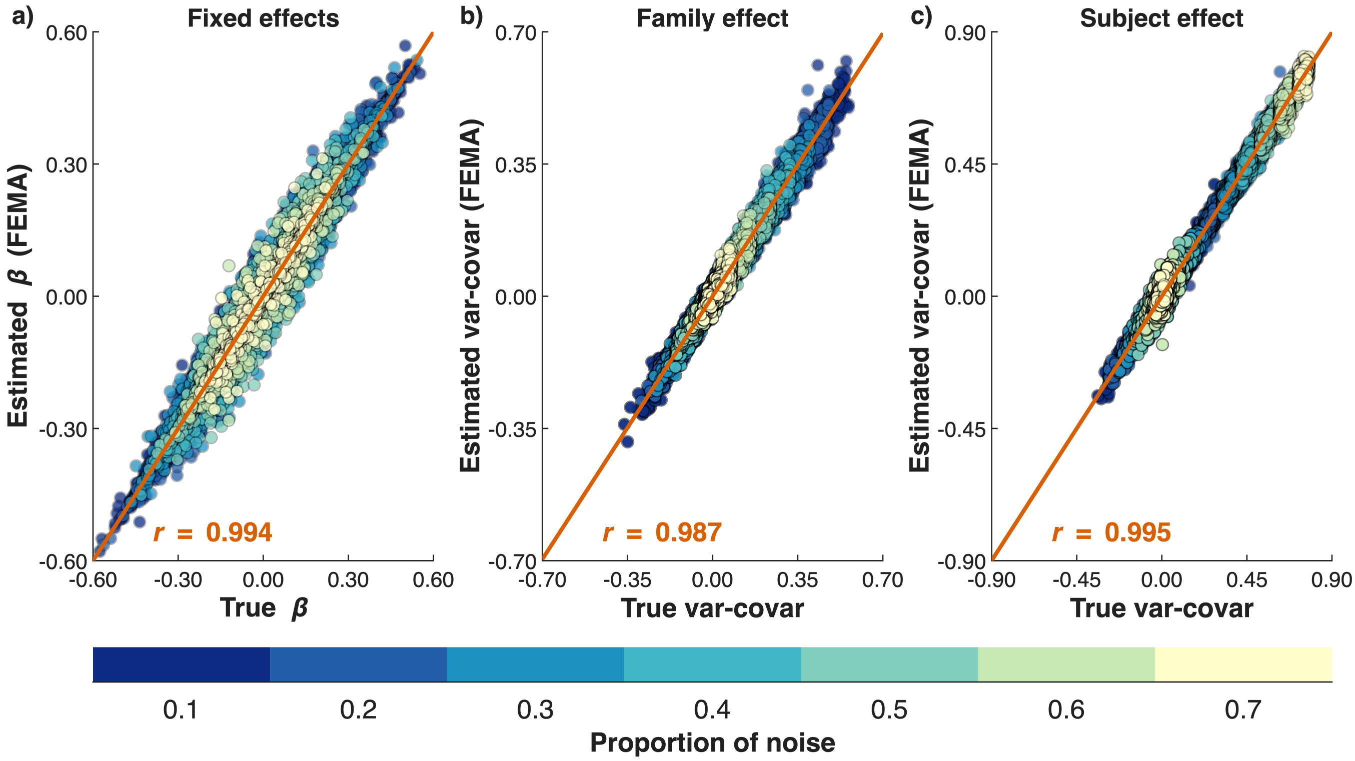


**Figure S10**: **Comparison of estimated parameters from FEMA with ground truth.** Scatterplots of estimated parameters against ground truth across 50 iterations and 84 simulation settings ($n_{\text{obs}} = 12{,}000$; $\min_{\text{numObs}} = 500$).

*Minimum number of observations: 600*

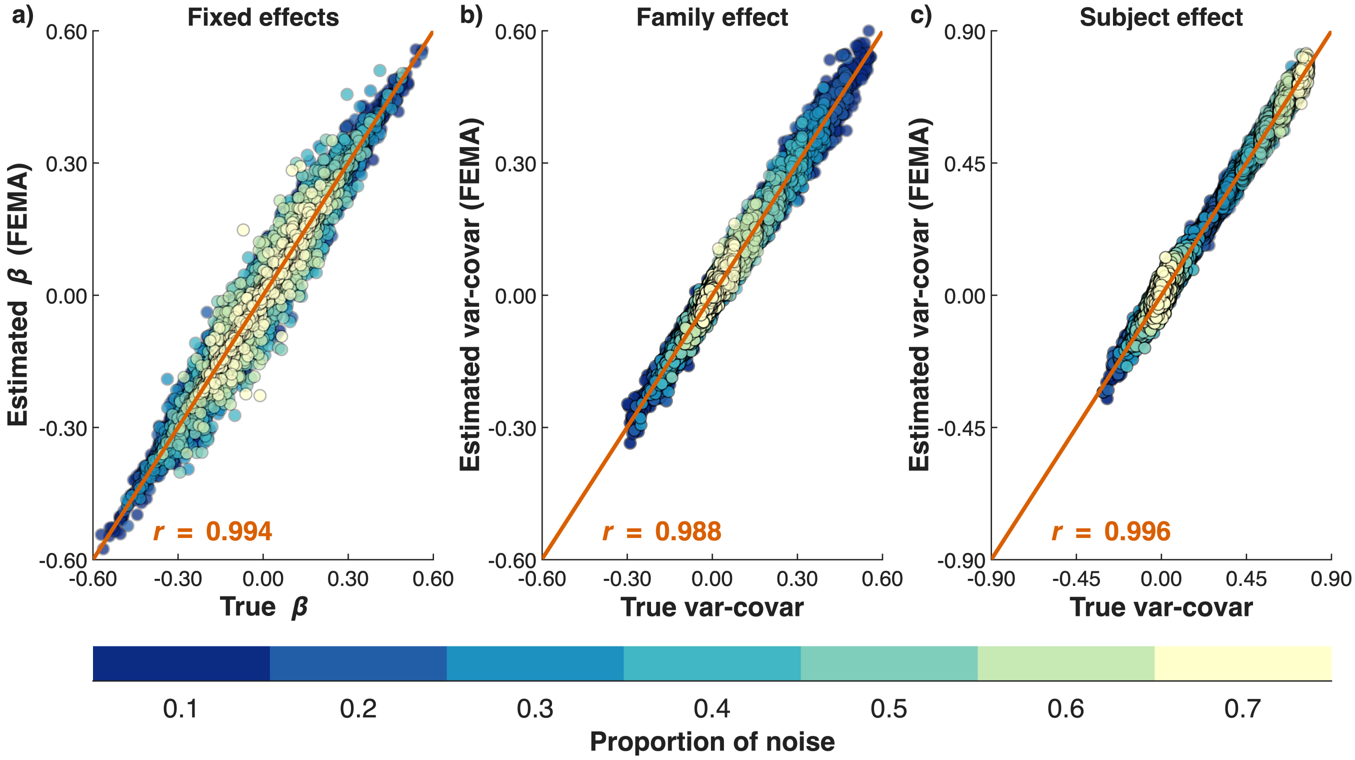


**Figure S11**: **Comparison of estimated parameters from FEMA with ground truth.** Scatterplots of estimated parameters against ground truth across 50 iterations and 84 simulation settings ($n_{\text{obs}} = 12{,}000$; $\text{min}_{\text{numObs}} = 600$).

*Minimum number of observations: 700*

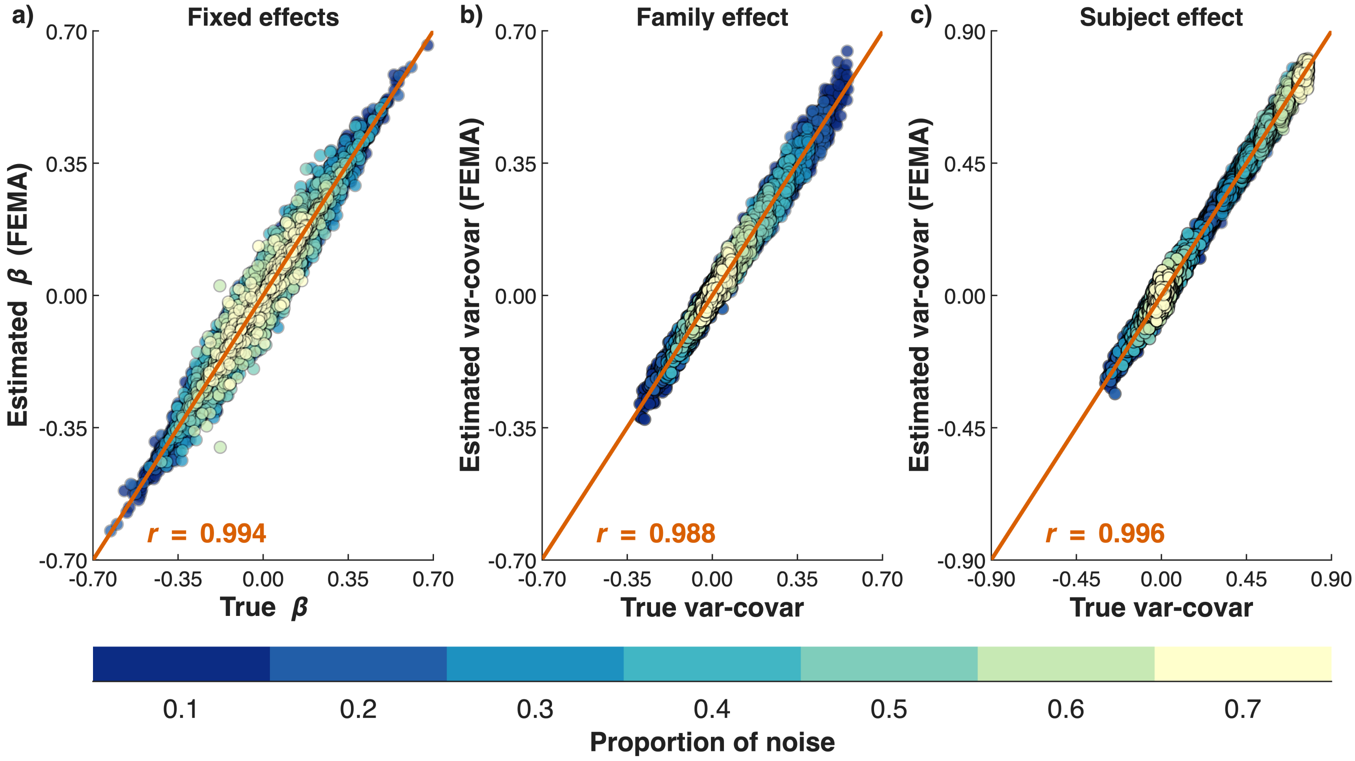


**Figure S12**: **Comparison of estimated parameters from FEMA with ground truth.** Scatterplots of estimated parameters against ground truth across 50 iterations and 84 simulation settings ($n_{\text{obs}} = 12{,}000$; $\text{min}_{\text{numObs}} = 700$).

*Minimum number of observations: 800*

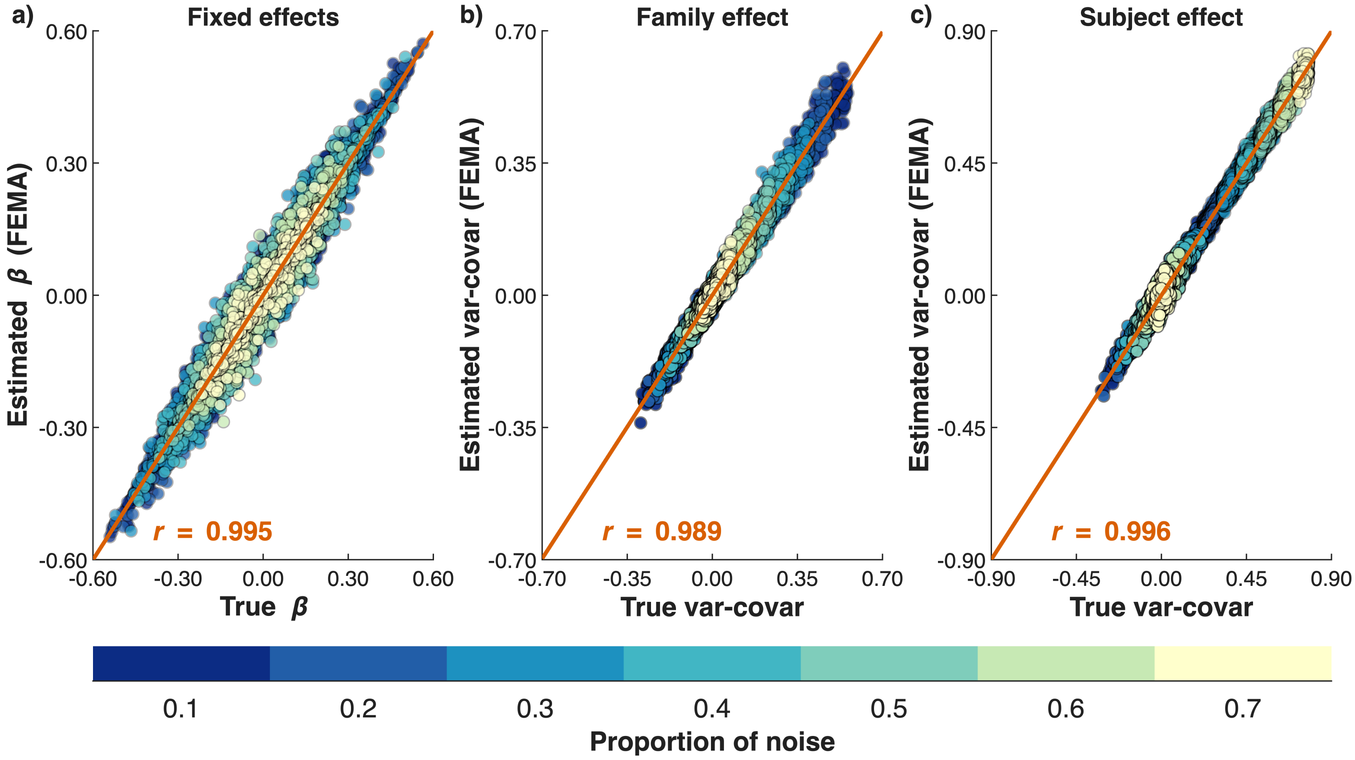


**Figure S13**: **Comparison of estimated parameters from FEMA with ground truth.** Scatterplots of estimated parameters against ground truth across 50 iterations and 84 simulation settings ($n_{\text{obs}} = 12{,}000$; $\text{min}_{\text{numObs}} = 700$).

#### Sample size: 15,000

*Minimum number of observations: 500*

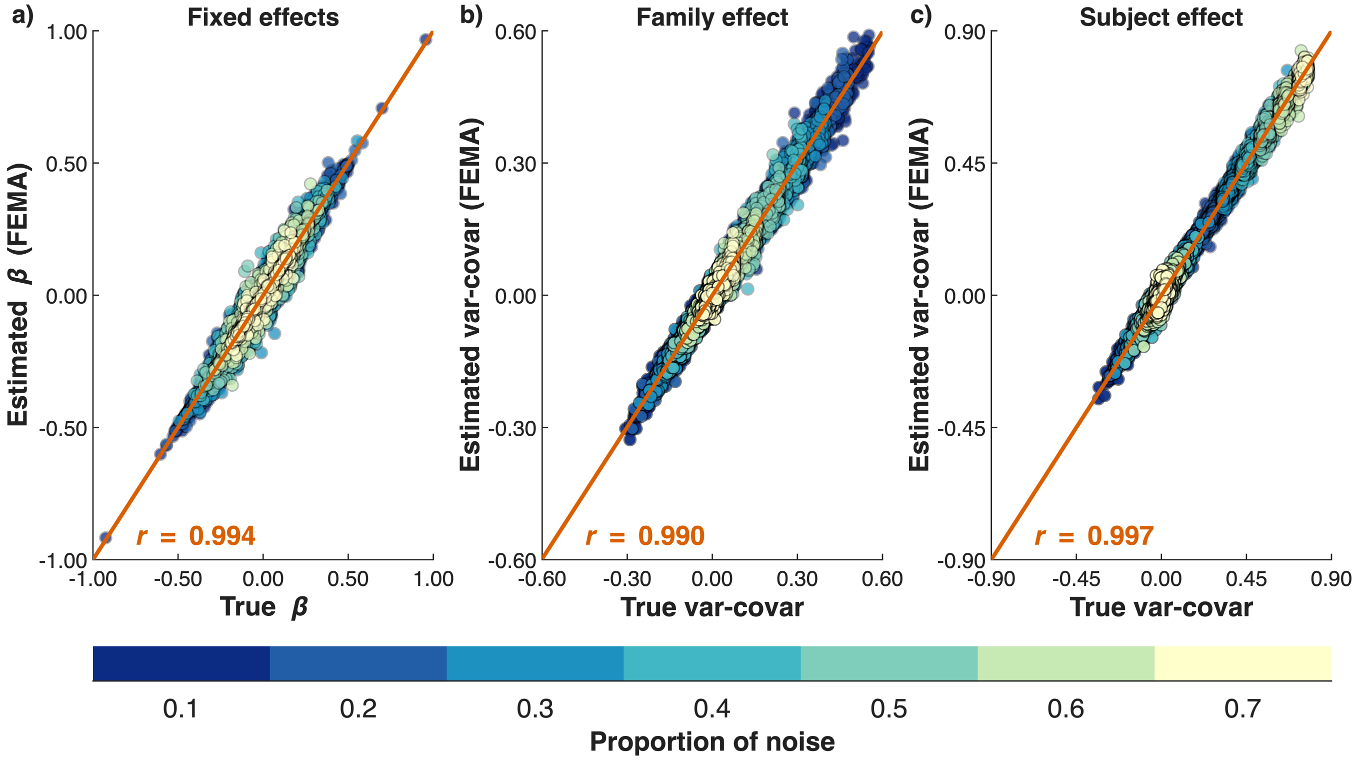


**Figure S14**: **Comparison of estimated parameters from FEMA with ground truth.** Scatterplots of estimated parameters against ground truth across 50 iterations and 84 simulation settings ($n_{\text{obs}} = 15{,}000$; $\min_{\text{numObs}} = 500$).

*Minimum number of observations: 600*

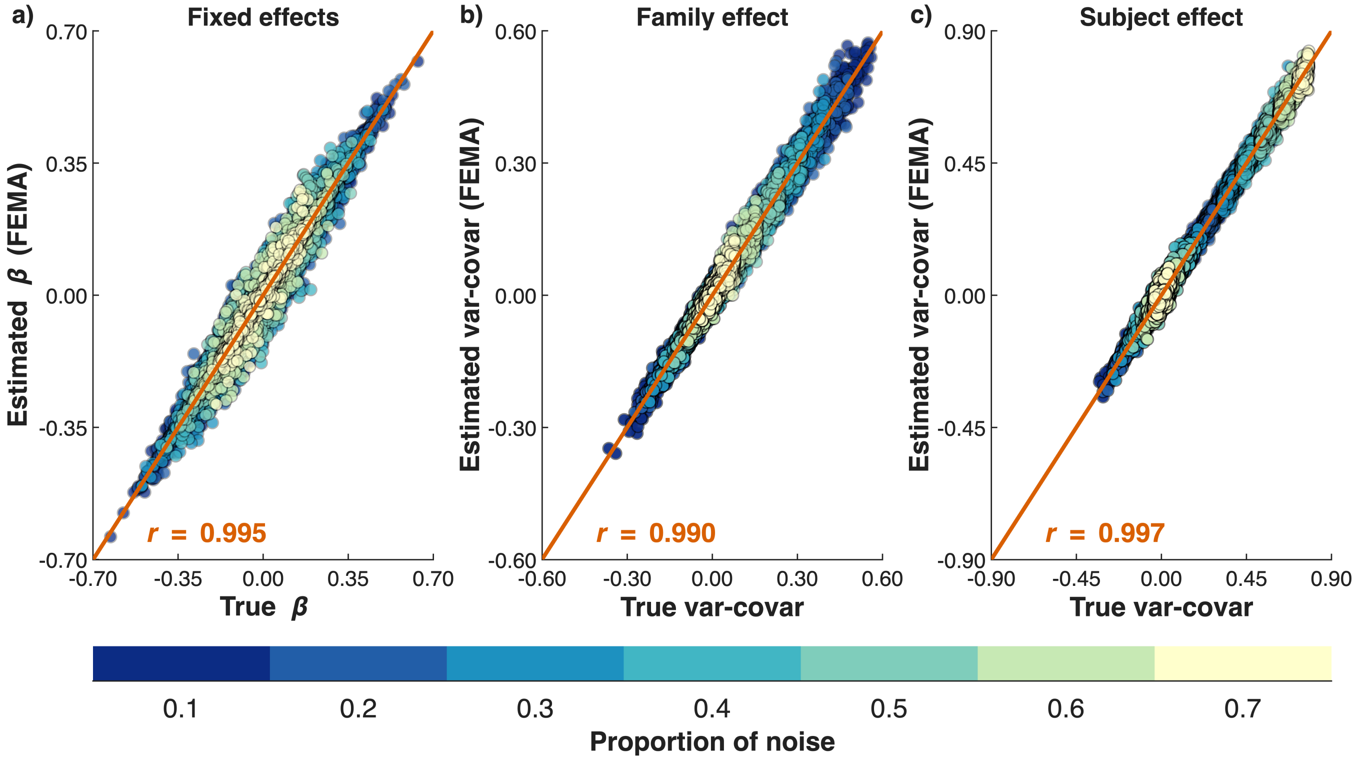


**Figure S15**: **Comparison of estimated parameters from FEMA with ground truth.** Scatterplots of estimated parameters against ground truth across 50 iterations and 84 simulation settings ($n_{\text{obs}} = 15{,}000$; $\min_{\text{numObs}} = 600$).

*Minimum number of observations: 700*

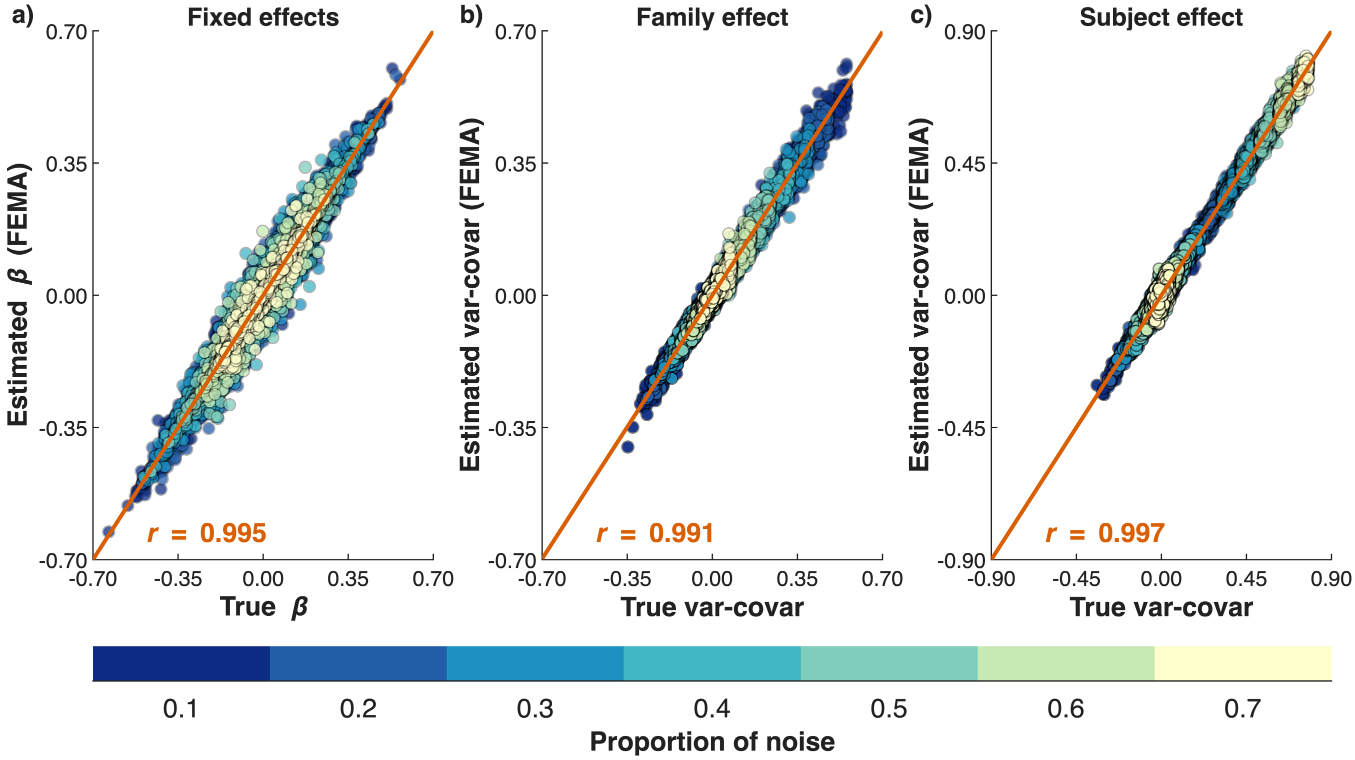


**Figure S16**: **Comparison of estimated parameters from FEMA with ground truth.** Scatterplots of estimated parameters against ground truth across 50 iterations and 84 simulation settings ($n_{\text{obs}} = 15{,}000$; $\text{min}_{\text{numObs}} = 700$).

*Minimum number of observations: 800*

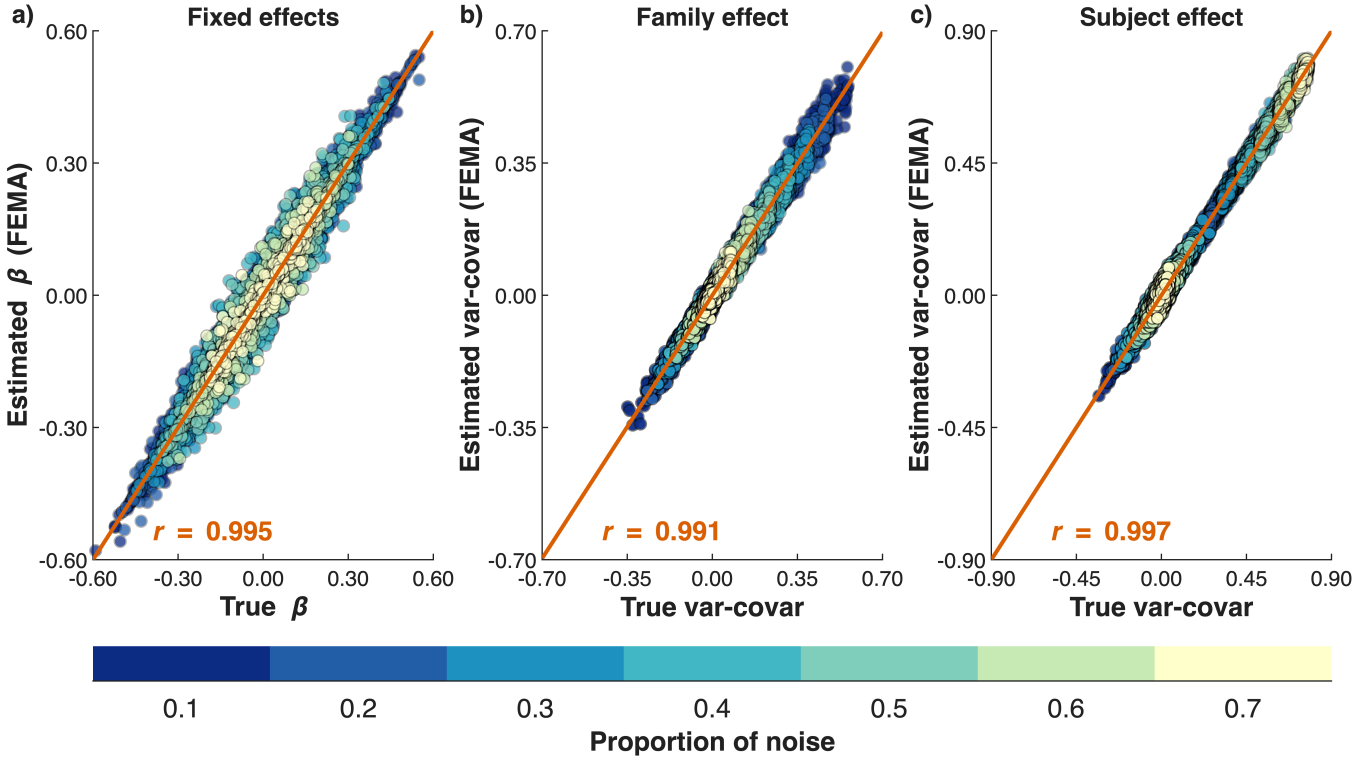


**Figure S17**: **Comparison of estimated parameters from FEMA with ground truth.** Scatterplots of estimated parameters against ground truth across 50 iterations and 84 simulation settings ($n_{\text{obs}} = 15{,}000$; $\text{min}_{\text{numObs}} = 800$).

**Sample size: 18,000**

*Minimum number of observations: 500*

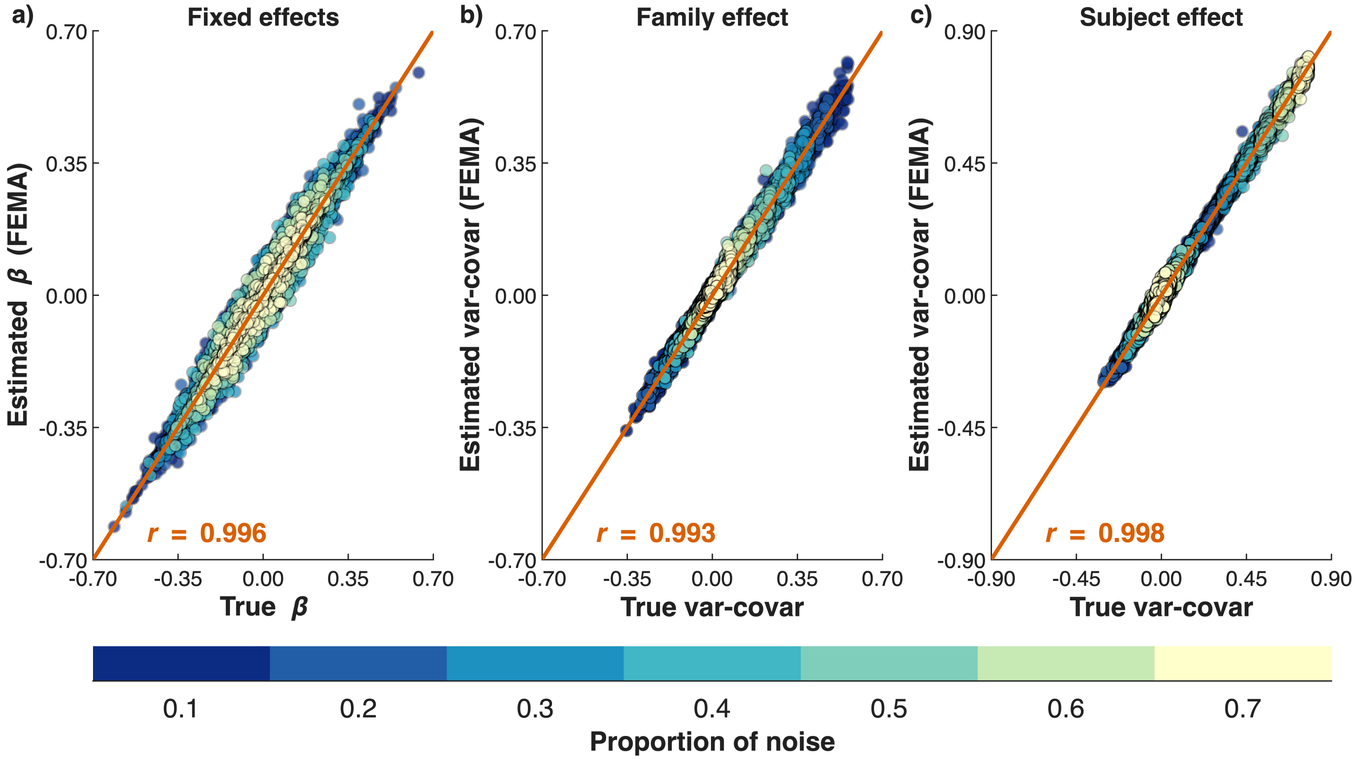


**Figure S18**: **Comparison of estimated parameters from FEMA with ground truth.** Scatterplots of estimated parameters against ground truth across 50 iterations and 84 simulation settings ($n_{\mathrm{obs}}$ = 18,000; $\mathrm{min}_{\mathrm{numObs}}$ = 500).

*Minimum number of observations: 600*

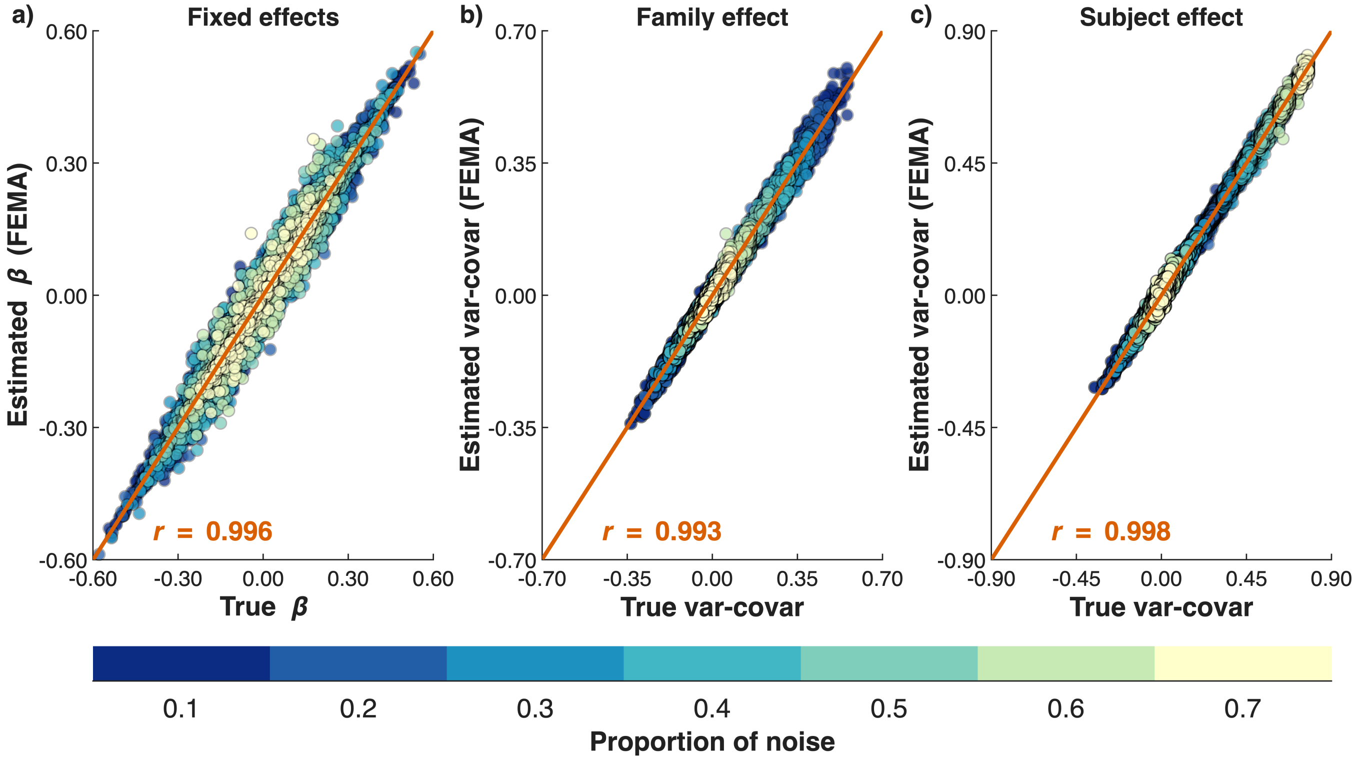


**Figure S19**: **Comparison of estimated parameters from FEMA with ground truth.** Scatterplots of estimated parameters against ground truth across 50 iterations and 84 simulation settings ($n_{\mathrm{obs}}$ = 18,000; $\mathrm{min}_{\mathrm{numObs}}$ = 600).

*Minimum number of observations: 700*

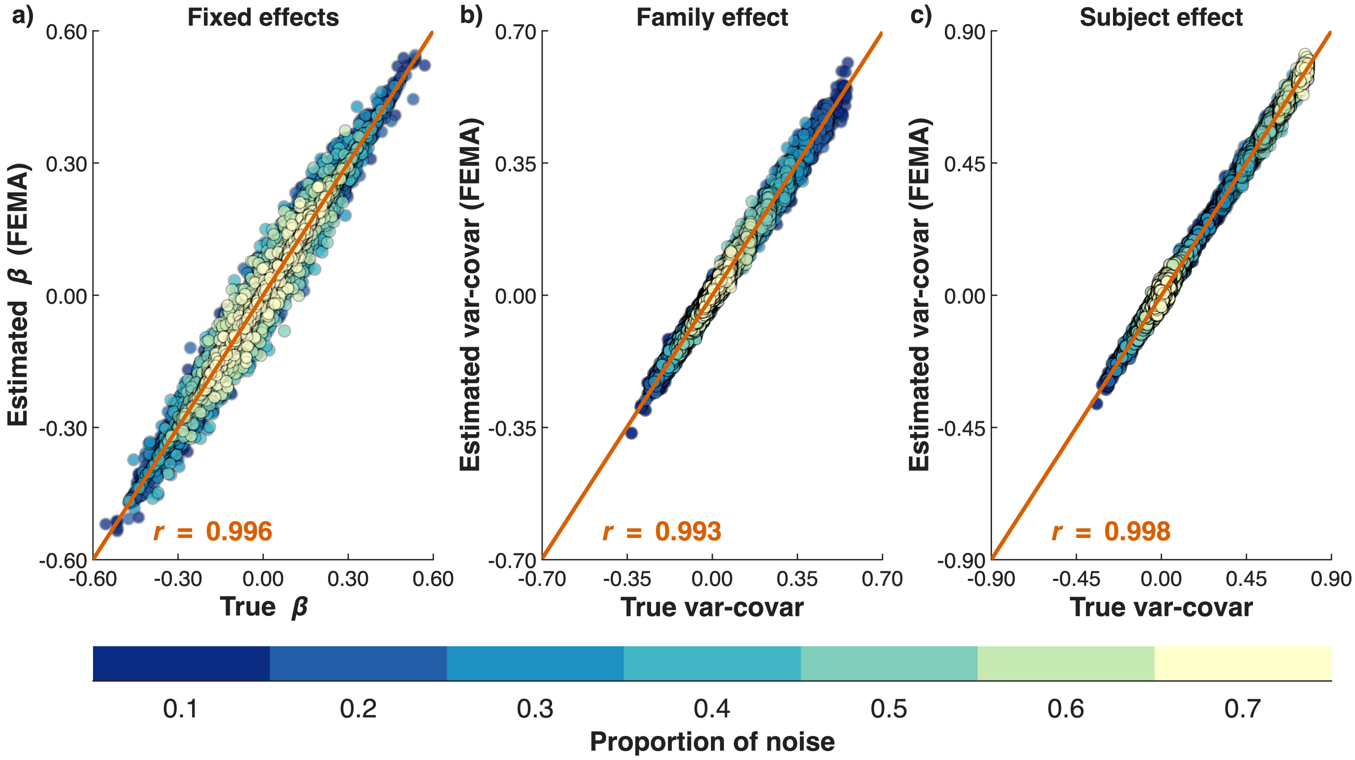


**Figure S20**: **Comparison of estimated parameters from FEMA with ground truth.** Scatterplots of estimated parameters against ground truth across 50 iterations and 84 simulation settings ($n_{\text{obs}}$ = 18,000; $\text{min}_{\text{numObs}}$ = 700).

*Minimum number of observations: 800*

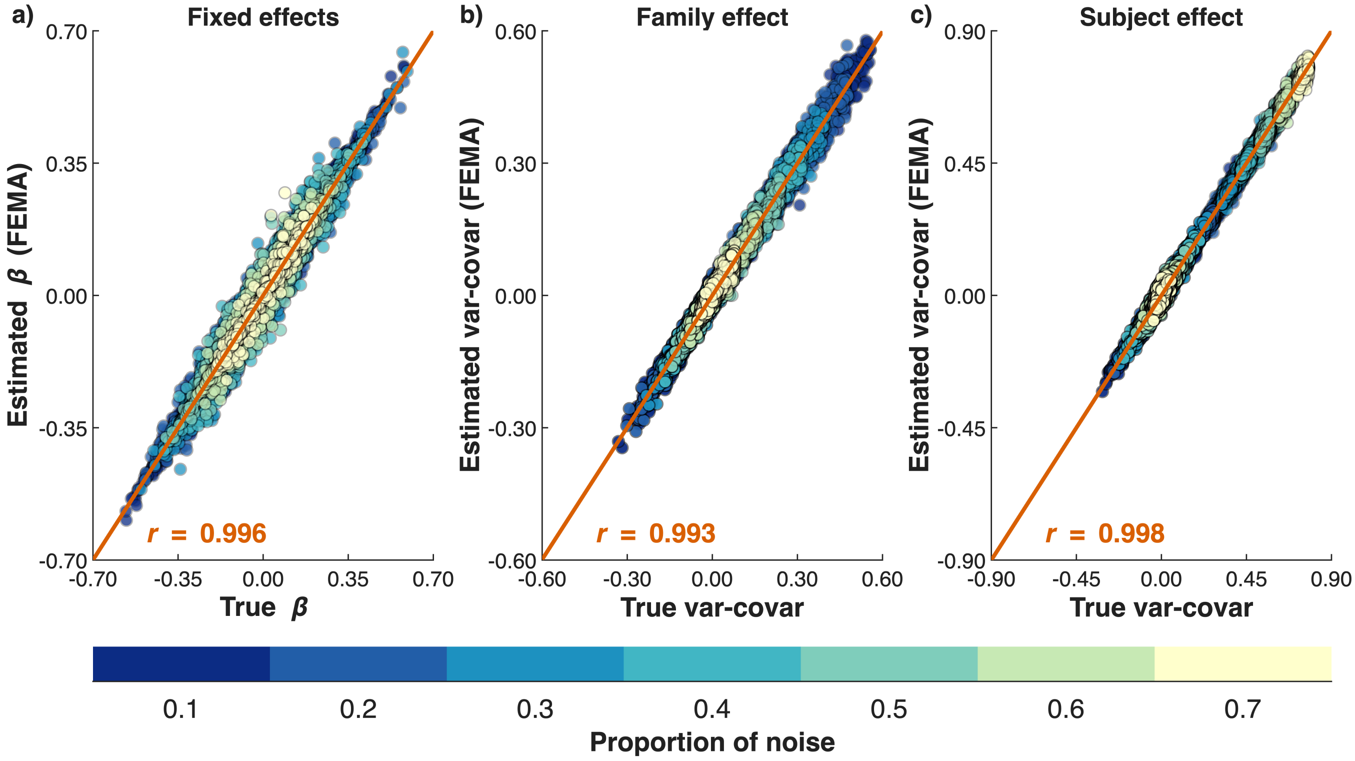


**Figure S21**: **Comparison of estimated parameters from FEMA with ground truth.** Scatterplots of estimated parameters against ground truth across 50 iterations and 84 simulation settings ($n_{\text{obs}}$ = 18,000; $\text{min}_{\text{numObs}}$ = 800).

**Sample size: 20,000**

*Minimum number of observations: 500*

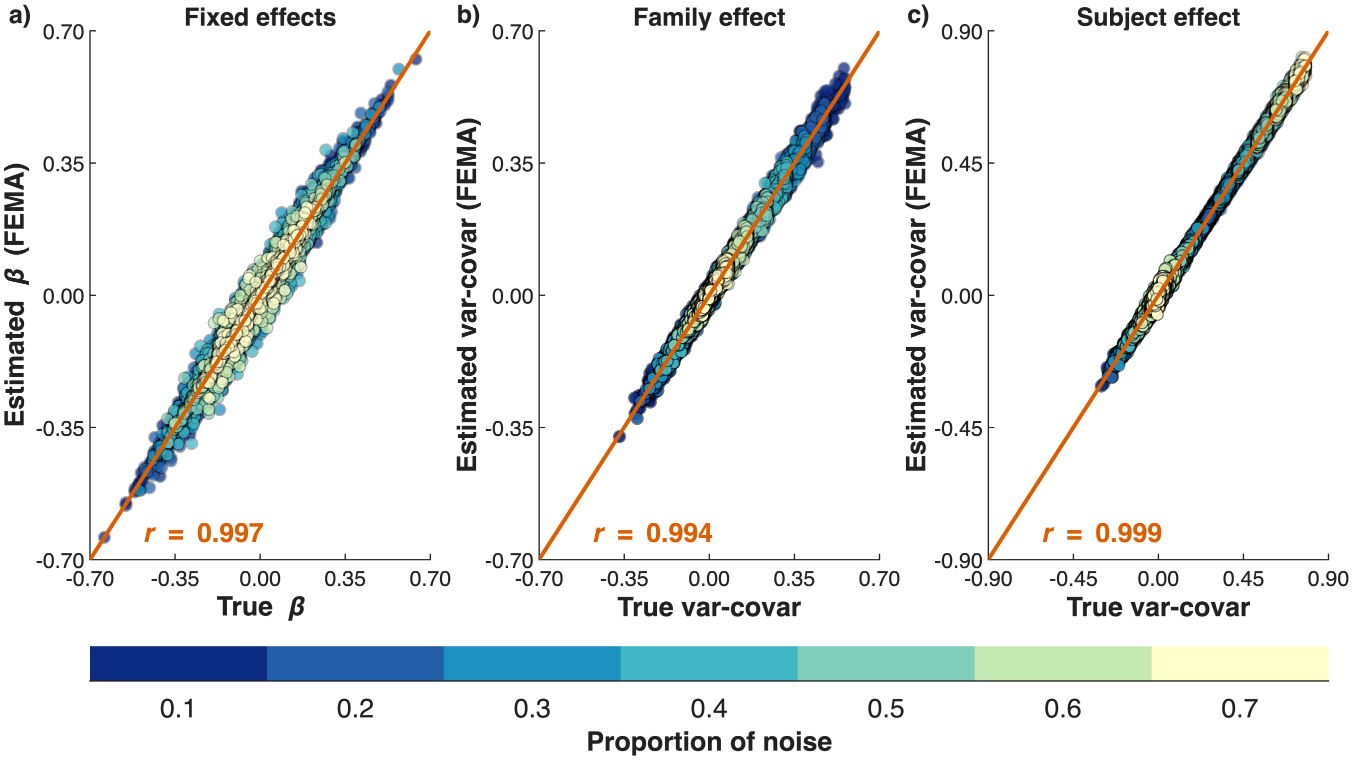


**Figure S22**: **Comparison of estimated parameters from FEMA with ground truth.** Scatterplots of estimated parameters against ground truth across 50 iterations and 84 simulation settings ($n_{obs}$ = 20,000; $min_{numObs}$ = 500).

*Minimum number of observations: 600*

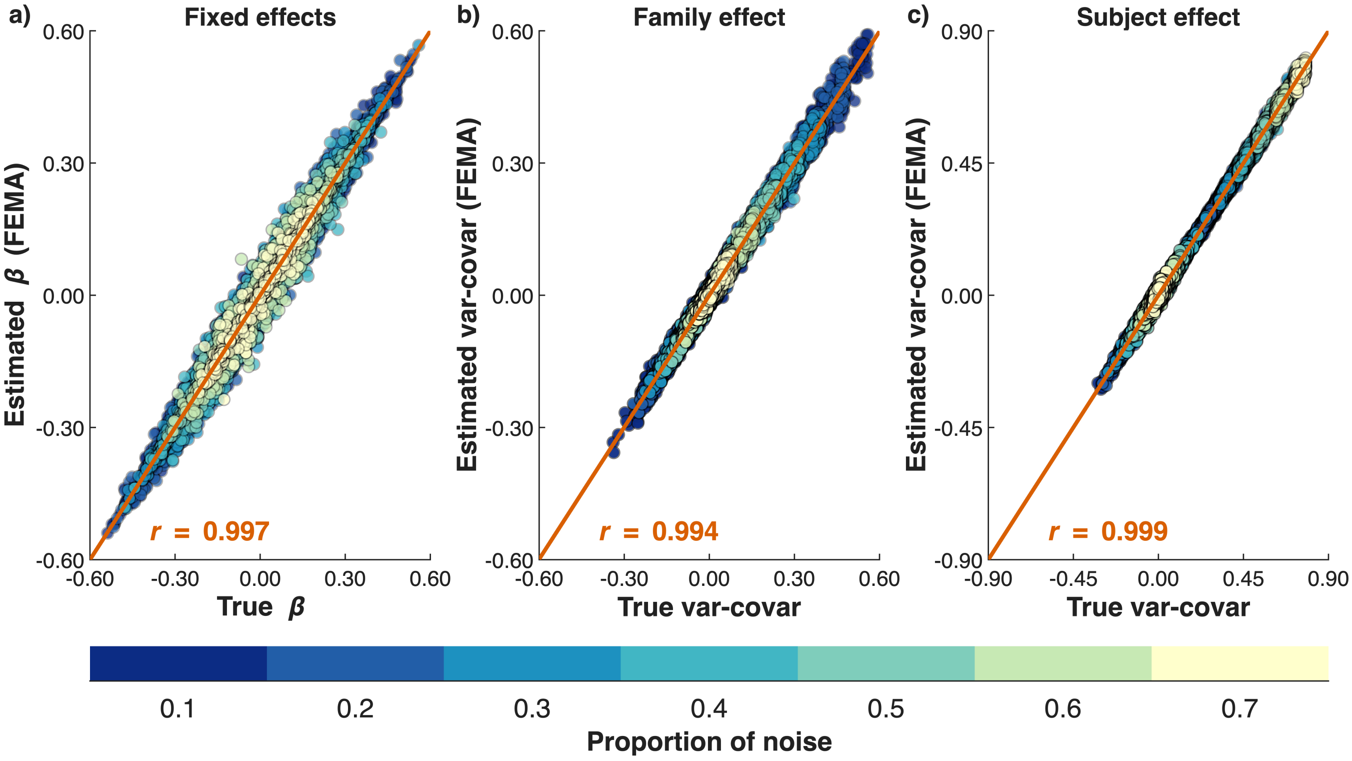


**Figure S23**: **Comparison of estimated parameters from FEMA with ground truth.** Scatterplots of estimated parameters against ground truth across 50 iterations and 84 simulation settings ($n_{obs}$ = 20,000; $min_{numObs}$ = 600).

*Minimum number of observations: 700*

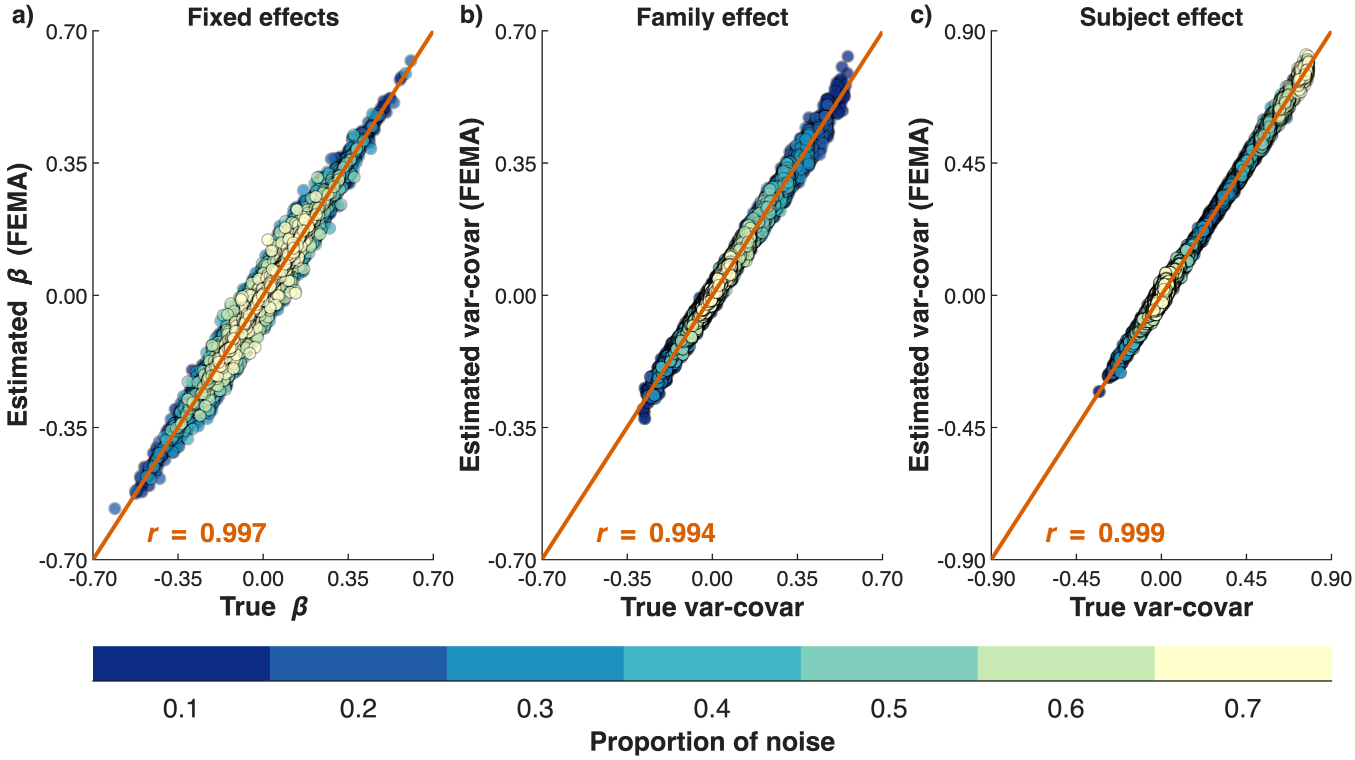


**Figure S24**: **Comparison of estimated parameters from FEMA with ground truth.** Scatterplots of estimated parameters against ground truth across 50 iterations and 84 simulation settings ($n_{\text{obs}} = 20{,}000$; $\text{min}_{\text{numObs}} = 700$).

*Minimum number of observations: 800*

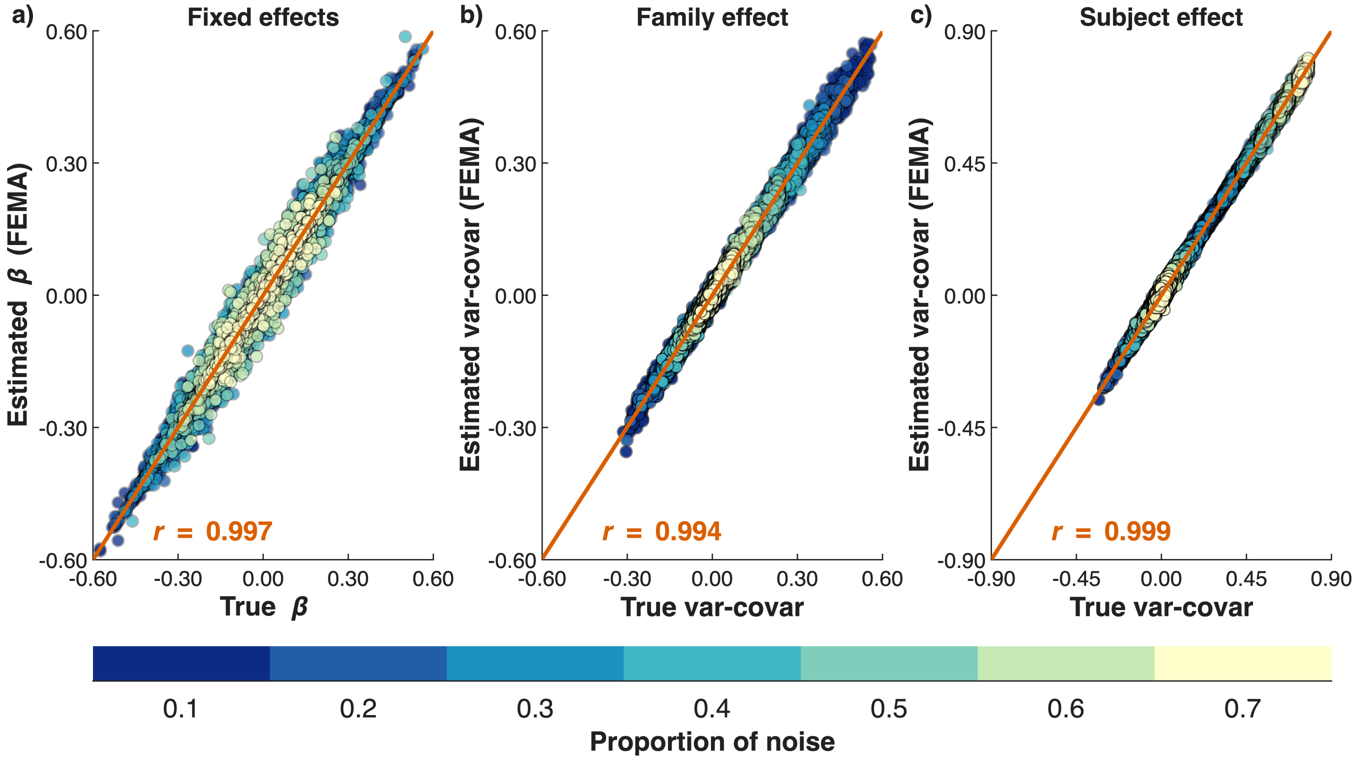


**Figure S25**: **Comparison of estimated parameters from FEMA with ground truth.** Scatterplots of estimated parameters against ground truth across 50 iterations and 84 simulation settings ($n_{\text{obs}} = 20{,}000$; $\text{min}_{\text{numObs}} = 800$).

#### Simulation 4: summary of comparison of estimated parameters

##### Comparison with ground truth

**Table S10**: **Summary of differences between the ground truth and the estimated parameters from FEMA**. Mean and standard deviation (across 84 experimental conditions) of the average (across 50 repeats) root mean squared error (RMSE) between ground truth and estimated coefficients from FEMA for differing total number of observations and minimum number of observations between pairs of visits

| $n_{Obs}$ | $\min_{numObs}$ | RMSE beta: truth – FEMA | RMSE family: truth – FEMA | RMSE subject: truth – FEMA |
|---|---|---|---|---|
| 12000 | 500 | 0.0150 ± 0.0025 | 0.0128 ± 0.0033 | 0.0152 ± 0.0042 |
| 12000 | 600 | 0.0145 ± 0.0025 | 0.0125 ± 0.0033 | 0.0148 ± 0.0042 |
| 12000 | 700 | 0.0143 ± 0.0024 | 0.0123 ± 0.0032 | 0.0145 ± 0.0040 |
| 12000 | 800 | 0.0140 ± 0.0023 | 0.0121 ± 0.0031 | 0.0143 ± 0.0040 |
| 15000 | 500 | 0.0140 ± 0.0024 | 0.0111 ± 0.0030 | 0.0124 ± 0.0036 |
| 15000 | 600 | 0.0139 ± 0.0024 | 0.0110 ± 0.0029 | 0.0122 ± 0.0036 |
| 15000 | 700 | 0.0138 ± 0.0022 | 0.0109 ± 0.0030 | 0.0120 ± 0.0035 |
| 15000 | 800 | 0.0133 ± 0.0021 | 0.0108 ± 0.0030 | 0.0118 ± 0.0034 |
| 18000 | 500 | 0.0124 ± 0.0020 | 0.0094 ± 0.0026 | 0.0096 ± 0.0030 |
| 18000 | 600 | 0.0124 ± 0.0021 | 0.0094 ± 0.0027 | 0.0096 ± 0.0029 |
| 18000 | 700 | 0.0124 ± 0.0020 | 0.0094 ± 0.0026 | 0.0096 ± 0.0030 |
| 18000 | 800 | 0.0123 ± 0.0021 | 0.0094 ± 0.0027 | 0.0095 ± 0.0030 |
| 20000 | 500 | 0.0111 ± 0.0018 | 0.0085 ± 0.0026 | 0.0082 ± 0.0027 |
| 20000 | 600 | 0.0111 ± 0.0017 | 0.0085 ± 0.0026 | 0.0082 ± 0.0027 |
| 20000 | 700 | 0.0111 ± 0.0018 | 0.0086 ± 0.0025 | 0.0082 ± 0.0027 |
| 20000 | 800 | 0.0111 ± 0.0018 | 0.0085 ± 0.0025 | 0.0082 ± 0.0027 |

#### Comparison with glmmTMB

**Table S11**: **Summary of differences between the estimated parameters from glmmTMB and the estimated parameters from FEMA**. Mean and standard deviation (across 84 experimental conditions) of the average (across 50 repeats) root mean squared error (RMSE) between ground truth and estimated coefficients from FEMA for differing total number of observations and minimum number of observations between pairs of visits

| $n_{\text{Obs}}$ | $\text{min}_{\text{numObs}}$ | RMSE beta: truth – FEMA | RMSE family: truth – FEMA | RMSE subject: truth – FEMA |
|---|---|---|---|---|
| 12000 | 500 | 0.0006 ± 0.0002 | 0.0053 ± 0.0021 | 0.0064 ± 0.0019 |
| 12000 | 600 | 0.0005 ± 0.0002 | 0.0050 ± 0.0021 | 0.0061 ± 0.0019 |
| 12000 | 700 | 0.0005 ± 0.0002 | 0.0050 ± 0.0021 | 0.0060 ± 0.0018 |
| 12000 | 800 | 0.0004 ± 0.0002 | 0.0048 ± 0.0020 | 0.0058 ± 0.0018 |
| 15000 | 500 | 0.0005 ± 0.0002 | 0.0043 ± 0.0018 | 0.0049 ± 0.0014 |
| 15000 | 600 | 0.0004 ± 0.0002 | 0.0042 ± 0.0017 | 0.0048 ± 0.0014 |
| 15000 | 700 | 0.0004 ± 0.0001 | 0.0041 ± 0.0018 | 0.0047 ± 0.0014 |
| 15000 | 800 | 0.0004 ± 0.0001 | 0.0040 ± 0.0017 | 0.0046 ± 0.0013 |
| 18000 | 500 | 0.0003 ± 0.0001 | 0.0030 ± 0.0013 | 0.0032 ± 0.0009 |
| 18000 | 600 | 0.0003 ± 0.0001 | 0.0030 ± 0.0013 | 0.0033 ± 0.0009 |
| 18000 | 700 | 0.0003 ± 0.0001 | 0.0030 ± 0.0013 | 0.0033 ± 0.0009 |
| 18000 | 800 | 0.0003 ± 0.0001 | 0.0030 ± 0.0013 | 0.0032 ± 0.0009 |
| 20000 | 500 | 0.0002 ± 0.0001 | 0.0022 ± 0.0009 | 0.0023 ± 0.0006 |
| 20000 | 600 | 0.0002 ± 0.0001 | 0.0022 ± 0.0010 | 0.0023 ± 0.0006 |
| 20000 | 700 | 0.0002 ± 0.0001 | 0.0022 ± 0.0010 | 0.0023 ± 0.0006 |
| 20000 | 800 | 0.0002 ± 0.0001 | 0.0022 ± 0.0010 | 0.0024 ± 0.0007 |

#### Simulation 5: false positive calibration across sample sizes

##### Sample size: 12,000

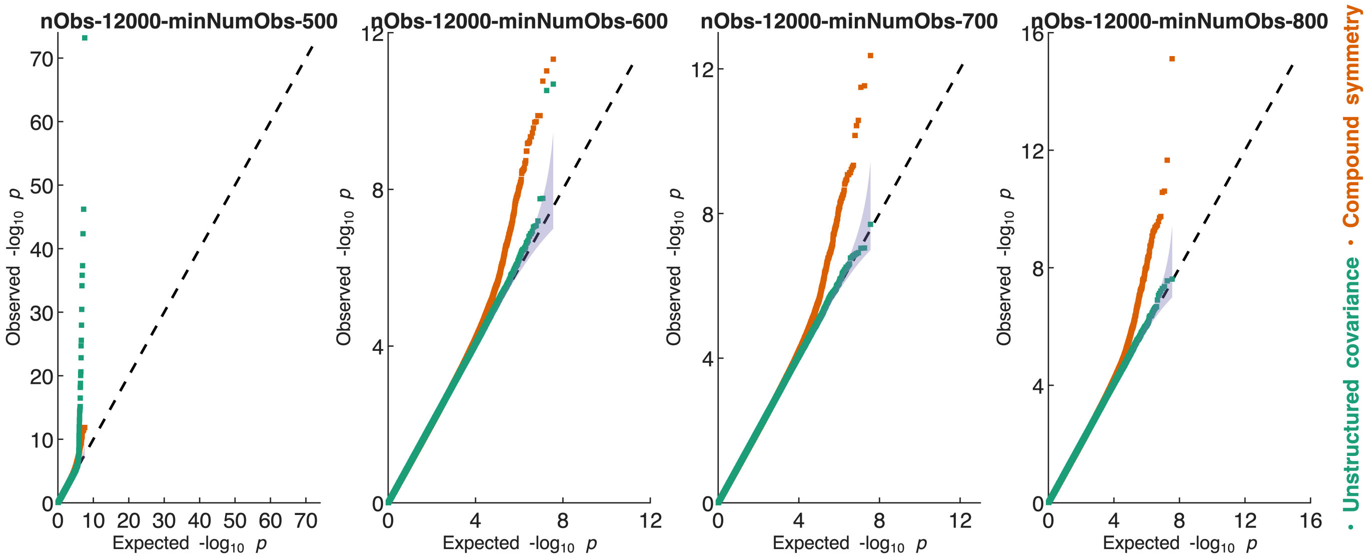


**Figure S26**: **Distribution of $-\log_{10}(p)$-values under the null for 12,000 observations and different minimum number of observations between visit pairs.** Each panel shows the distribution of $-\log_{10} p$-values across 1000 iterations of 36 simulation settings for unstructured covariance (green) and compound symmetry (orange); each iteration consisted of 100 $X$ variables and 10 outcome variables; the purple filled area indicates the 95% confidence interval based on inverse beta distribution.

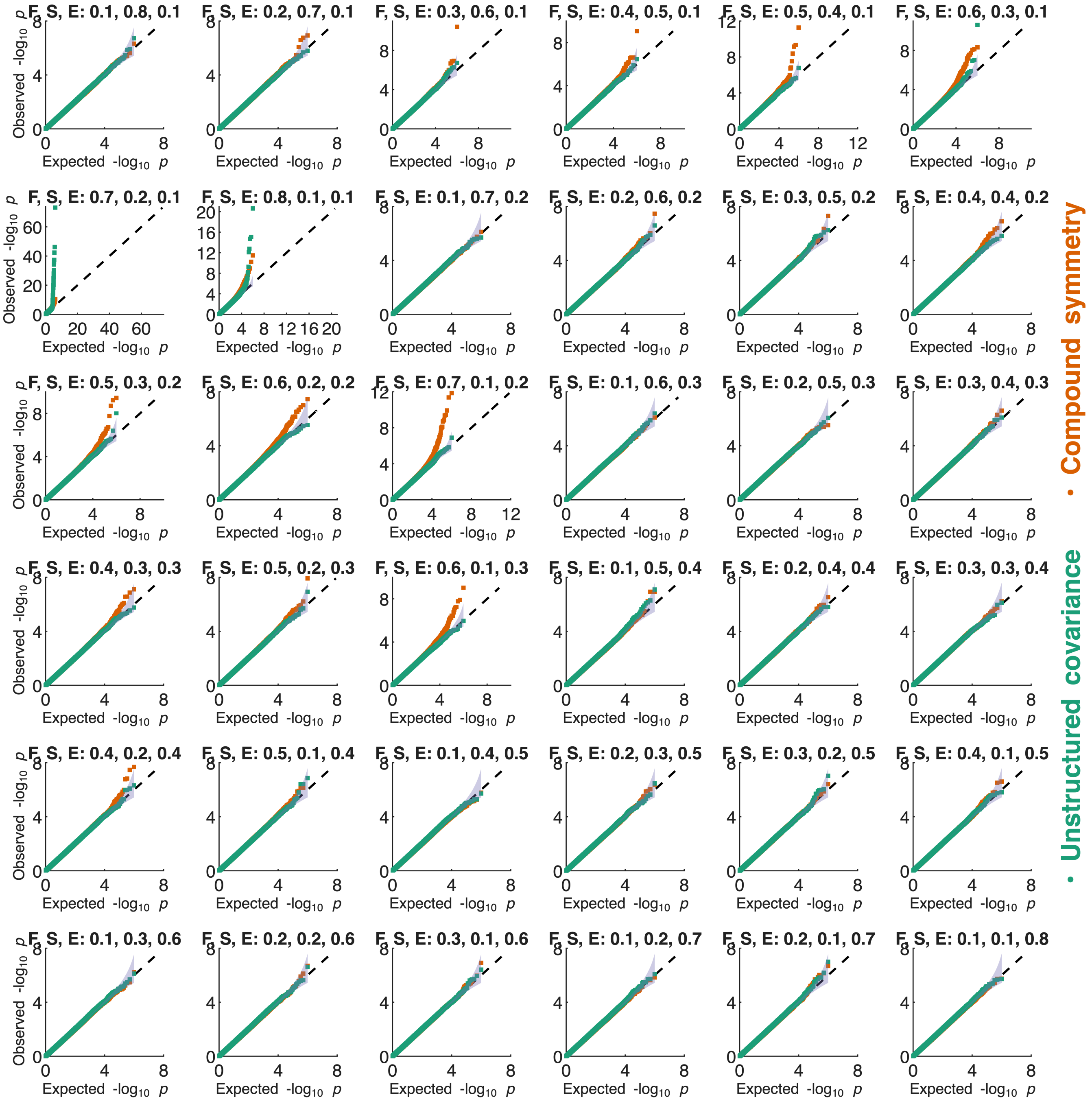


**Figure S27**: **Distribution of $-\log_{10}(p)$-values under the null for 12,000 observations and 500 minimum number of observations between visit pairs**. The simulation setting is indicated on the top of each Q-Q plot indicating the amounts of variances (in the phenotype) explained by family (F), subject (S), and noise (E); the *x*-axes indicate the expected $-\log_{10}(p)$ values under the null hypothesis while the *y*-axes show the observed $-\log_{10}(p)$ values across 1000 repeats, 100 $X$ variables, and 10 $y$ variables. The purple filled area indicates the 95% confidence interval based on inverse beta distribution.

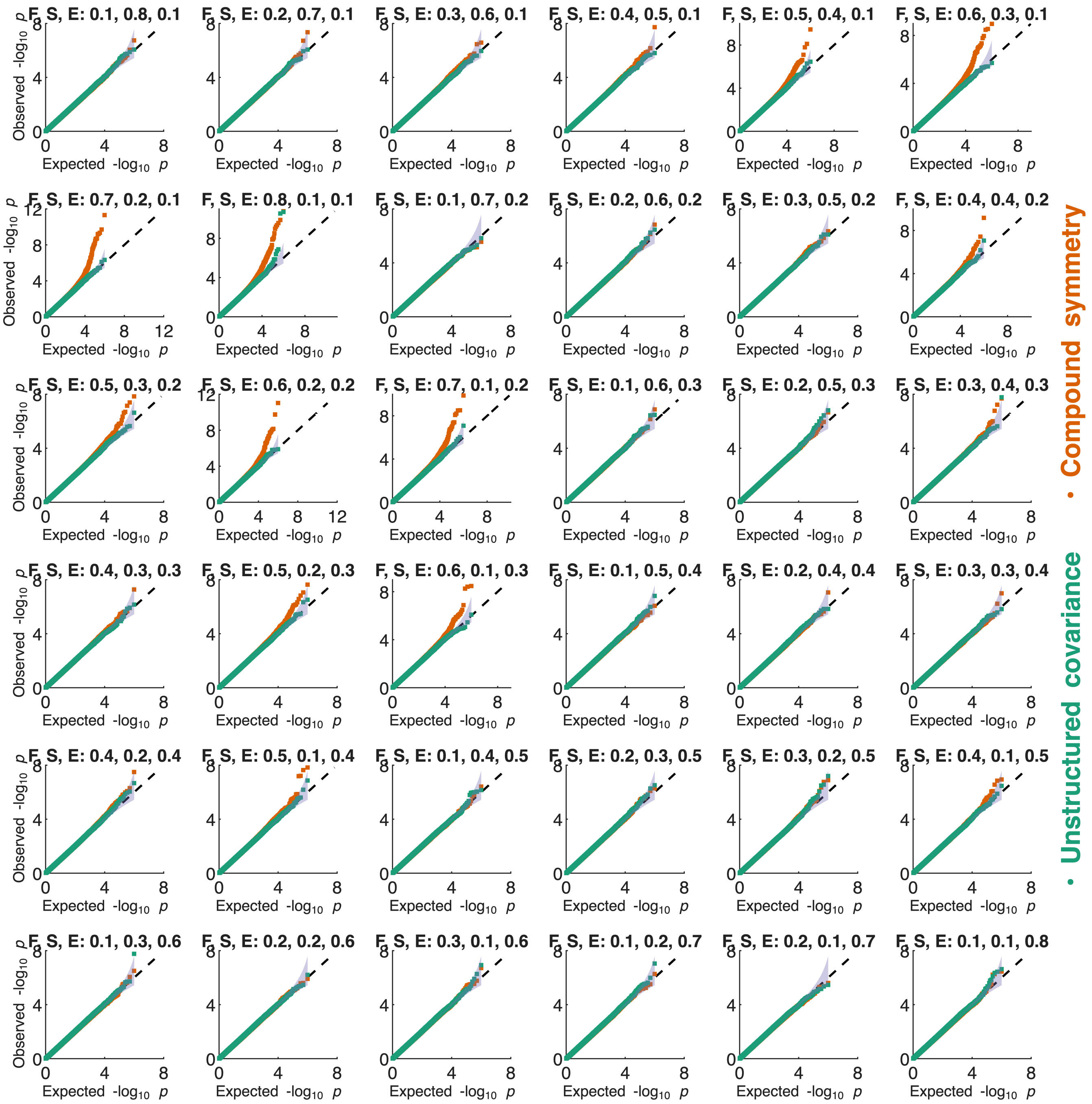


**Figure S28**: **Distribution of $-\log_{10}(p)$-values under the null for 12,000 observations and 600 minimum number of observations between visit pairs**. The simulation setting is indicated on the top of each Q-Q plot indicating the amounts of variances (in the phenotype) explained by family (F), subject (S), and noise (E); the $x$-axes indicate the expected $-\log_{10}(p)$ values under the null hypothesis while the $y$-axes show the observed $-\log_{10}(p)$ values across 1000 repeats, 100 $X$ variables, and 10 $y$ variables. The purple filled area indicates the 95% confidence interval based on inverse beta distribution.

#### Sample size 15,000

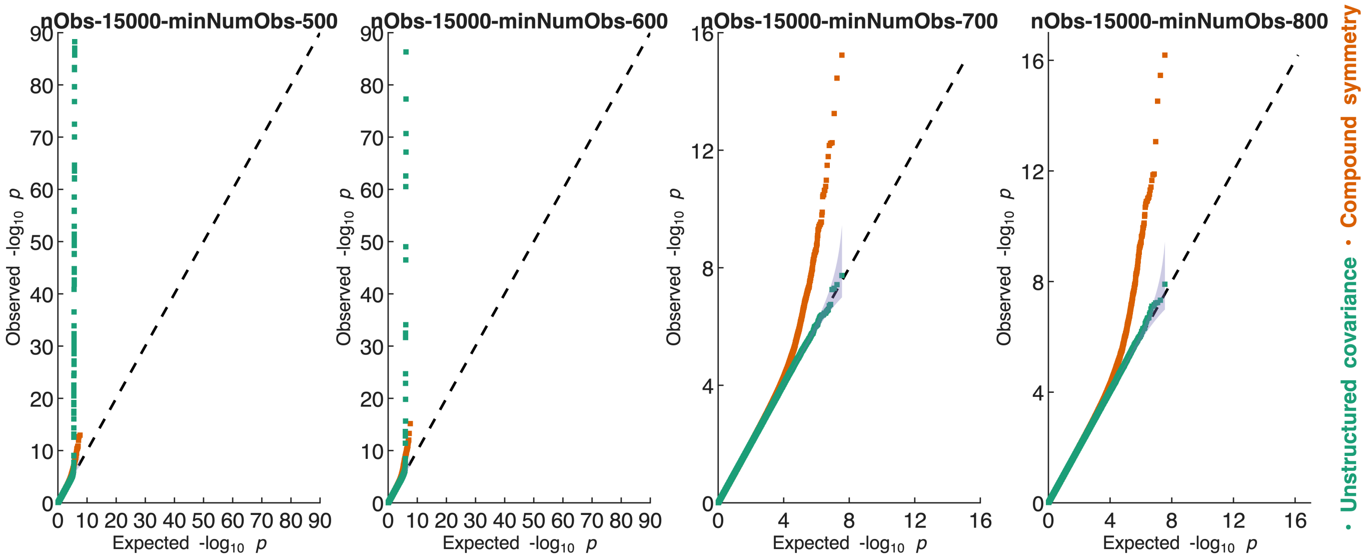


**Figure S29**: **Distribution of $-\log_{10}(p)$-values under the null for 15,000 observations and different minimum number of observations between visit pairs.** Each panel shows the distribution of $-\log_{10} p$-values across 1000 iterations of 36 simulation settings for unstructured covariance (green) and compound symmetry (orange); each iteration consisted of 100 $X$ variables and 10 outcome variables; the purple filled area indicates the 95% confidence interval based on inverse beta distribution. Note that the $y$-axis is truncated in the first two panels.

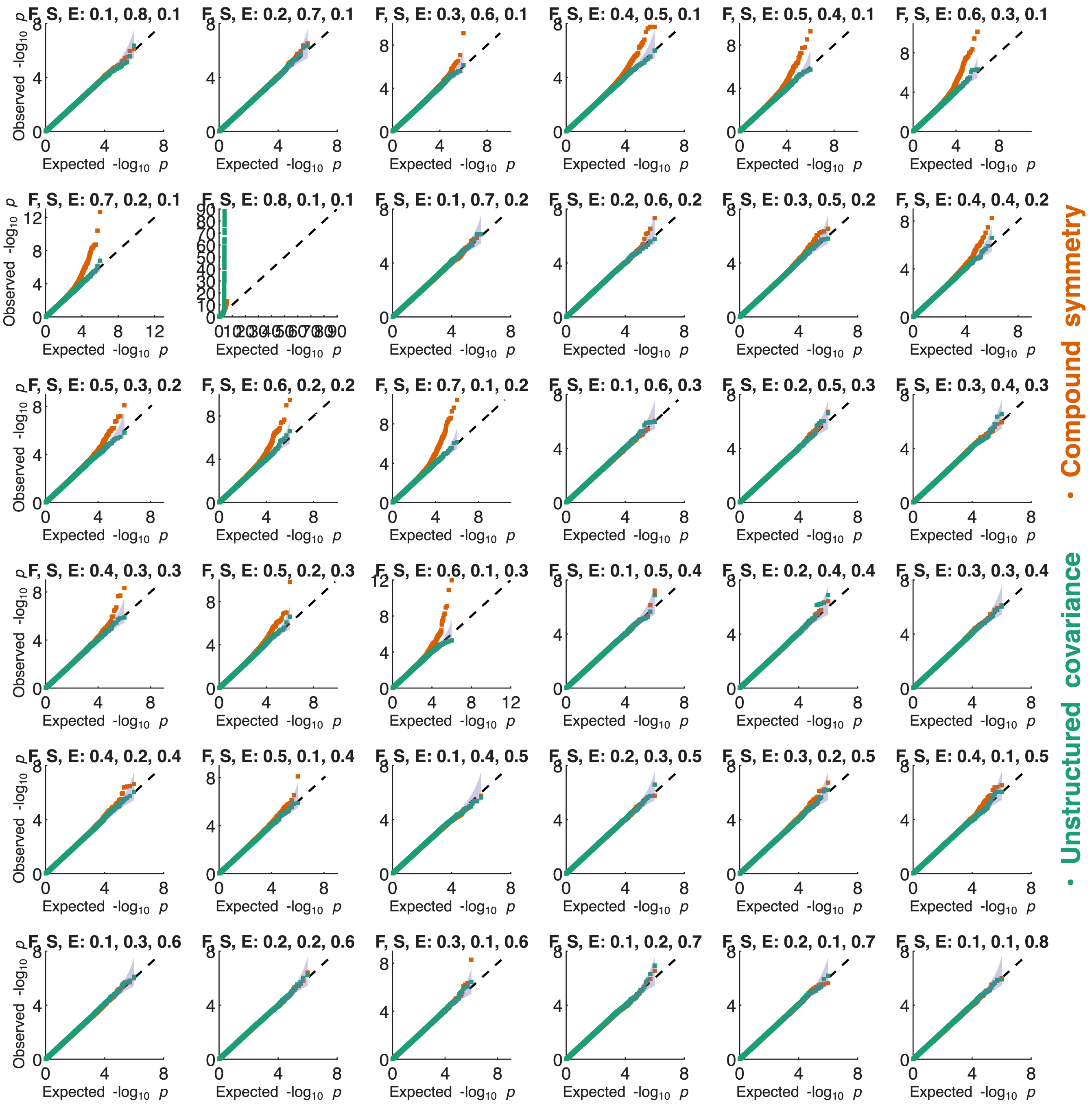


**Figure S30**: **Distribution of $-\log_{10}(p)$-values under the null for 15,000 observations and 500 minimum number of observations between visit pairs**. The simulation setting is indicated on the top of each Q-Q plot indicating the amounts of variances (in the phenotype) explained by family (F), subject (S), and noise (E); the $x$-axes indicate the expected $-\log_{10}(p)$ values under the null hypothesis while the $y$-axes show the observed $-\log_{10}(p)$ values across 1000 repeats, 100 $X$ variables, and 10 $y$ variables. The purple filled area indicates the 95% confidence interval based on inverse beta distribution. Note that the $y$-axis is truncated for the plot in second row, second column.

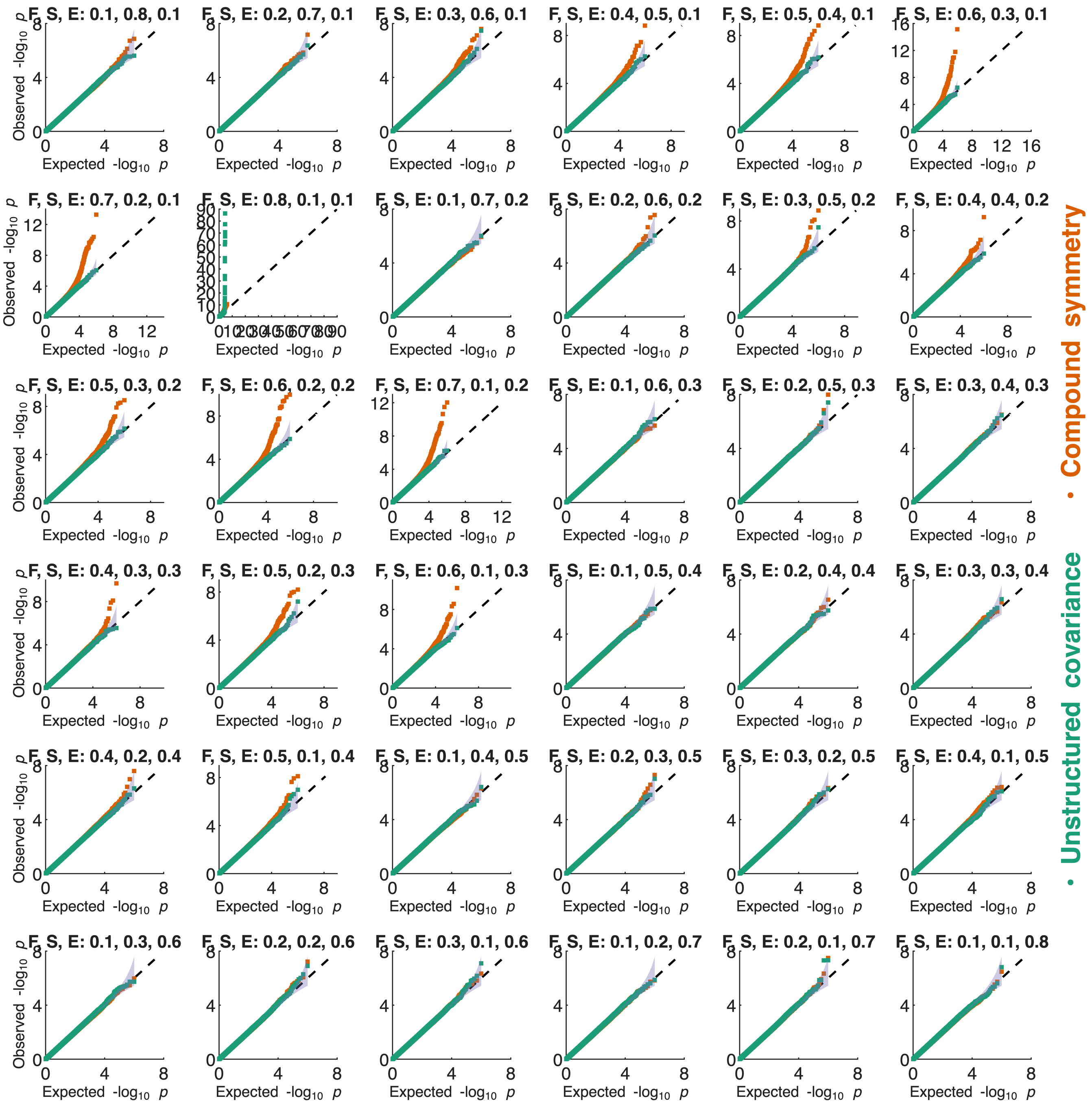


**Figure S31**: **Distribution of $-\log_{10}(p)$-values under the null for 15,000 observations and 600 minimum number of observations between visit pairs**. The simulation setting is indicated on the top of each Q-Q plot indicating the amounts of variances (in the phenotype) explained by family (F), subject (S), and noise (E); the $x$-axes indicate the expected $-\log_{10}(p)$ values under the null hypothesis while the $y$-axes show the observed $-\log_{10}(p)$ values across 1000 repeats, 100 $X$ variables, and 10 $y$ variables. The purple filled area indicates the 95% confidence interval based on inverse beta distribution. Note that the $y$-axis is truncated for the plot in second row, second column.

#### Sample size 18,000

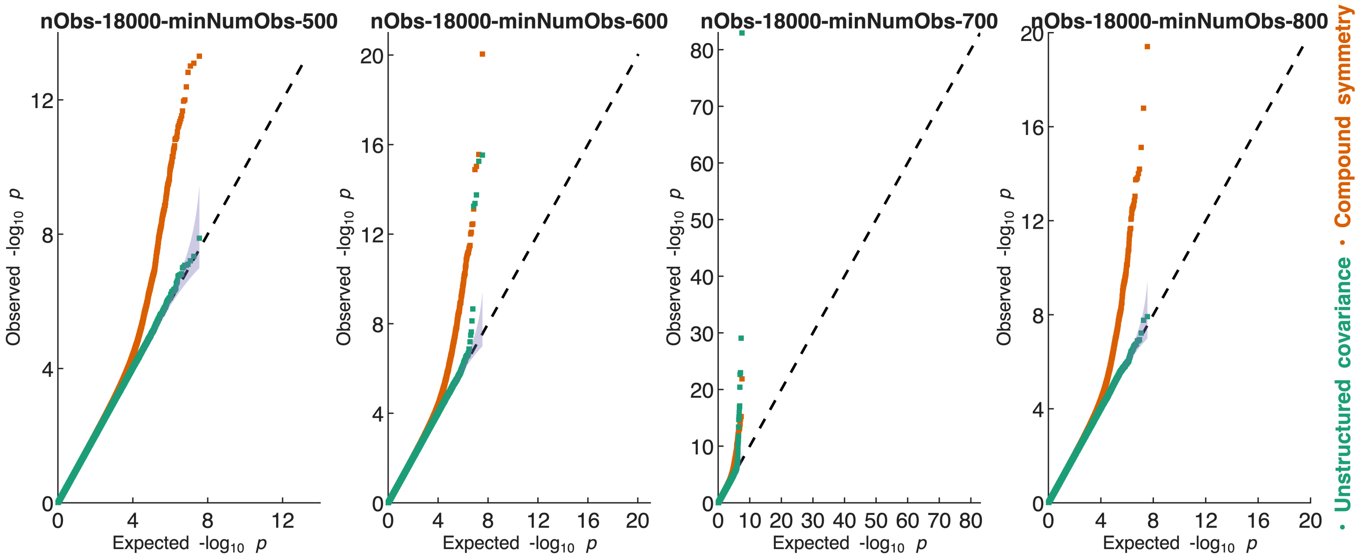


**Figure S32**: **Distribution of $-\log_{10}(p)$-values under the null for 18,000 observations and different minimum number of observations between visit pairs.** Each panel shows the distribution of $-\log_{10} p$-values across 1000 iterations of 36 simulation settings for unstructured covariance (green) and compound symmetry (orange); each iteration consisted of 100 $X$ variables and 10 outcome variables; the purple filled area indicates the 95% confidence interval based on inverse beta distribution.

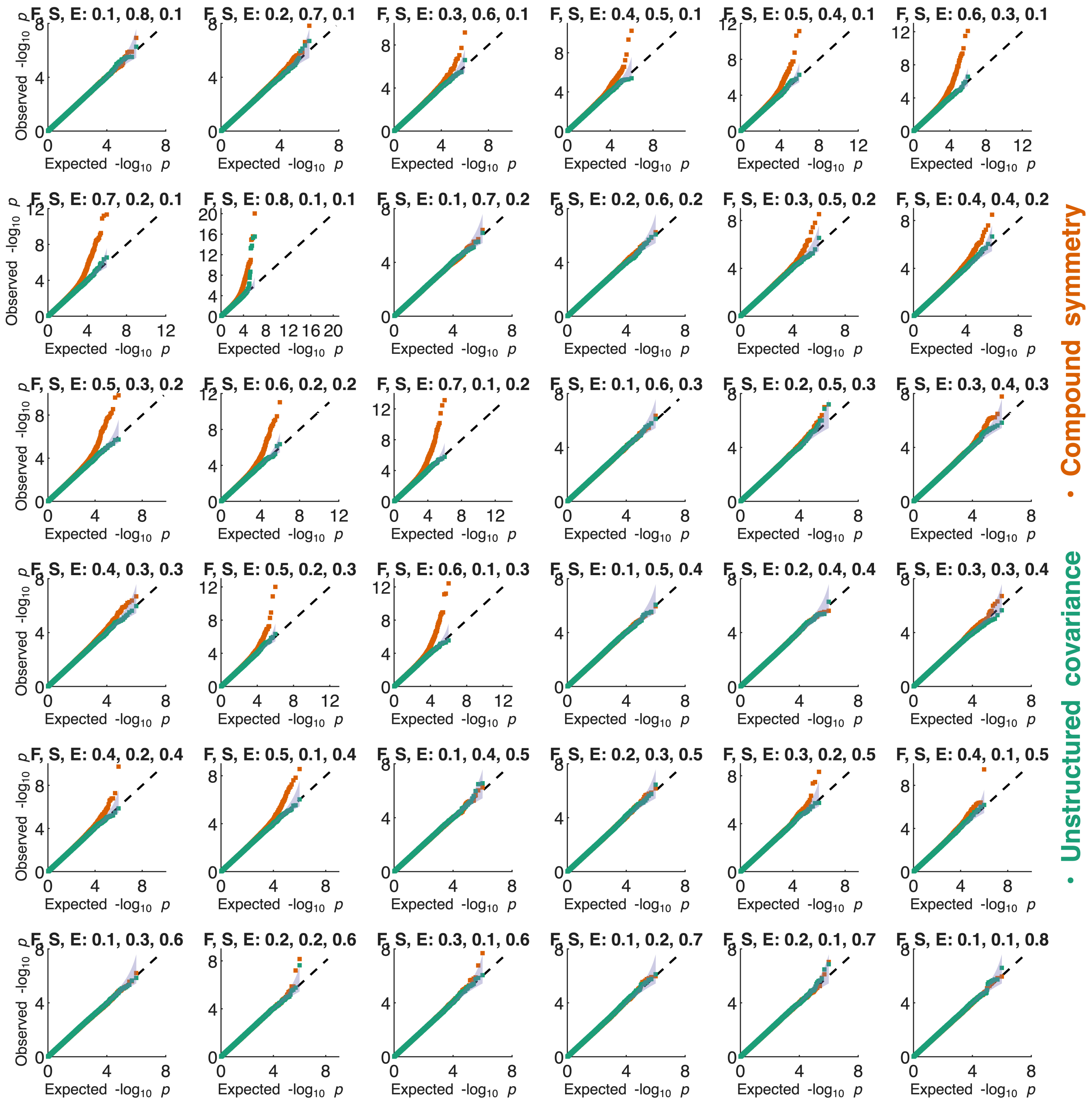


**Figure S33**: **Distribution of $-\log_{10}(p)$-values under the null for 18,000 observations and 600 minimum number of observations between visit pairs**. The simulation setting is indicated on the top of each Q-Q plot indicating the amounts of variances (in the phenotype) explained by family (F), subject (S), and noise (E); the $x$-axes indicate the expected $-\log_{10}(p)$ values under the null hypothesis while the $y$-axes show the observed $-\log_{10}(p)$ values across 1000 repeats, 100 $X$ variables, and 10 $y$ variables. The purple filled area indicates the 95% confidence interval based on inverse beta distribution.

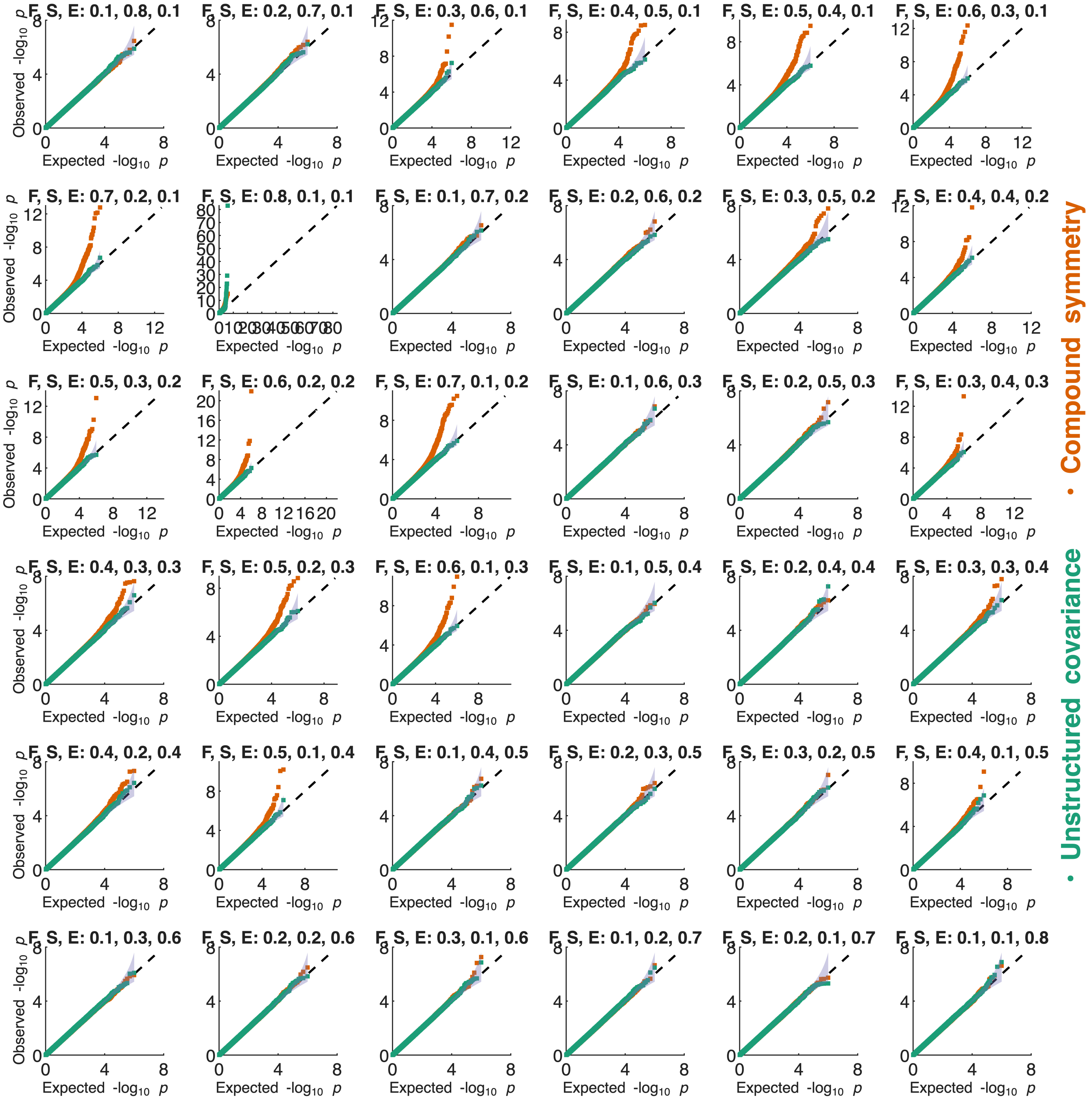


**Figure S34**: **Distribution of $-\log_{10}(p)$-values under the null for 18,000 observations and 700 minimum number of observations between visit pairs**. The simulation setting is indicated on the top of each Q-Q plot indicating the amounts of variances (in the phenotype) explained by family (F), subject (S), and noise (E); the $x$-axes indicate the expected $-\log_{10}(p)$ values under the null hypothesis while the $y$-axes show the observed $-\log_{10}(p)$ values across 1000 repeats, 100 $X$ variables, and 10 $y$ variables. The purple filled area indicates the 95% confidence interval based on inverse beta distribution.

#### Sample size 20,000

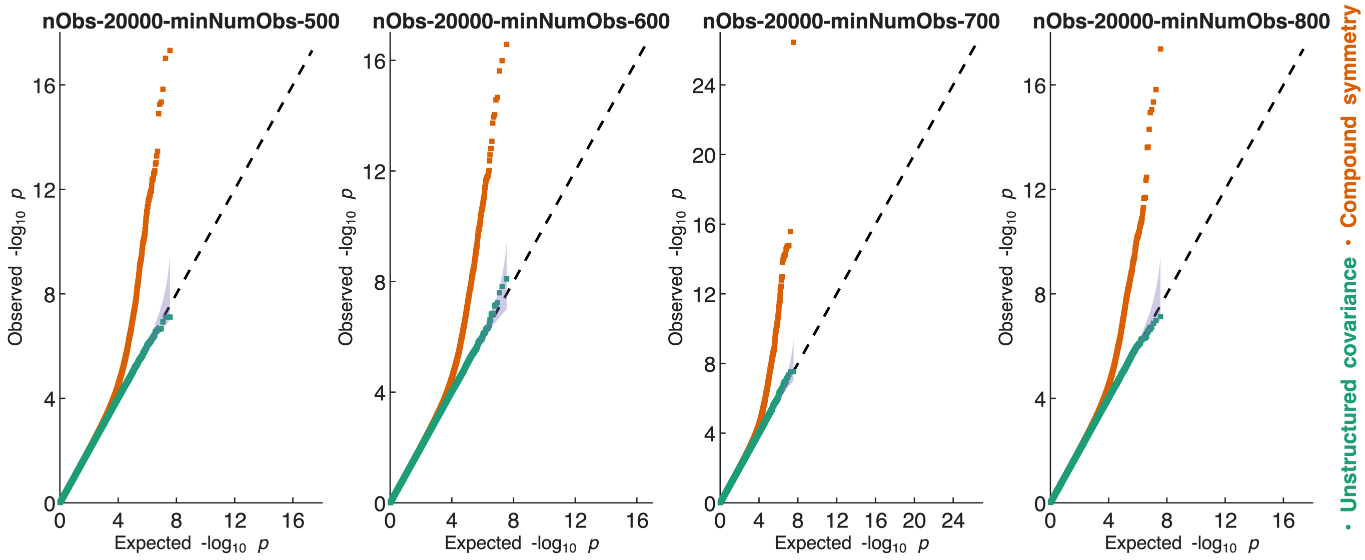


**Figure S35**: **Distribution of $-\log_{10}(p)$-values under the null for 20,000 observations and different minimum number of observations between visit pairs.** Each panel shows the distribution of $-\log_{10} p$-values across 1000 iterations of 36 simulation settings for unstructured covariance (green) and compound symmetry (orange); each iteration consisted of 100 $X$ variables and 10 outcome variables; the purple filled area indicates the 95% confidence interval based on inverse beta distribution.

#### Examples of scenarios where the unstructured covariance resulted in inflated *p*-values

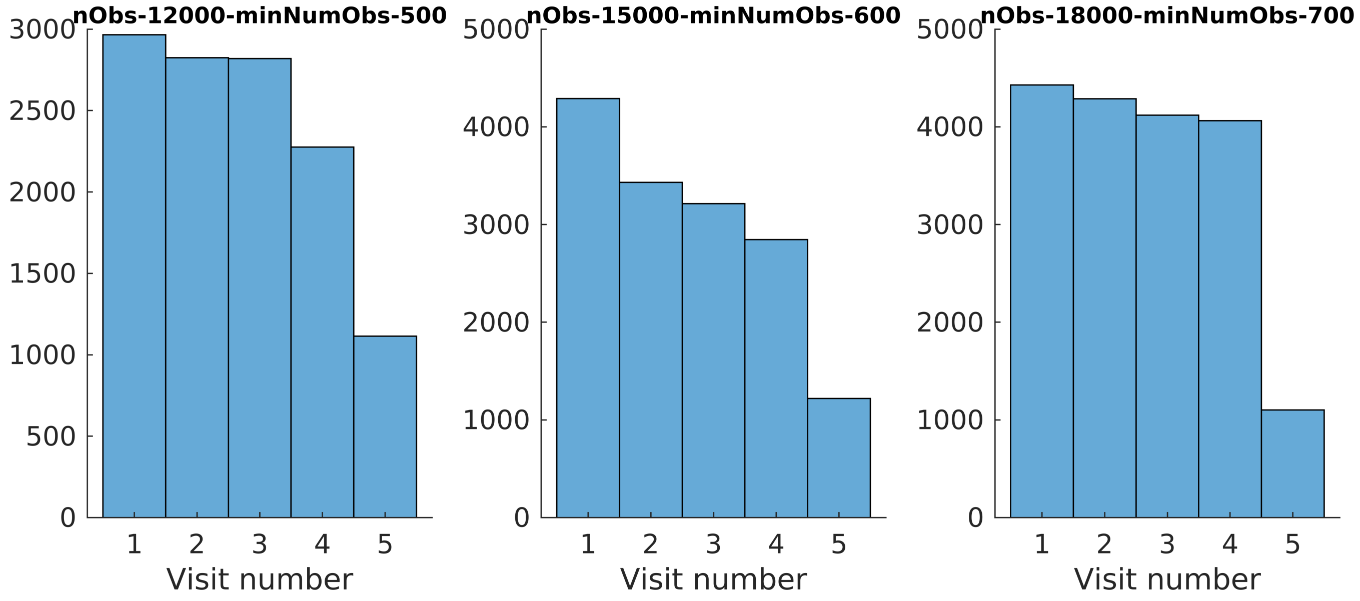


**Figure S36**: **Three examples where the $p$-values for unstructured covariance were not well calibrated**. In these three examples, by chance, the simulation setting created a situation where the last visit had limited sample size; therefore, when estimating the covariance for the last pair of visits, the overlapping sample size would be close to (or equal to) $\min_{\text{numObs}}$, likely leading to unstable covariance estimation, leading to inflated $p$-values.

### Phenotype

#### Age

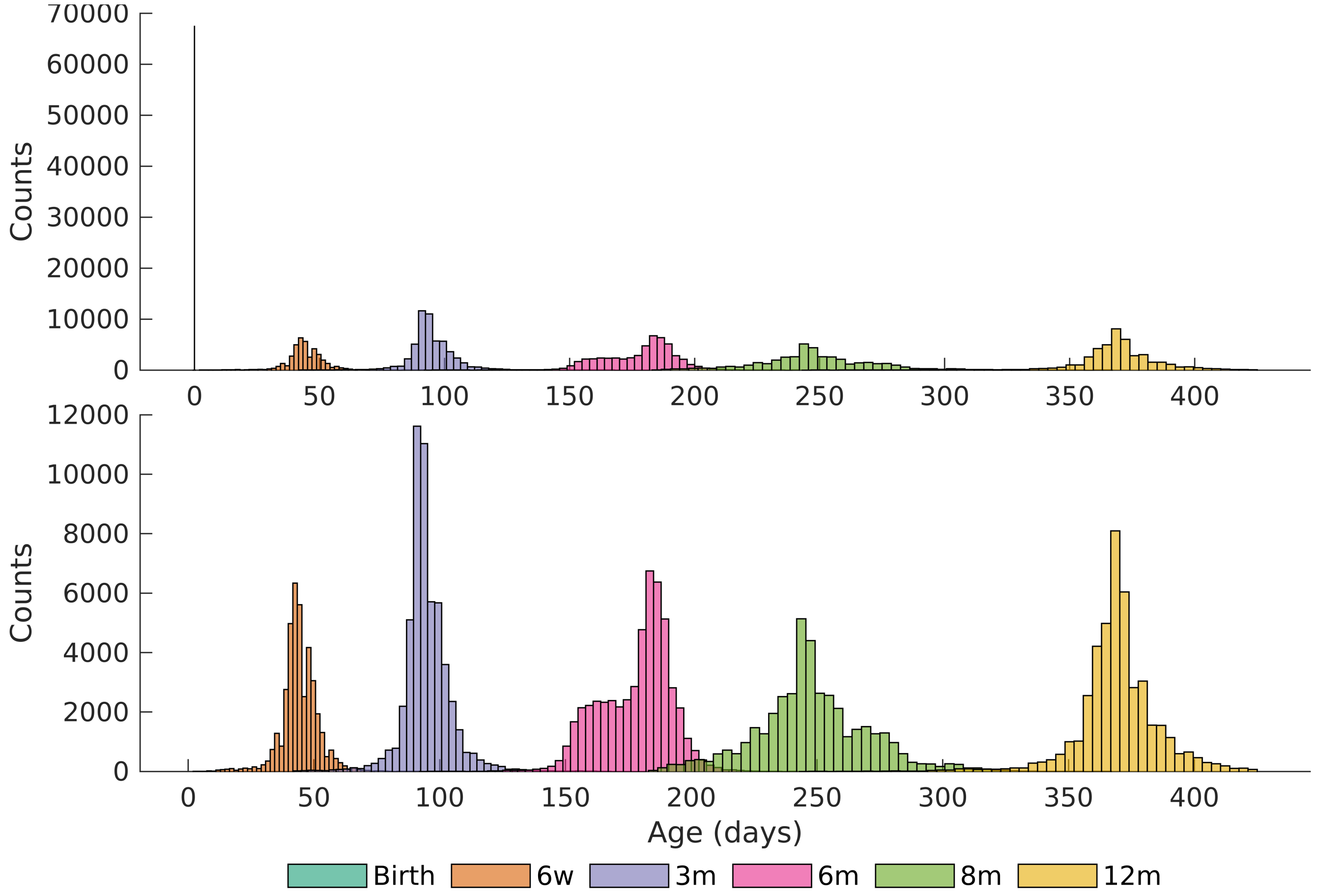


**Figure S37: Histogram of age for each visit.** The top panel shows the histogram of age across the six time points while the lower panel shows a zoomed-in version of the same (excluding the birth time point).

**Length**

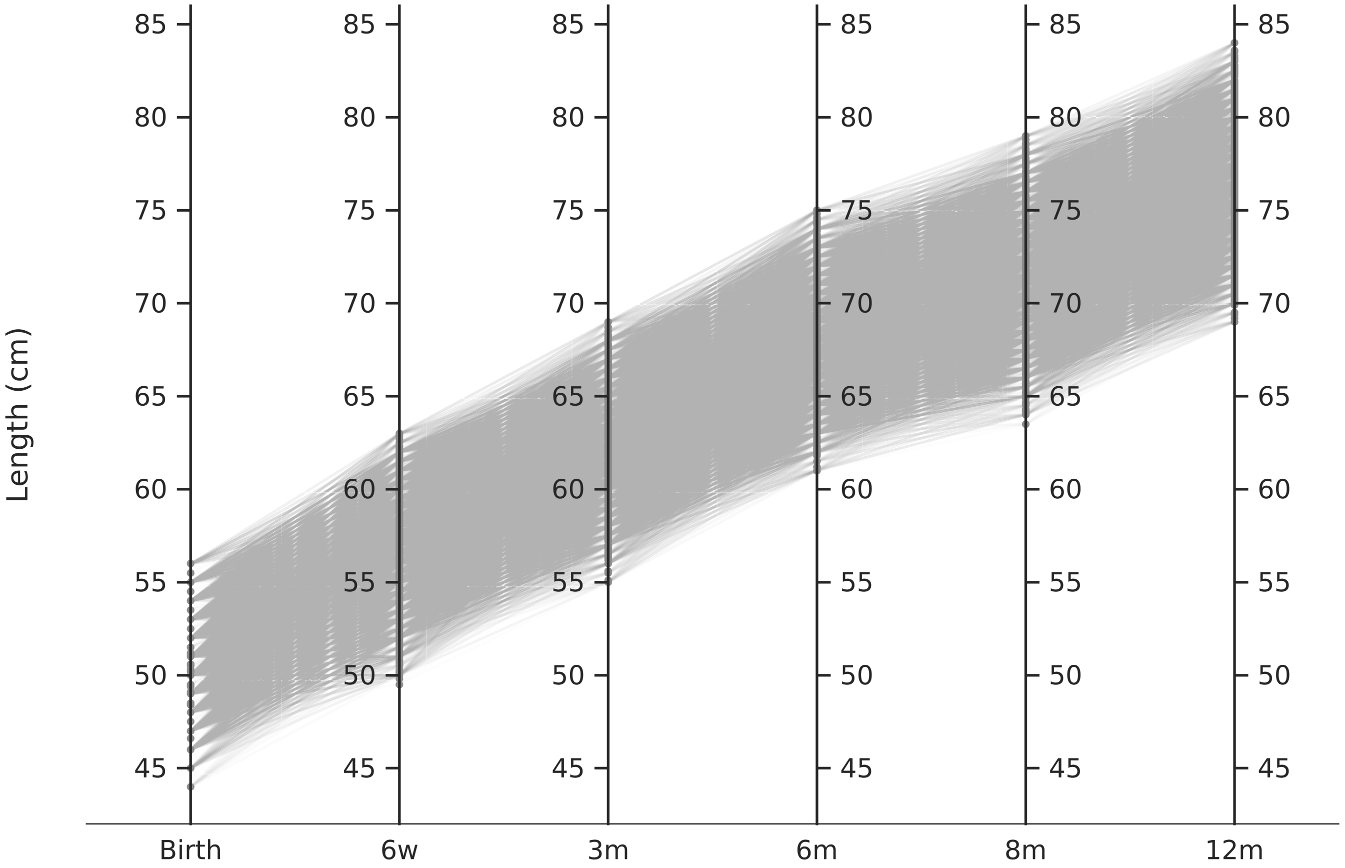


**Figure S38: Trajectories of length in the MoBa sample.** Each line shows the trajectory of length (in centimeters) from birth to the first year of life for each MoBa participant (*n* = 68,273 infants; 299,447 observations).

**Weight**

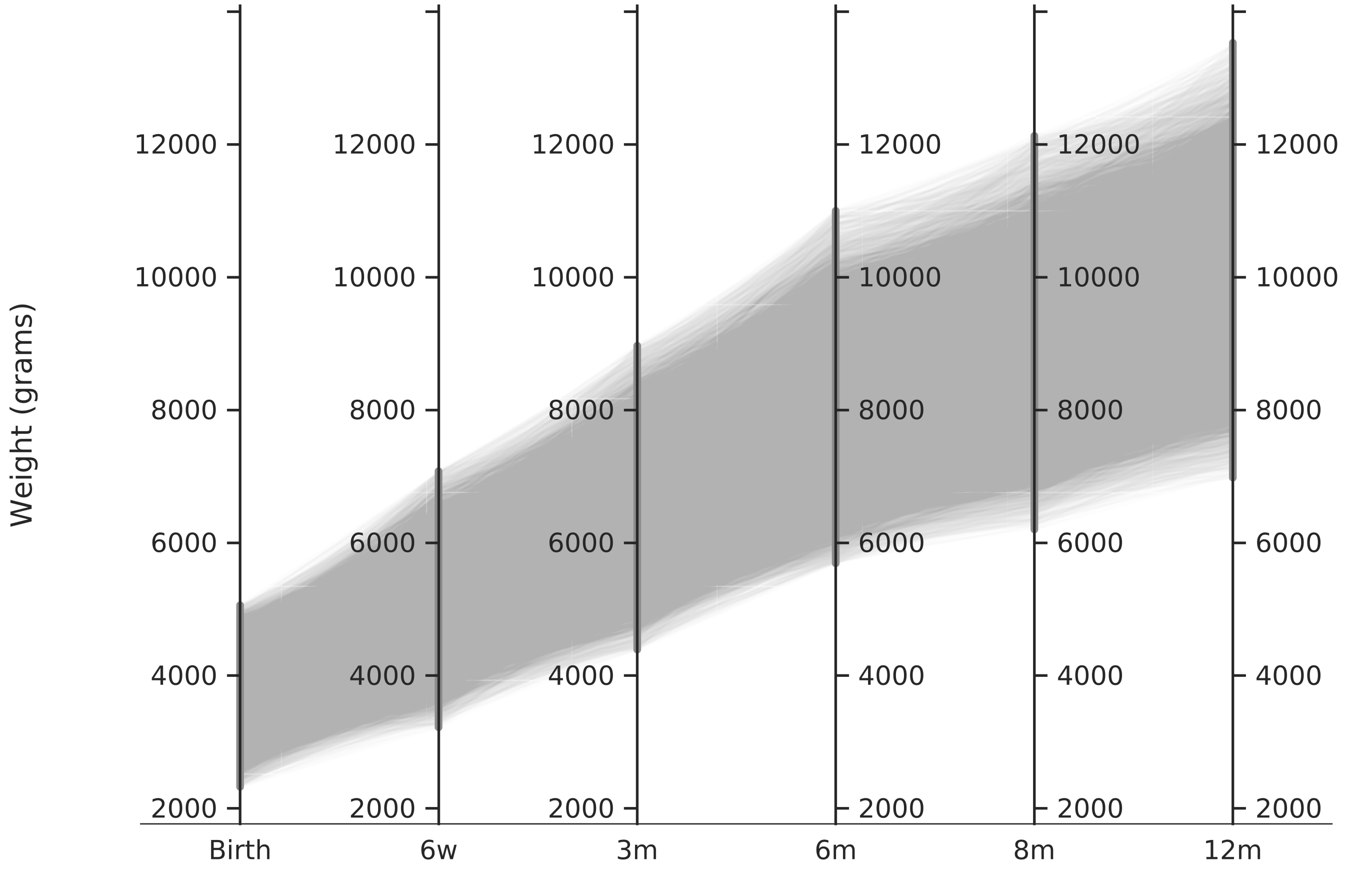


**Figure S39: Trajectories of weight in the MoBa sample.** Each line shows the trajectory of weight (in grams) from birth to the first year of life for each MoBa participant (*n* = 68,273 infants; 299,447 observations).

**BMI**

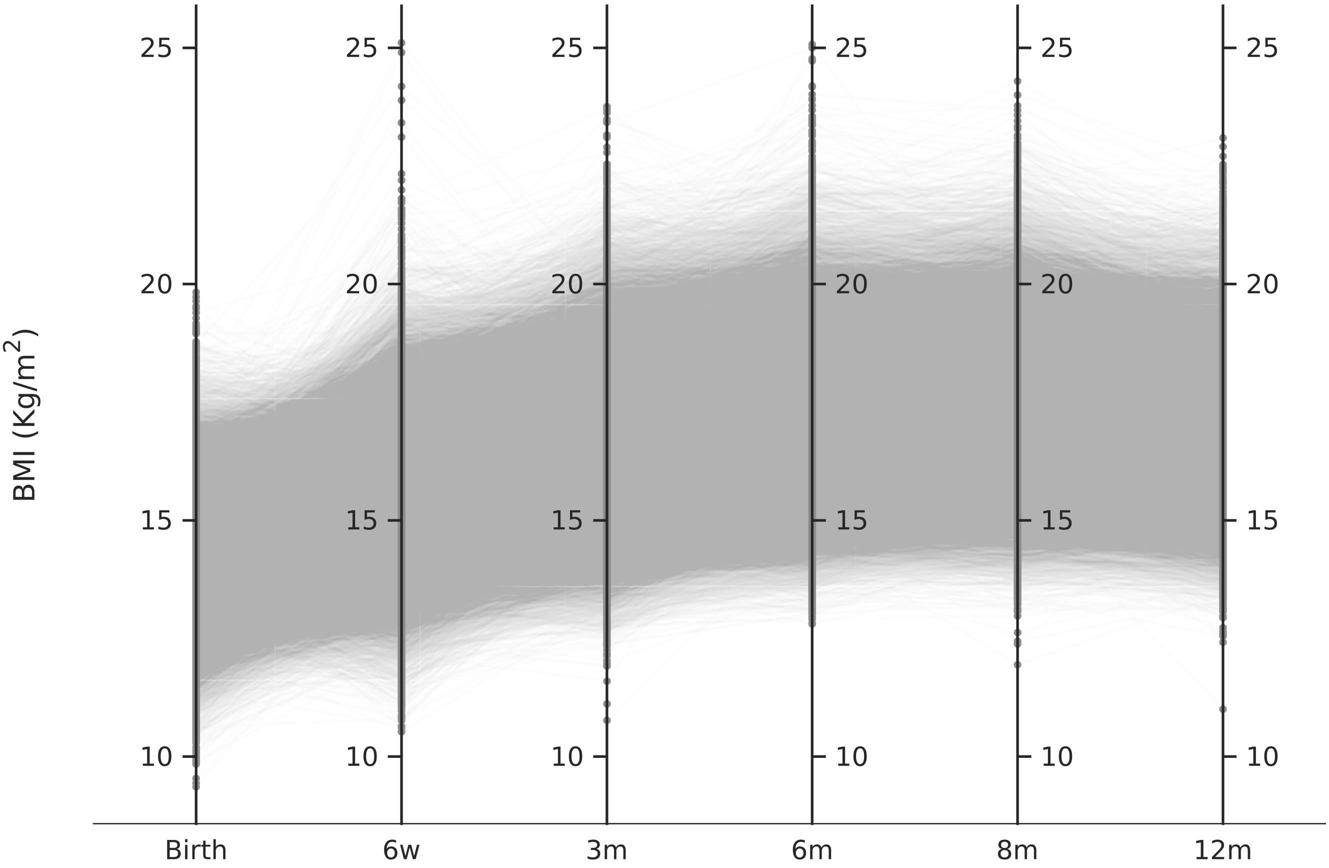


**Figure S40: Trajectories of BMI in the MoBa sample.** Each line shows the trajectory of BMI (kilograms per square meters) from birth to the first year of life for each MoBa participant (*n* = 68,273 infants; 299,447 observations).

**Knot placement for creating spline basis functions**

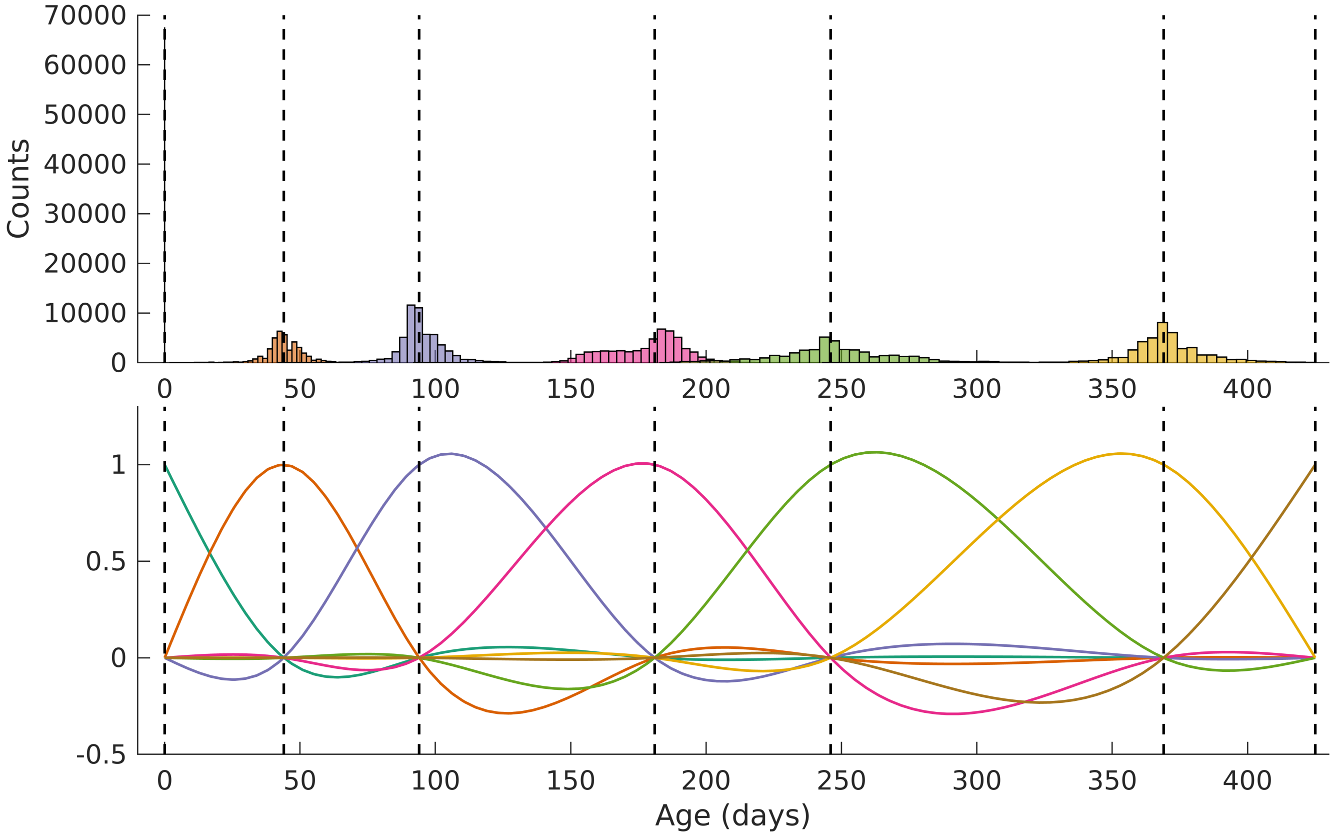


**Figure S41: Placement of knots for creating spline basis functions.** The top panel shows the histogram of age (in days) while the lower panel shows the created spline basis functions of age; the dashed vertical black lines indicate the placement of knots.

**Basis functions used for analysis**

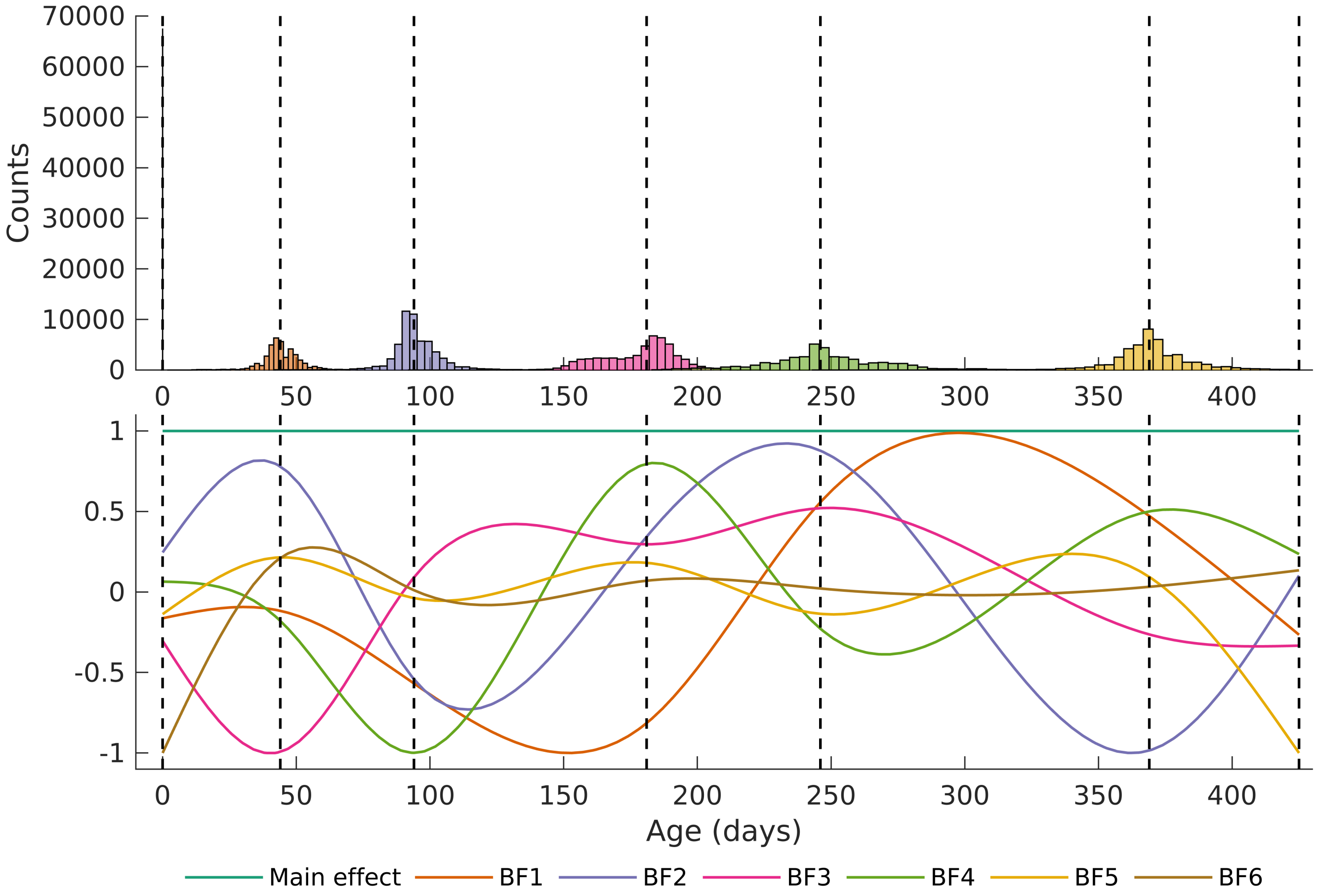


**Figure S42: Modified basis functions that were used for analysis.** The top panel shows the histogram of age (in days) while the lower panel shows the created spline basis functions of age, followed by a singular value decomposition, and addition of the constant term; the dashed vertical black lines indicate the placement of knots.

### Q-Q plots

#### Length

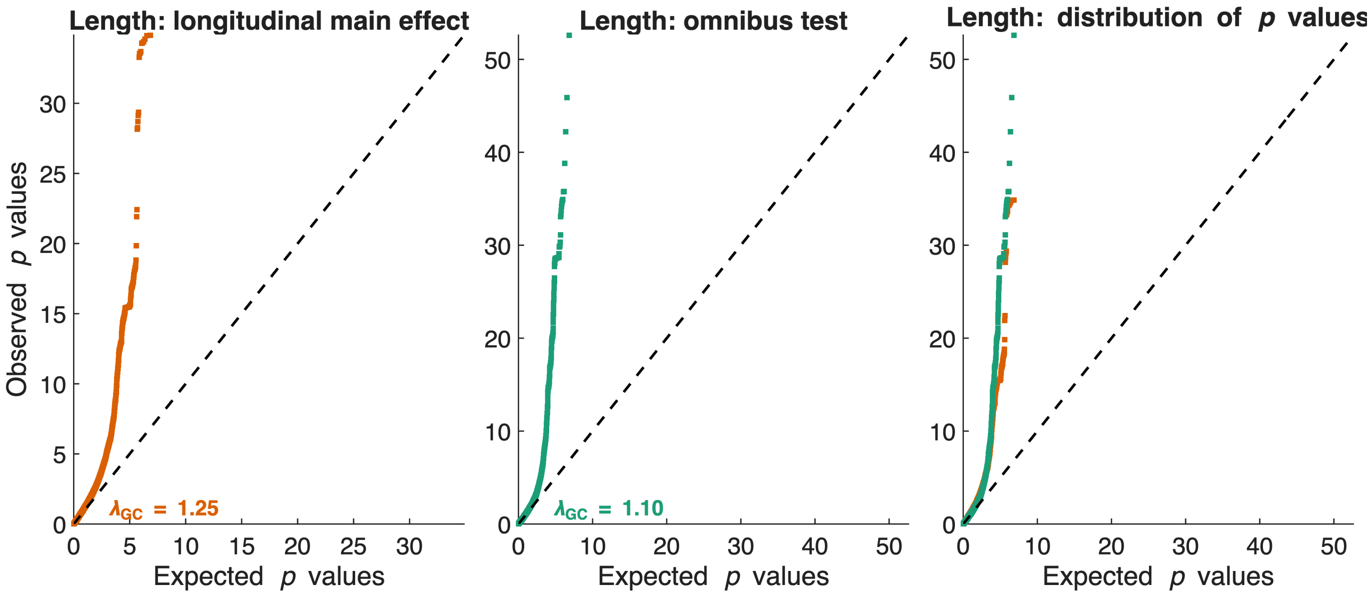


**Figure S43**: Q-Q plots for length from standard longitudinal GWAS without modeling smooth effects of age or interaction of SNPs with smooth functions of age (**left**); from omnibus test across the main effect of age and the interaction of SNPs with the smooth functions of age (**middle**); and both the sets of *p*-values overlaid (**right**).

#### Weight

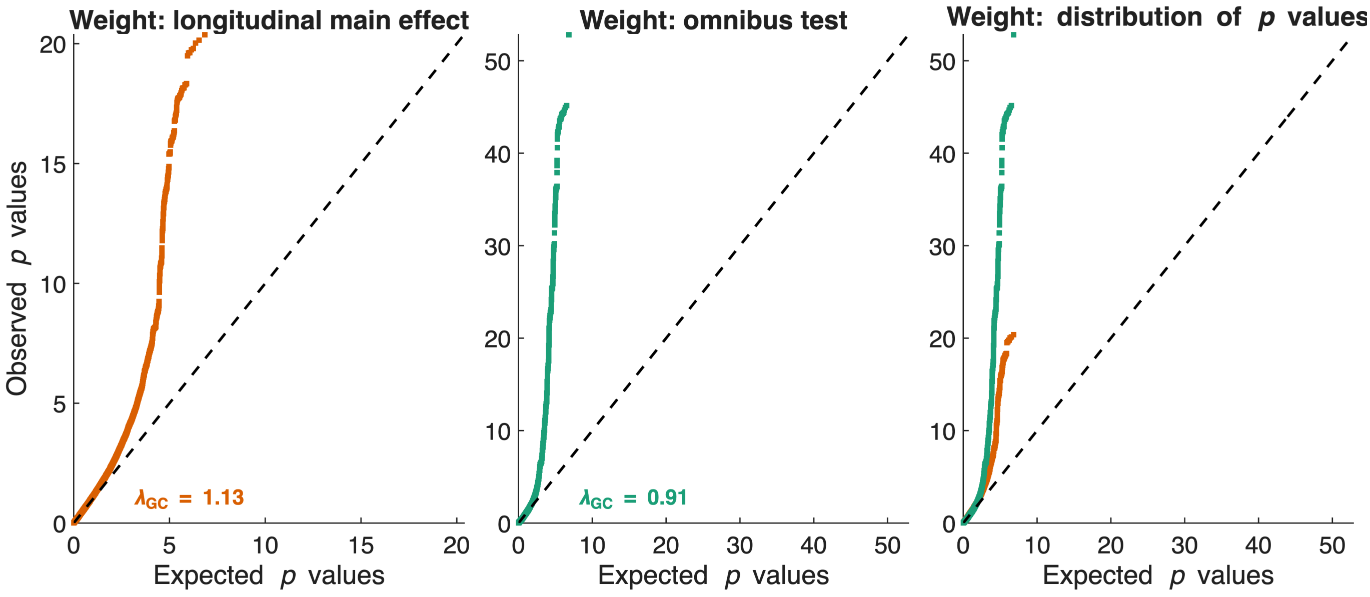


**Figure S44**: Q-Q plots for weight from standard longitudinal GWAS without modeling smooth effects of age or interaction of SNPs with smooth functions of age (**left**); from omnibus test across the main effect of age and the interaction of SNPs with the smooth functions of age (**middle**); and both the sets of *p*-values overlaid (**right**).

**BMI**

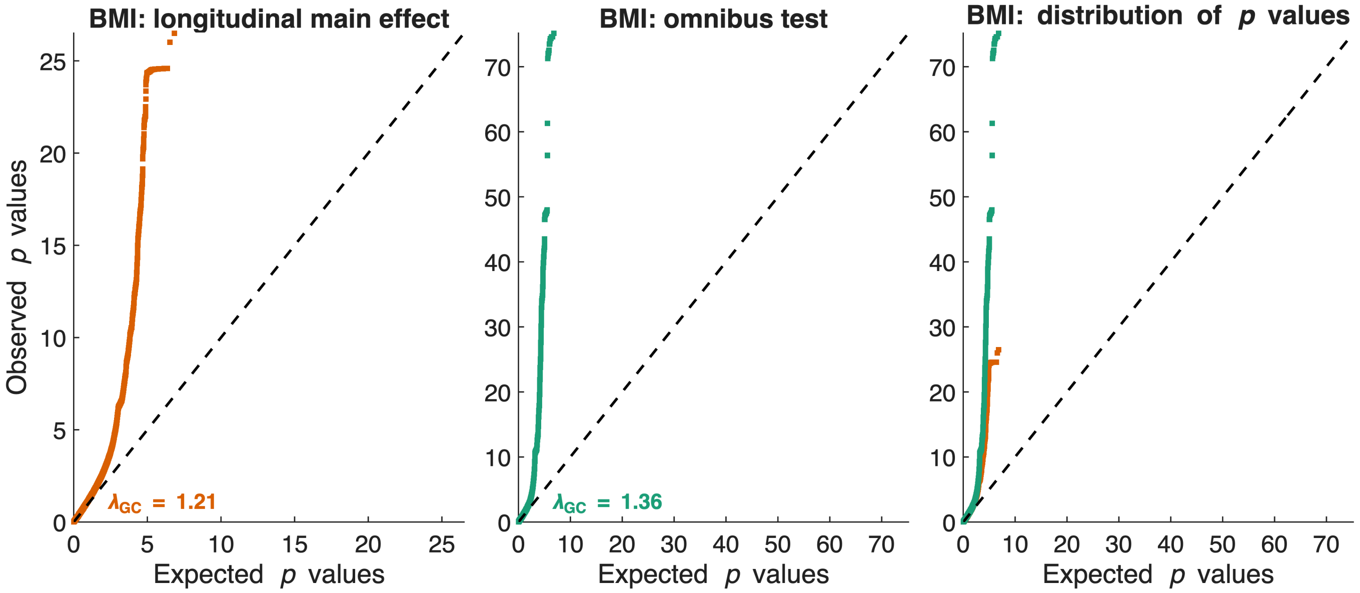


**Figure S45**: Q-Q plots for BMI from standard longitudinal GWAS without modeling smooth effects of age or interaction of SNPs with smooth functions of age (**left**); from omnibus test across the main effect of age and the interaction of SNPs with the smooth functions of age (**middle**); and both the sets of *p*-values overlaid (**right**).

#### Comparison between two-stage regression and full regression

When fitting GWAS-like models, we assume that the effect of each SNP is small and would not have an impact on the estimation of the random effects covariance components. To evaluate the impact of this assumption, for a selected subset of SNPs, we performed a comparison of parameter estimation from fitting a full model vs. when fitting a two-stage model. Concretely, from the time-varying GWAS analyses of length, weight, and BMI, we extracted top ten SNPs per chromosome, per phenotype. This resulted in a subset of 1,110 SNPs. For each of these SNPs $g$, we fit the full regression model (i.e., 1,110 separate models):

$$\text{Phenotype} \sim 1 + g + g \odot s(\text{age}) + s(\text{age}) + \text{sex} + \text{PC}_{1-20} + \text{Batch} + \text{us}(1|\text{Family}) + \text{us}(1|\text{GRM}) + \text{us}(1|\text{Subject}) \quad \text{(S1)}$$

Then, we compared the estimated model parameters from the full model against the estimated parameters when performing two-stage regression described in the main manuscript. The results indicate that the two approaches yield comparable estimates.

##### Comparison of beta coefficients

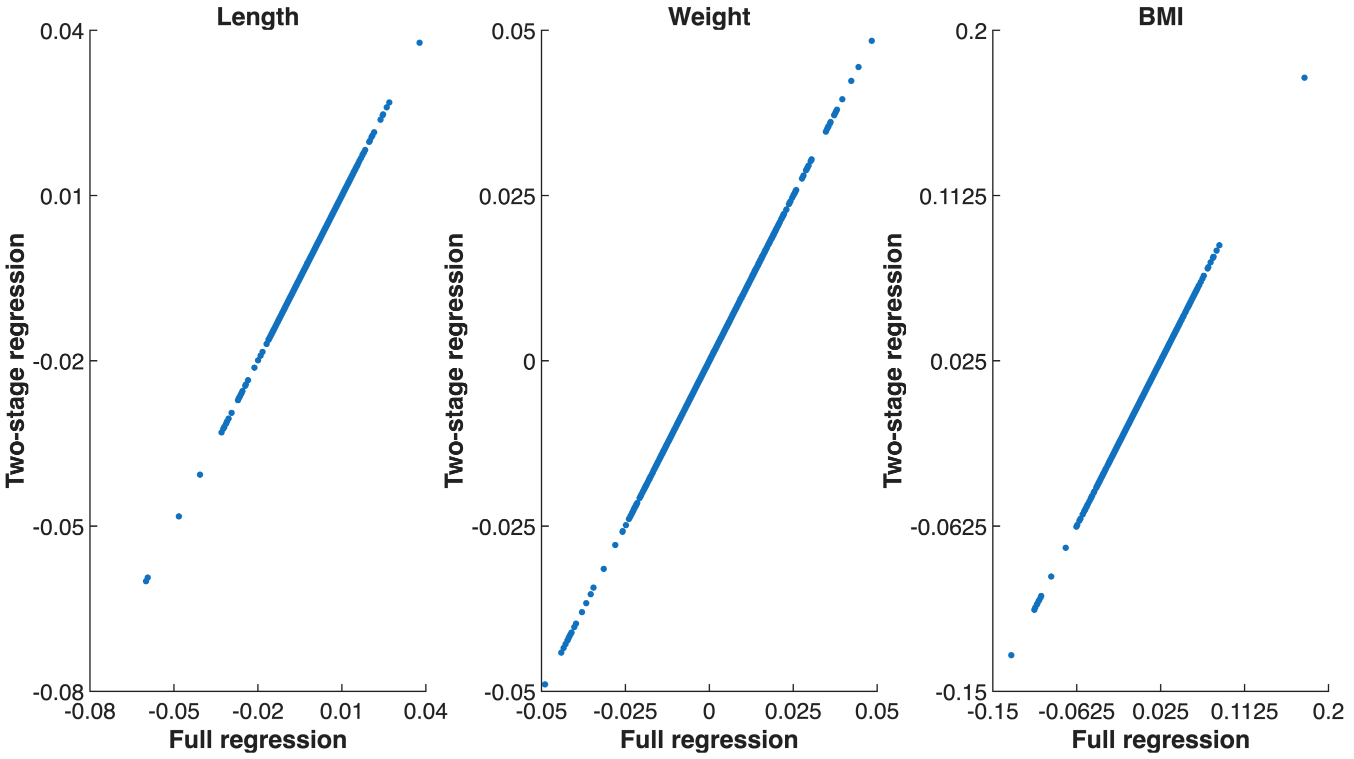


**Figure S46:** Comparison of beta coefficients (main effect and the interaction effects of SNPs with smooth basis functions of age) between full regression (where all terms were estimated together) vs. two-stage regression (where the stage-1 regression and the estimation of random effects covariance terms did not include genetics).

#### Comparison of standard error

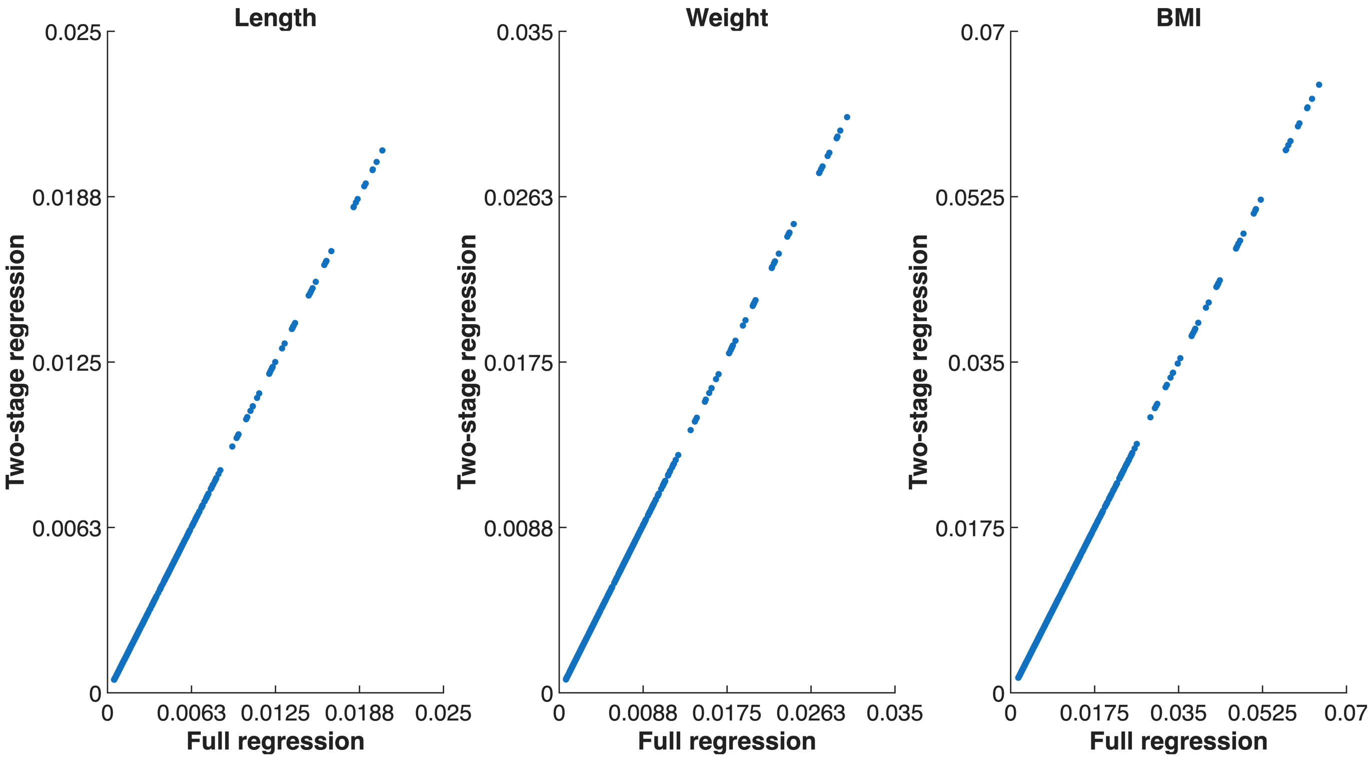


**Figure S47:** Comparison of standard errors (main effect and the interaction effects of SNPs with smooth basis functions of age) between full regression (where all terms were estimated together) vs. two-stage regression (where the stage-1 regression and the estimation of random effects covariance terms did not include genetics).

#### Comparison of *T* statistics

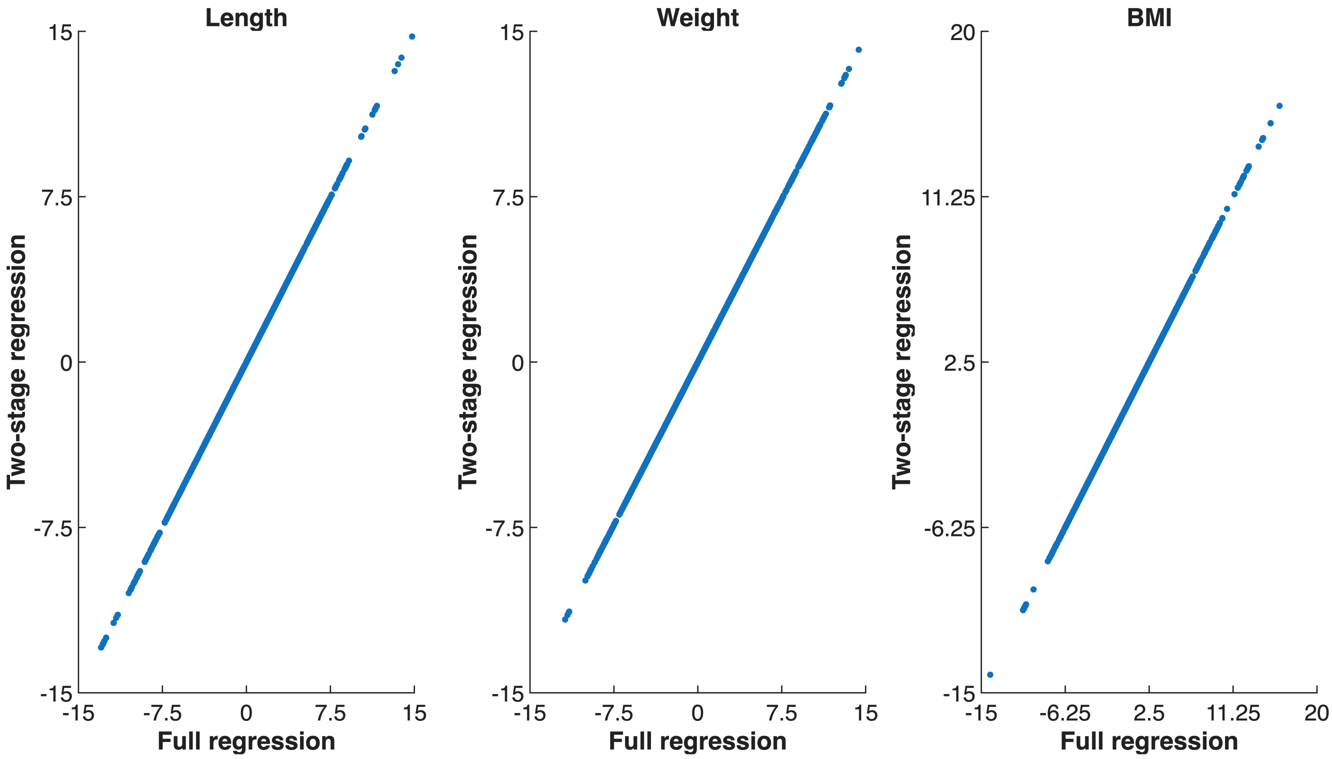


**Figure S48:** Comparison of *T* statistics (main effect and the interaction effects of SNPs with smooth basis functions of age) between full regression (where all terms were estimated together) vs. two-stage regression (where the stage-1 regression and the estimation of random effects covariance terms did not include genetics).

**Comparison of $-\log_{10}(p)$ values**

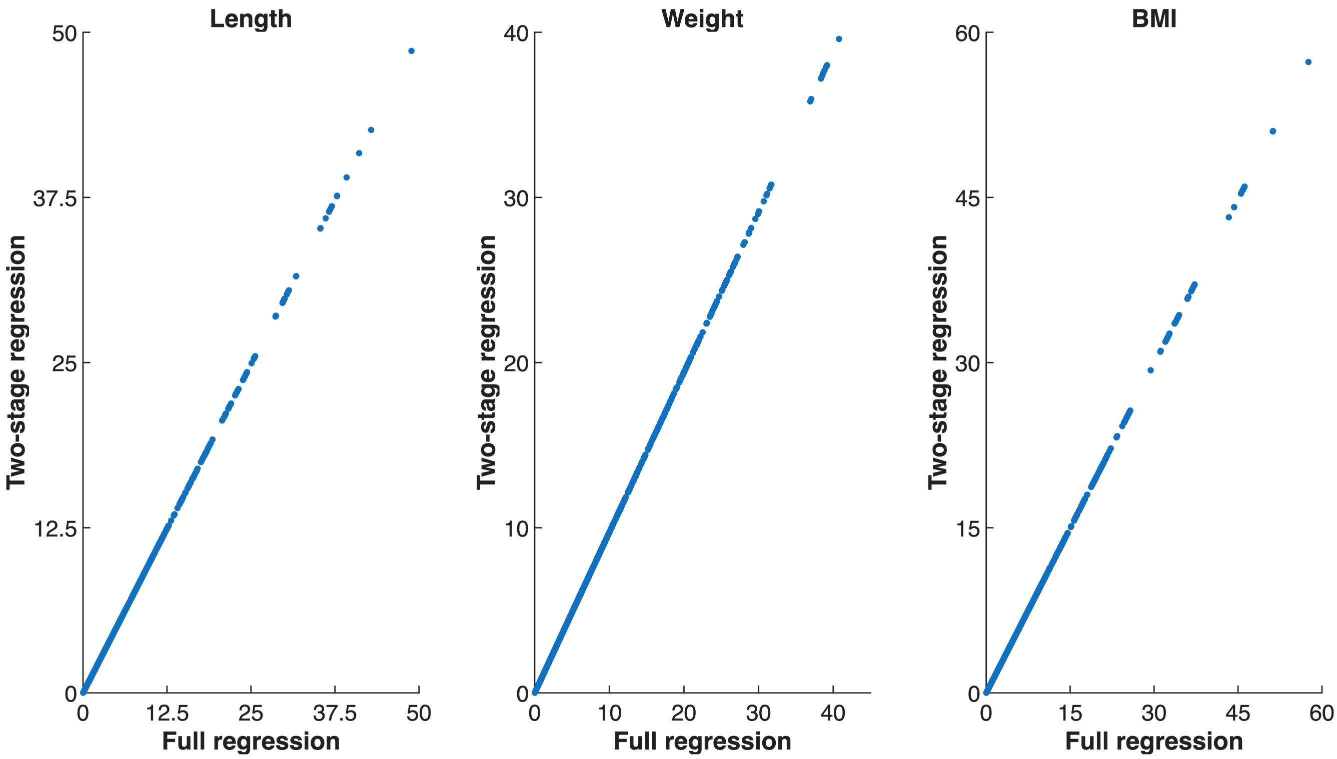


**Figure S49:** Comparison of $-\log_{10} p$ values (main effect and the interaction effects of SNPs with smooth basis functions of age) between full regression (where all terms were estimated together) vs. two-stage regression (where the stage-1 regression and the estimation of random effects covariance terms did not include genetics).

#### Comparison of Wald *F*

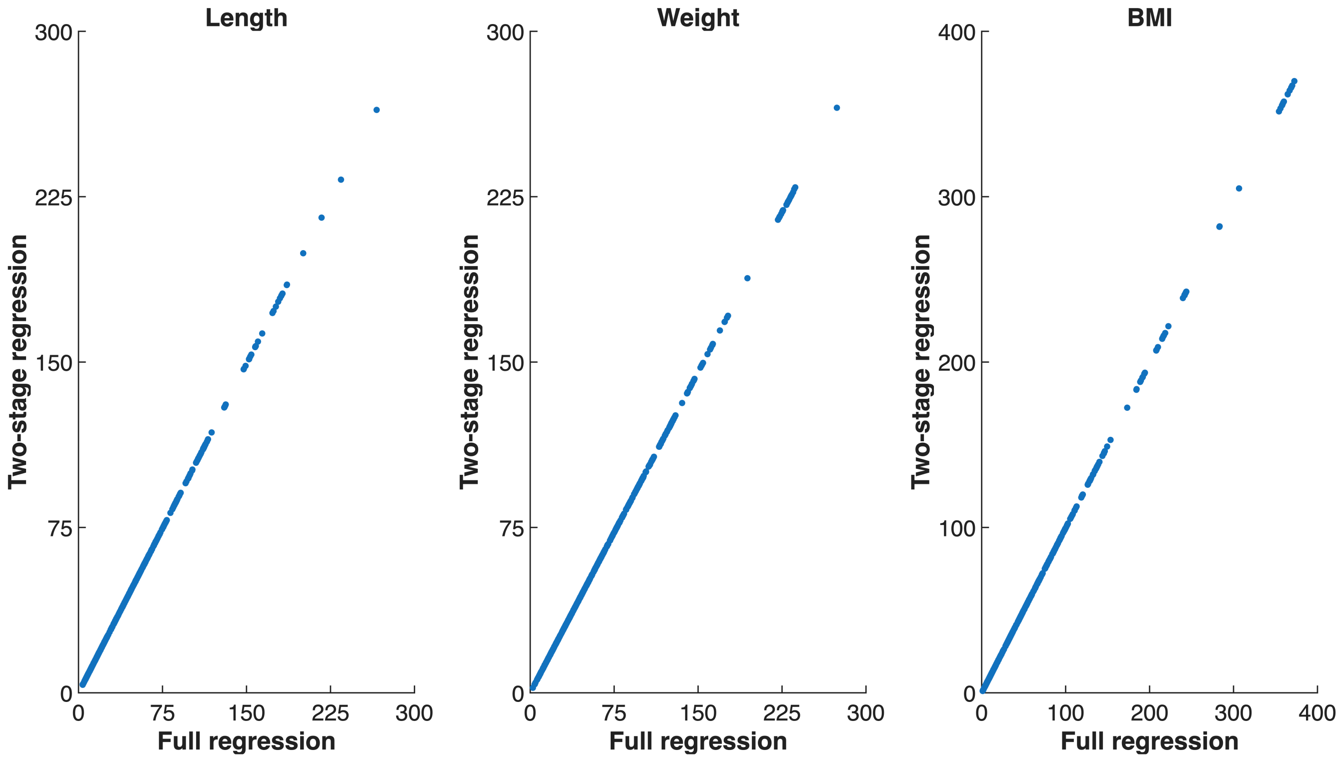


**Figure S50:** Comparison of Wald *F* statistics (across the main effect and the interaction effects of SNPs with smooth basis functions of age) between full regression (where all terms were estimated together) vs. two-stage regression (where the stage-1 regression and the estimation of random effects covariance terms did not include genetics).

#### Comparison of $-\log_{10}(\text{Wald } p)$ values

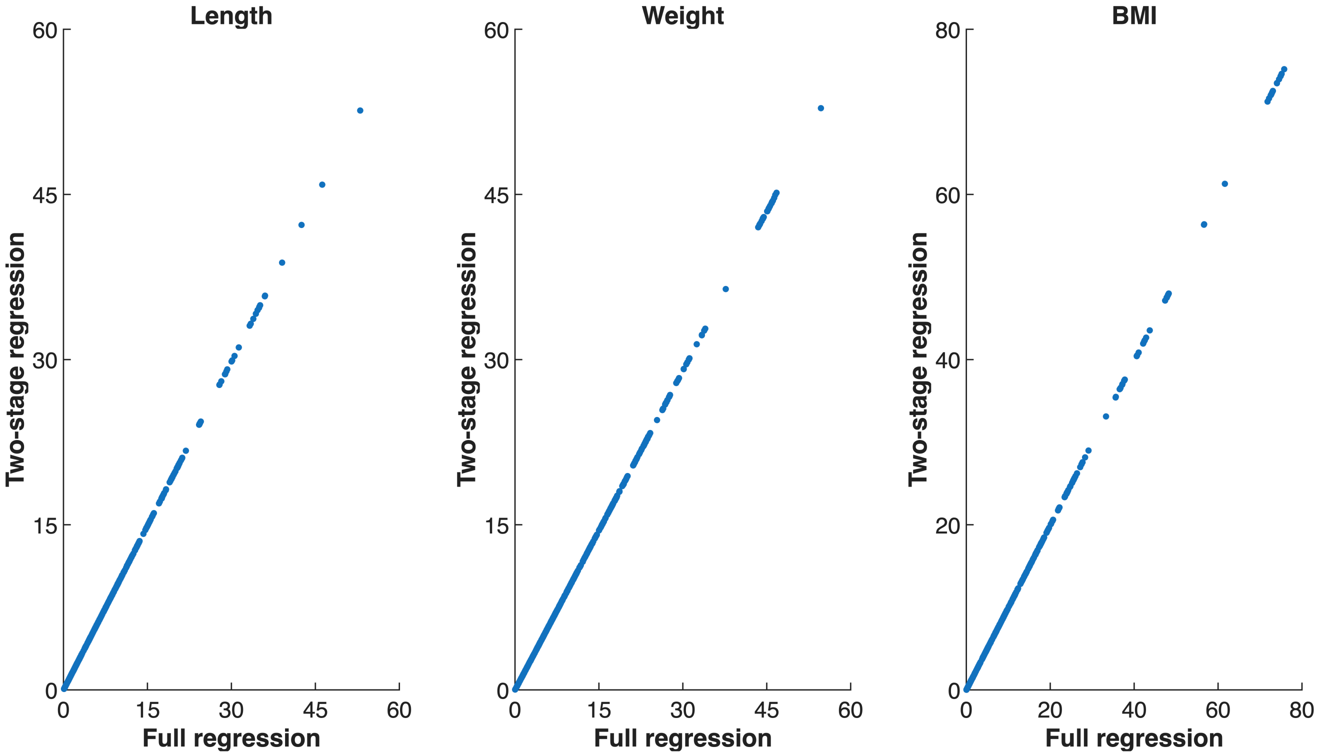


**Figure S51:** Comparison of $-\log_{10}$ Wald $p$ (across the main effect and the interaction effects of SNPs with smooth basis functions of age) between full regression (where all terms were estimated together) vs. two-stage regression (where the stage-1 regression and the estimation of random effects covariance terms did not include genetics).
